# Differentiable Spin-Center Diffusion Model Enables DEER-Guided Generative Pipeline of Weighted Protein Ensembles

**DOI:** 10.64898/2026.10.29.762916

**Authors:** Te-Yu Kao, Jens Meiler, Hassane S. Mchaourab

## Abstract

Biological function entails the interconversion of a protein between distinct conformations, an equilibrium modulated by ligands and other physicochemical conditions. Double electron-electron resonance (DEER) spectroscopy captures these ensembles as distance distributions, P(r), between spin label pairs. However, the width of P(r) convolves protein conformational heterogeneity with the rotameric flexibility of the label. Because current spin label models overestimate this contribution, translating P(r) into ensembles has remained indirect and predominantly qualitative. Here, we developed spin-center (SC) diffusion, a differentiable model of the spin label with a tunable spin label rotameric width. Coupled to a frozen Boltz-2 structure generator, SC diffusion enables the entire experimental P(r) to guide ensemble generation at inference time. We demonstrate the robustness of this pipeline on synthetic P(r) of known composition, recovering the ensemble populations, including both components of a bimodal profile. Guided by experimental P(r) measured under different biochemical conditions, ensembles of the transporters PfMATE, LmrP, MhsT and P-glycoprotein were modeled and free energy differences derived from the ensembles in a condition-dependent manner. By embedding a differentiable label model in structure generation, our approach interprets DEER distributions as weighted conformational ensembles, adding a missing thermodynamic grounding to generative protein structure prediction methods.

## Introduction

Double electron–electron resonance (DEER) spectroscopy measures the distance distribution, P(r), between two spin labels attached to a protein^1,2^. Because every molecule in the sample contributes, P(r) reports on the conformational equilibrium of the protein rather than on a single structure, a property that has been harnessed extensively to characterize such equilibria in membrane proteins^3–8^. The measurement is carried out on samples flash-frozen from ambient temperature, a step that traps the conformations interconverting in solution. The conformational dynamics of the protein are thereby converted into a heterogeneous ensemble of static conformations across the sample. The width of P(r), however, convolves this conformational heterogeneity with the rotameric flexibility of the spin label side chain. Translating DEER data into conformational ensembles therefore requires disentangling the two contributions.

The current practice simulates the label contribution from a single structure. Rotamer library and accessible-volume models such as MMM^9,10^ or MTSSLwizard^11^ position label conformers on a fixed backbone, from which P(r) is predicted. Because these models tend to overestimate the flexibility of the label^12^, they obscure the contribution of conformational heterogeneity in the interpretation of experimental P(r). An alternative to deconvolution of the two contributions is to fit P(r) directly with an ensemble of structures. Restrained-ensemble molecular dynamics (reMD)^13,14^ and ensemble-biased metadynamics (EBMetaD)^15,16^ fit many structures by biasing molecular dynamics simulations toward the target P(r). Other methods select or reweight a pool of candidate conformations^17–19^ derived from molecular dynamics, from AlphaFold2 predictions^20^ diversified by subsampling or clustering of the multiple sequence alignment (MSA)^21,22^, or from DEERFold, which fine-tunes AlphaFold2 with DEER P(r)^23^. Reweighting, however, is limited to the conformations present in the pool. This limitation can be circumvented by deep generative structure models, which sample conformations without recourse to a fixed pool. AlphaFold3, Boltz-1 and Boltz-2^24–26^ generate atomic coordinates through a diffusion module^27–29^ conditioned on a Pairformer trunk, while related generators emulate equilibrium ensembles^30^. Such generators have been guided through their noisy coordinates with crystallographic, NMR or cryo-EM data^31^, as well as with user-specified distance restraints^32^, whereas a frozen network can also be guided by backpropagating a loss into its inputs or internal representations^33,34^.

Guiding a structure generator with DEER data, however, imposes two requirements that existing spin label models do not satisfy. The first is a differentiable translation from protein coordinates to the P(r) of the spin label, in which the rotameric contribution to the width does not overwhelm the contribution that reflects conformational heterogeneity. MMM and MTSSLwizard predict distribution widths from static backbones that exceed the experimental ones for most pairs. Moreover, because they select or weight discrete conformers by steric clashes^9,11^, their predictions are not differentiable with respect to the structure^35^. The second is an objective defined on distances pooled over many structures. Because P(r) is a population-weighted average over the ensemble, no single model is expected to reproduce it. The loss must therefore compare the P(r) pooled over all members of the ensemble with the experimental P(r) to account for the distinct components of a multimodal distribution. DEERFold, the closest predecessor pipeline for DEER-retrained structure prediction, meets neither requirement in full. Its input is the component of P(r) that corresponds to a single state, whereby the restraints act on each structure individually. Its fine-tuning also relied on the distribution widths predicted by the chiLife implementation^36^ of the rotamer library model MMM.

Here, we developed spin-center (SC) diffusion, a differentiable model that predicts nitroxide distributions from the local environment of a labeled site, with a rotameric width tunable to fall below experimental widths. We then integrated SC diffusion with a frozen Boltz-2 network into a pipeline that generates DEER-guided ensembles whose pooled P(r) match the entire target P(r), without molecular dynamics simulations or a precomputed pool. We used this pipeline to (i) recover known populations from synthetic conformational distributions of the transporter PfMATE, (ii) generate condition-specific ensembles for four membrane transporters, and (iii) derive free energy differences between conformations of these transporters. The ensembles shifted predominantly toward the corresponding reference structures, while the sign of the change in the free energy differences reflects the experimental equilibrium shift between conformations under these conditions. Using Boltz-2 as a structure generator allows DEER data to be combined with other information, such as cryo-EM or crystal structures, in integrative studies^19^, whereas ligands permit protein–ligand interfaces to be examined across the ensemble. By coupling a differentiable spin label model to a frozen structure generator, our approach enables DEER P(r) to guide structure generation, thereby transforming DEER data into weighted conformational ensembles.

## Results

To apply the pipeline to experimental data, we selected four membrane transporters with published DEER P(r) measured under different biochemical conditions, each with reference structures or models thereof. PfMATE^37^ is an archaeal H^+^-driven multidrug and toxic compound extrusion transporter. We used published DEER P(r) of 42 pairs measured at pH 7.5 versus pH 4 in nanodiscs^38^, with the outward-facing (OF, PDB code: 6GWH) and inward-facing (IF, PDB code: 6FHZ) crystal structures^39^ as references. LmrP is a bacterial proton-coupled multidrug antiporter of the major facilitator superfamily^5^. Its nine DEER pairs, measured at pH 8 versus pH 5^5^, were used as presented in the study that predicted its IF conformation with AlphaFold2^40^. The references are the OF structure (PDB code: 6T1Z)^41^ and an IF prediction from CASP14 target T1024^40,42^. MhsT^43^ is a bacterial member of the neurotransmitter:sodium symporter family. It was characterized with 22 pairs in detergent micelles, 20 of which were also measured in nanodiscs, each in the apo state, with Na^+^, or with Na^+^ and tryptophan (Trp)^44^. Its references are candidate models derived from AlphaFold2-ensemble clusters reported in that study. P-glycoprotein (Pgp) is a mammalian ATP-binding cassette (ABC) exporter of xenobiotics^6^. For Pgp, 28 pairs measured in the apo versus ADP-vanadate (ADP-Vi) conditions derive from two studies^6,45^, with a human IF structure (PDB code: 7A65)^46^ and a mouse OF structure bound to ATP and Mg^2+^ (PDB code: 7ZK4)^47^ as references. Templates, then ligands, were evaluated as additional Boltz-2 inputs.

### A differentiable spin label model with tunable rotameric width

To determine whether existing label models accommodate conformational heterogeneity, we benchmarked the chiLife implementations^36^ of MMM^9^ and MTSSLwizard^11^ against DEER P(r)^4,14,38^ for 141 pairs on five structures: T4 lysozyme (T4L)^48^, PfMATE in the OF or IF state^37,39^ and the ABC exporter MsbA in the nucleotide-bound or apo state^49,50^. Although both methods predict mean distances that correlate strongly with experiment, their predicted widths exceed the experimental widths for 68% (MMM) and 74% (MTSSLwizard) of pairs (Fig. 1A). Thus, these rotamer models are inadequate to deconvolve the spin label rotamer contribution from that of conformational heterogeneity. To model P(r) with a tunable width as well as a differentiable function, we capitalized on a diffusion model, whose width of P(r) predictions can be modulated by varying the input noise^28,51–54^. We trained SC diffusion, a conditional diffusion model^27,55^ that predicts nitroxide distributions from a PointNet encoding^56^ of the heavy atoms within 12 Å of the labeled C_α_ (Fig. 1B). Training relied on MTSSLwizard predictions at 823,735 solvent-exposed helix and strand sites in 17,592 protein chains from the OpenFold training set^57,58^ (Supplementary Fig. 1, Methods).

**Figure 1.**
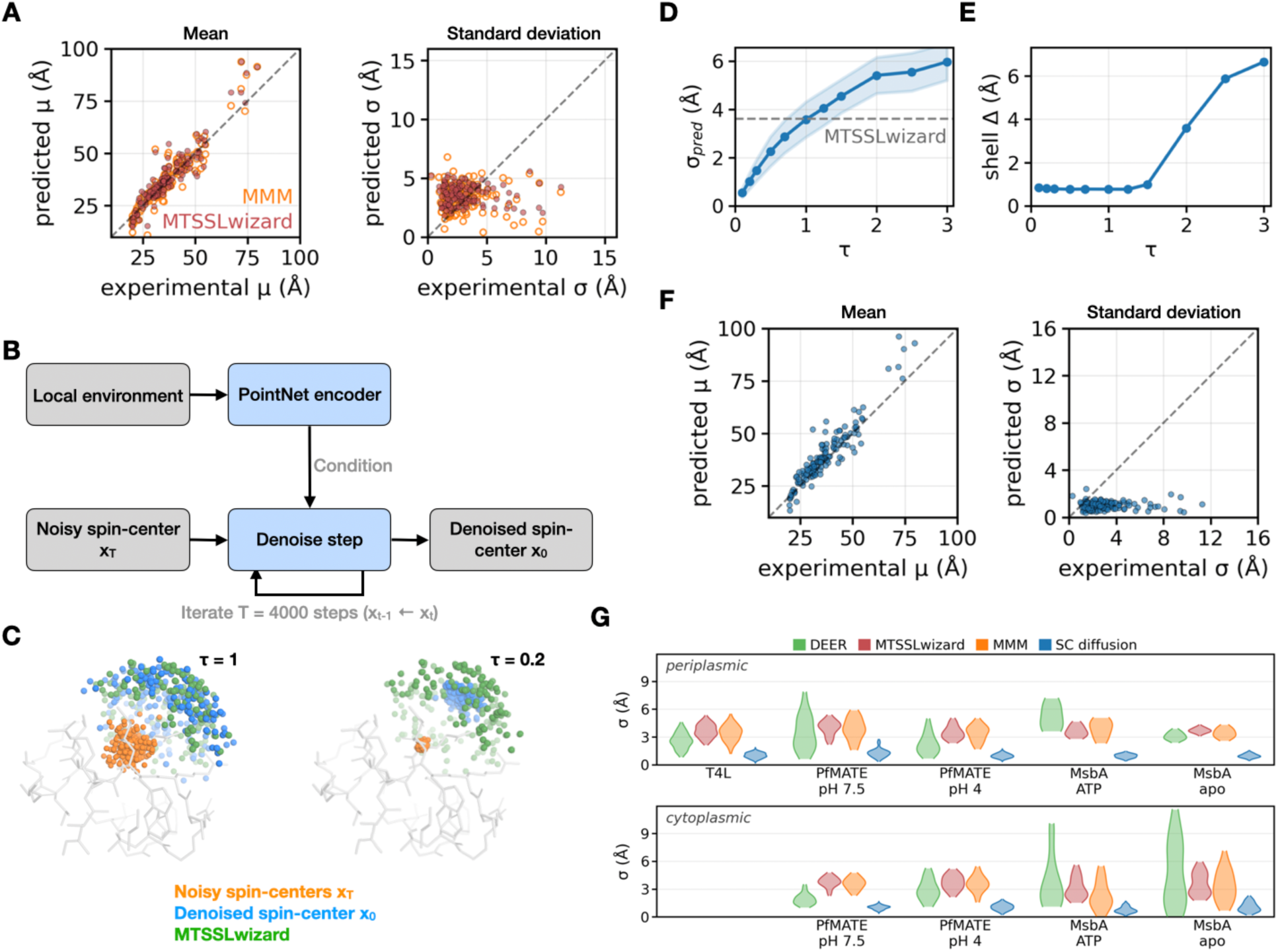
Spin-center (SC) diffusion with tunable temperature. **A**, Predicted versus experimental^4,14,38^ mean μ and s.d. σ for 141 pairs by MMM (orange) and MTSSLwizard (red). Dashed, y = x. PfMATE and MsbA experimental σ are single-Gaussian major-peak widths. Pearson r (MMM/MTSSLwizard), 0.929/0.930 for μ. **B**, Schematic of SC diffusion. **C**, T4L site 128 at τ = 1 and 0.2. Orange, starting noise x_T_. Blue, denoised spin centers x_0_. Green, MTSSLwizard. SC diffusion-MTSSLwizard centroid distance at τ = 0.2, 1.46 Å. **D**, Mean SC diffusion σ_pred_ over the 141 pairs versus τ (±1 s.d. band). Dashed, MTSSLwizard mean, 3.608 Å (SC diffusion, 3.571 Å at τ = 1). **E**, Distance between the SC diffusion and MTSSLwizard shell centers (Methods), 0.76–0.84 Å for τ ≤ 1.25. **F**, As A, for SC diffusion at τ = 0.2 (Pearson r, 0.923 for μ). Mean absolute error of μ, 4.36 Å (MTSSLwizard 3.56 Å, MMM 3.47 Å). **G**, Per-pair σ as violin plots (SC diffusion at τ = 0.2), with water-soluble T4L in the periplasmic row for layout. σ_pred_ ≥ σ_exp_ in 7 (SC diffusion), 104 (MTSSLwizard) and 96 (MMM) of 141 pairs.

Sampling is initiated from nitroxide positions drawn from a three-dimensional Gaussian distribution centered on the labeled C_α_, whose standard deviation, in Å, is set by the temperature τ (Methods). For example, at T4L site 128, lowering τ from 1 to 0.2 narrows the predicted nitroxide distribution from a root-mean-square spread of 5.36 Å, close to the 5.53 Å predicted by MTSSLwizard, to 1.50 Å (Fig. 1C). Across all 141 pairs, the average width increases monotonically with τ, intersecting the MTSSLwizard average at τ = 1 (Fig. 1D). The temperature also displaces the shell center (see Methods), by a median of 0.46 Å between τ = 0.1 and 1, whereas its offset from the MTSSLwizard shell center remains nearly constant up to τ = 1.25 (Fig. 1E). At τ = 0.2, the predicted mean distances obtained from the P(r) remain comparably correlated with experiment (Pearson r = 0.923, compared with 0.929 for MMM and 0.930 for MTSSLwizard), with a mean absolute error of 4.36 Å, compared with 3.47 Å and 3.56 Å, respectively (Fig. 1F). The widths, in contrast, are narrowed, falling below the experimental widths for 134 of 141 pairs.

The temperature τ is a design choice rather than an intrinsic property of the spin label. Because nearly all measured widths exceed 1 Å (138 of 141 pairs), we selected τ = 0.2, which yields a predicted width of approximately 1 Å (Supplementary Fig. 2). Across soluble and membrane protein sites, SC diffusion is the only method whose mean width falls below the DEER mean in all nine groups (Fig. 1G, Supplementary Fig. 3). If the rotameric ensembles of the spin label and the conformational heterogeneity contribute independently to the DEER widths, i.e., their variances are additive, then 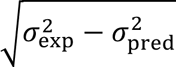 represents the width attributable to conformational heterogeneity.

### DEER-guided ensemble generation by combining SC diffusion with a structure generator

To translate P(r) into ensembles, we integrated SC diffusion with Boltz-2^26^ into a pipeline that generates an ensemble yielding a pooled P(r) that matches the target P(r) (Fig. 2A, Methods). Each model in the ensemble receives its own copy of the single and pair representations that enter the Boltz-2 Pairformer. In de novo runs, Boltz-2 derives the starting representations from the sequence and MSA alone, which renders all models initially similar. For each of the predicted models, SC diffusion predicts nitroxide distributions at every label site. The P(r) of all models are pooled into a single predicted P(r) per pair, summarized by its 48 quantiles, the distances below which fractions 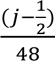 of the distribution lie (j = 1,…,48). Because the target quantiles constitute an equivalent representation of the target P(r), their deviation from the predicted quantiles serves as the principal driving loss, whose minimization yields ensembles that match the target P(r).

**Figure 2.**
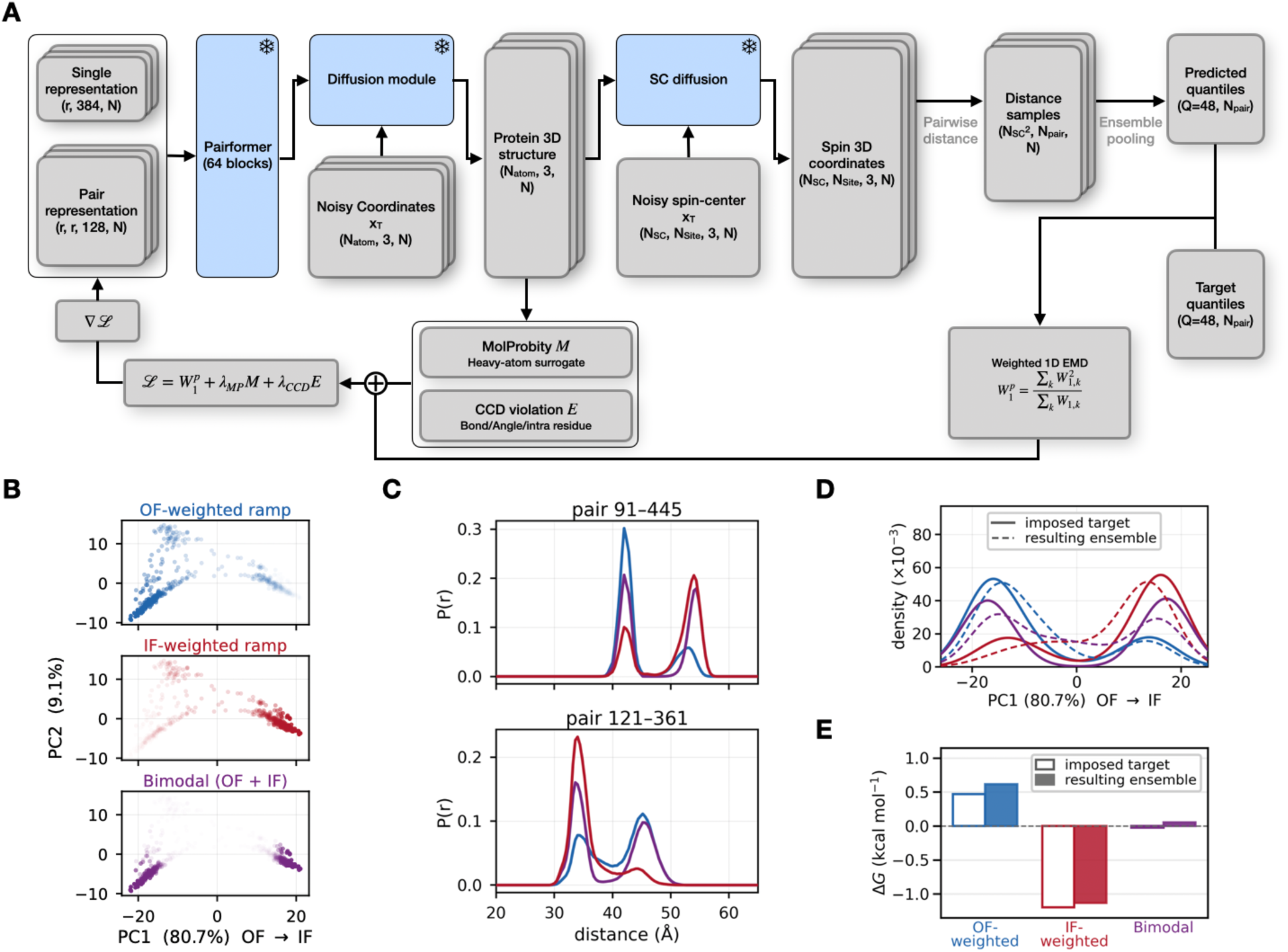
DEER guidance scheme and synthetic-target tests. **A**, Snowflakes mark frozen Boltz-2^26^ and spin-center (SC) diffusion weights. Gradients ∇ℒ are backpropagated into per-model copies of the single and pair representations. *W*_1,*k*_, Wasserstein-1 distance of pair k. Each λ is set per guided step by a gradient-norm budget (Methods). **B**, Pool of 500 PfMATE DEERFold^23^ models projected onto the first two principal components, with PC1 from outward-facing (OF) to inward-facing (IF). Opacity increases with the imposed population. Colors (subpanel titles) apply to B–E. **C**, Imposed target P(r) for two of the 40 fitted pairs. **D**, PC1 kernel densities of imposed targets and resulting ensembles (N = 96 models). **E**, Free energy difference Δ*G* = *G*_IF_ − *G*_OF_at an assumed 298 K, read from landscapes on the gate distances at fixed OF and IF positions (Methods).

The DEER loss, a weighted earth mover’s distance (EMD)^59^ between these quantiles, is supplemented by two terms that penalize structural distortions, a differentiable heavy atom surrogate of the MolProbity score^60^ and a penalty on Chemical Component Dictionary (CCD)^61^ bound violations. In one dimension, the EMD between two P(r) equals their Wasserstein-1 distance, the mean absolute difference between their quantiles, which we denote *W*_1,*k*_ for pair k. The weighted EMD, 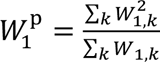, weights each pair by its own *W*_1,*k*_, such that poorly fitted pairs contribute stronger gradients. Because every step of the pipeline is differentiable, this loss was minimized with the Adam optimizer^62^, backpropagating its gradient through the SC diffusion model, the Boltz-2 diffusion module and the Pairformer into the representations (Methods). Guidance commenced at the highest-noise diffusion step (Supplementary Fig. 4). Because each model receives its own gradients and diffusion noise, the models diverge from one another, their structures evolving such that the pooled P(r) converges toward the target P(r). After optimization, every structure was relaxed by restrained energy minimization as in AlphaFold2^20^. Model weights *θ*^∗^, summing to one, were then fitted to minimize the average over pairs of the EMD between the weighted P(r) and the target P(r). We report this average at *θ*^∗^ as 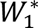.

### Pipeline validation with synthetic ensembles

To validate the pipeline on targets with known ground truth, we constructed synthetic targets from the pool of 500 DEERFold^23^ models of PfMATE generated by simulated DEER distance restraints (Fig. 2B–D, Methods). The models of the pool were ranked from OF to IF by the difference between their root-mean-square deviations (RMSD) from the OF and IF crystal structures, RMSD_OF_ − RMSD_IF_(Supplementary Fig. 5A). Three population profiles were constructed over the pool: one weighted toward OF, one toward IF, and a bimodal profile with two equally populated regions (Fig. 2B, Supplementary Fig. 5B). For the 40 pairs most sensitive to the OF-to-IF transition, each target P(r) is the sum of the SC diffusion predictions for the pool, weighted by the population profile. The three profiles alter the relative populations of peaks at nearly fixed positions (Fig. 2C). To evaluate the resulting ensembles, each was projected onto the first principal component (PC1) of the pool’s C_α_ coordinates, which spans the OF-to-IF transition (Supplementary Fig. 5C). Because PC1 is not part of the loss, agreement along this coordinate is not imposed by the fit. These targets represent ideal cases, computed with the same SC diffusion model, free of experimental noise and analytical error.

Runs against the three targets (Supplementary Table 1) yielded ensembles with the dominant population at the correct end for the OF- and IF-weighted profiles, with both populations recovered for the bimodal target (Fig. 2D). However, on each side of PC1, the ensembles were generally broader than the targets, overpopulating the center of the axis. Each ensemble resided closest to its own target in the PC1–PC2 plane (Supplementary Fig. 6), matching its target P(r) with 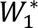 of 0.28 to 0.35 Å (Supplementary Fig. 7). Three replicates of the bimodal run, differing only in the Boltz-2 and SC diffusion seeds, yielded similar PC1 densities and 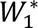 values (Supplementary Fig. 8). On these ideal targets, we also computed the free energy difference between OF and IF from the free energy landscape on the two gate distances, read at fixed OF and IF positions (see below, Methods). At an assumed 298 K, the errors were 0.14, 0.07 and 0.08 kcal mol^−1^ for the OF-weighted, IF-weighted and bimodal targets, respectively (Fig. 2E).

The selection of pairs proved more consequential than their number (Supplementary Fig. 9), reflecting the importance of the information content. With the k pairs that most discriminate between the two conformations, the PC1 density became bimodal from k = 3 onward, whereas the 40 pairs with the smallest OF–IF distance difference collapsed the ensemble onto a single OF-side mode. Because τ is a design choice, the pipeline was also executed at τ = 0.1, 0.2, 0.5 and 1.0. Varying τ had little effect on the bimodal feature, although τ = 1.0 overpopulated the center of PC1 beyond the range observed at τ = 0.1 and 0.2 (Supplementary Fig. 10).

### DEER-guided ensembles track condition-dependent conformational shifts

To test whether the pipeline resolves conformational changes induced by shifts in biochemical conditions, we performed one de novo run per condition against the experimental DEER P(r) (Supplementary Table 1). Given that the experimental structures used as references in Fig. 3 were released before the Boltz-2 training cutoff^26^, model similarity to these references may reflect recall by Boltz-2. Recall alone, however, cannot account for differences between ensembles of the same protein, since their runs shared the default generator input, differing only in the target P(r).

**Figure 3.**
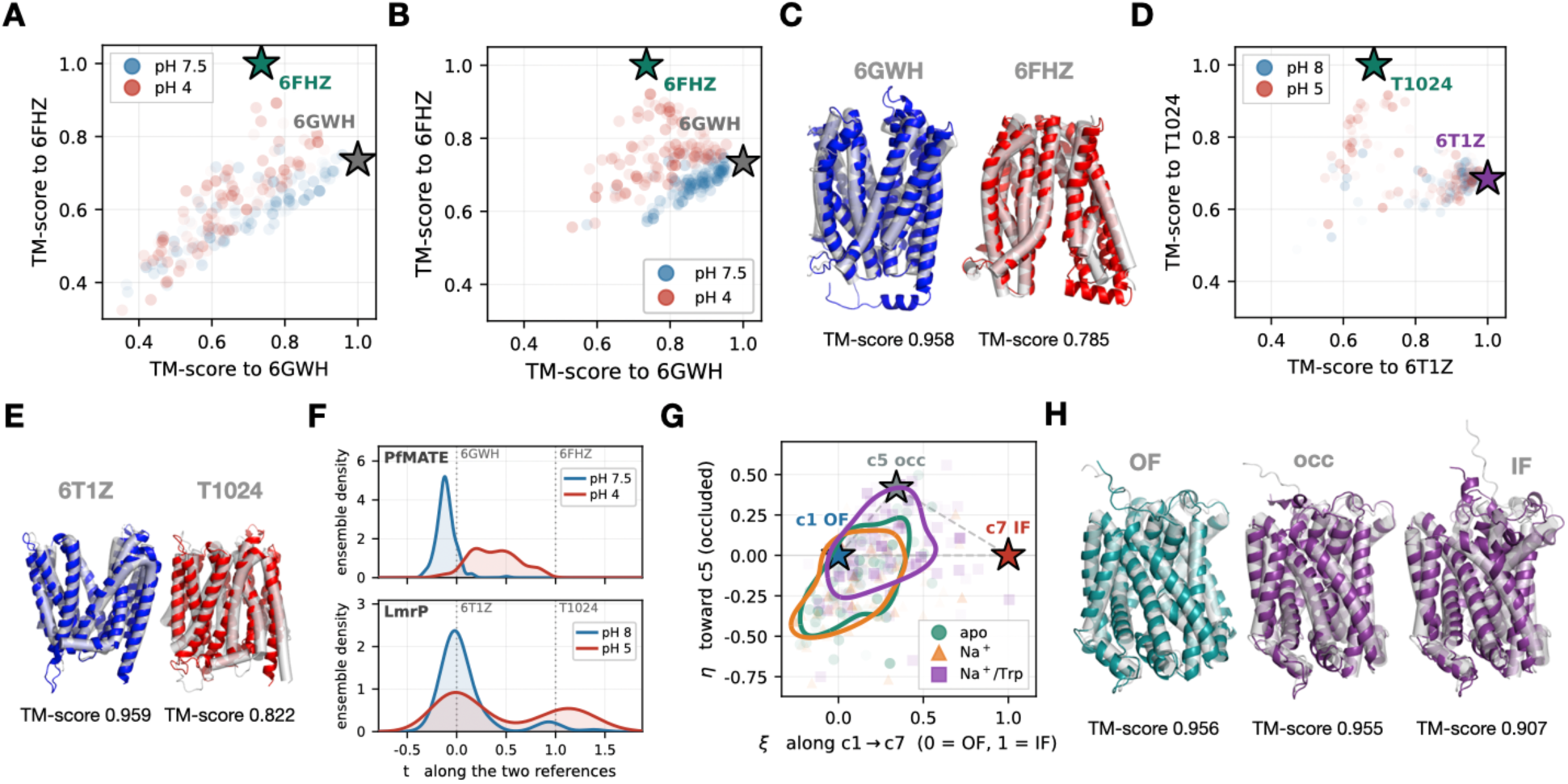
Ensembles fitted to DEER data from different conditions. **A**, TM-scores of the models of PfMATE ensembles^38^ (N = 96 per run, opacity scaled by *θ*^∗^) to 6GWH (outward-facing, OF) and 6FHZ (inward-facing, IF). Stars mark the reference structures (also in B and D). **B**, As A, for tail-truncated targets. **C**, Model of each run in A, on the gray reference, with the largest mean inner product between predicted and target P(r) among models with nonzero *θ*^∗^ (Methods). **D**, As A, for LmrP^5,40^, versus 6T1Z and the CASP14 prediction T1024. **E**, Model of each LmrP run in D, selected and shown as in C. **F**, *θ*^∗^-weighted densities of t, the projection on the axis between the two references (dotted), for B (top) and D (bottom), with separate y scales. **G**, MhsT in detergent micelles^44^, with stars for the candidate cluster models c1 (OF), c5 (occluded, occ) and c7 (IF). Contours, 88th percentile of each *θ*^∗^-weighted density. **H**, Models with nonzero *θ*^∗^ closest by C_α_ RMSD to c1 (apo run of G) and to c5 and c7 (Na^+^/Trp run), on gray references.

For PfMATE (Supplementary Fig. 11), Fig. 3A presents the template-modeling scores (TM-scores)^63^ of each model to 6GWH (OF) and 6FHZ (IF). The pH 4 ensemble is shifted toward IF relative to pH 7.5, albeit partially, as both remain closer to OF (*θ*^∗^-weighted TM-scores to 6GWH and 6FHZ: 0.705 versus 0.592 at pH 7.5, 0.648 versus 0.621 at pH 4). Distorted structures (minimum TM-score below 0.55) are associated with the long tails of the experimental P(r). Typically, such long tails may reflect protein aggregation that varies between mutants. When ensembles were guided toward targets with the long tails truncated (tail-truncated targets, Supplementary Fig. 12), distorted structures were nearly eliminated, from 69 to 1 of 192 (Fig. 3B), while the pH 4 ensemble retained its shift toward IF. Figure 3C depicts the model of each run in Fig. 3A whose prediction best matches the target P(r), selected independently of the reference structures (Methods).

For LmrP (Supplementary Fig. 13), the pH 8 ensemble is localized near the OF structure 6T1Z (Fig. 3D). The pH 5 ensemble, by contrast, is more dispersed, shifting toward T1024 with 42.1% of its *θ*^∗^ mass closer to this reference, compared with 8.8% at pH 8. Figure 3E depicts the models selected by the same criterion. PfMATE and LmrP belong to different families, yet both consist of two six-helix lobes that move relative to each other in the OF-to-IF transition^5,38,39,41^, with low pH favoring IF in both^5,38,64^. We therefore projected each ensemble of Fig. 3B,D onto the axis connecting its OF to its IF reference. Both low-pH ensembles are shifted toward IF relative to high pH (Fig. 3F).

For MhsT, we fitted the apo, Na^+^ and Na^+^/Trp P(r) measured in detergent (DM), with neither ions nor substrate in the generator input (Supplementary Fig. 14). The references are candidate OF (c1), occluded (c5) and IF (c7) models whose states were assigned in the study that reported these data^44^. Agreement with these models therefore constitutes a consistency check rather than independent validation. Figure 3G projects the predicted models on two axes, ξ from OF to IF, η toward occluded. The apo and Na^+^ ensembles overlap near OF, whereas the Na^+^/Trp ensemble is shifted toward IF, with a concomitant shift toward occluded. Figure 3H depicts the models closest to c1, c5 and c7, selected by structural similarity rather than by the DEER data. The nanodisc (ND) data, measured on a subset of the DM pairs, yield the same pattern (Supplementary Figs. 15 and 16).

### DEER-guided ensembles of Pgp reveal previously unobserved conformations

The data set for Pgp consists of pairs that monitor the global IF-to-OF transitions and pairs that monitor the local structures near the nucleotide binding sites (NBSs), the regions of the nucleotide binding domains (NBDs) where ATP is bound and hydrolyzed. The latter pairs were designed to test the hypothesis that Pgp ATP hydrolysis is asymmetric, i.e., the two NBSs are not catalytically equivalent despite their conserved sequence motifs. The IF-to-OF transition is represented in Fig. 4 with one pair each in the NBDs, the intracellular transmembrane domain (TMD) and the extracellular TMD. In Fig. 4, the run guided by the ADP-Vi P(r) included two ATP and two Mg^2+^ in the Boltz-2 input (ATP/Mg^2+^ run). The ATP/Mg^2+^ ensemble shifts toward the OF structure 7ZK4, with 41.6% of its *θ*^∗^ mass closer to 7ZK4 by TM-score, compared with 8.9% in the apo state (Fig. 4A and Supplementary Fig. 17). In the ADP-Vi conditions, the P(r) of the extracellular pair shifts to longer distances, whereas those of the intracellular and NBD pairs shift to shorter distances, changes that the ensembles mirror (Fig. 4B,C).

**Figure 4.**
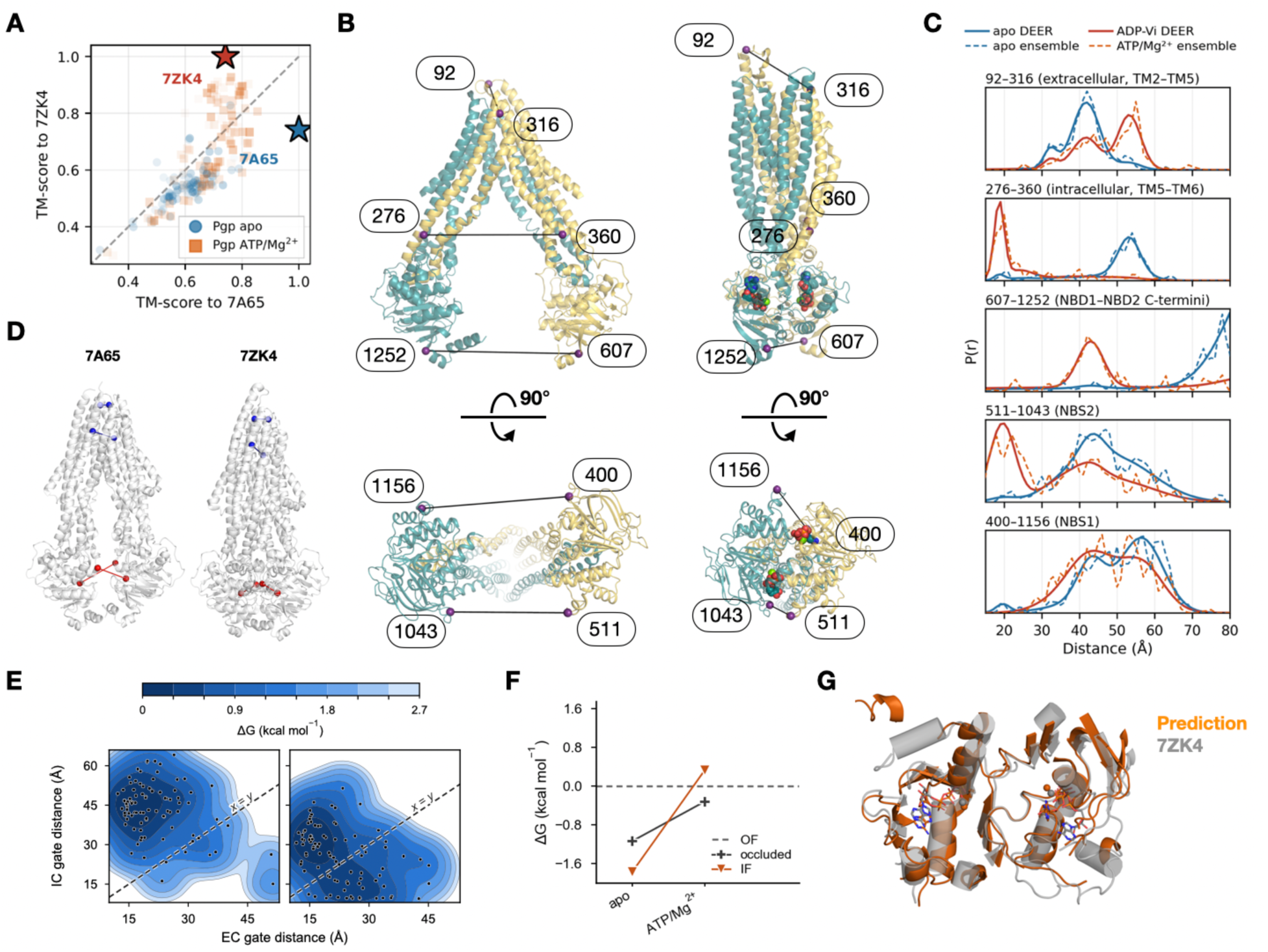
DEER-guided ensembles of Pgp in the apo and ADP-vanadate conditions. **A**, TM-scores of the models of the apo run and of the ADP-vanadate (ADP-Vi) run with two ATP and two Mg^2+^ in the input (ATP/Mg^2+^)^6,45^ (N = 96 per run, opacity scaled by *θ*^∗^) to 7A65 (inward-facing, IF) and 7ZK4 (outward-facing, OF). Stars mark the reference structures. Dashed, equal TM-score. **B**, Model of the apo run (left) and of the ATP/Mg^2+^ run (right, ATP and Mg^2+^ as spheres), selected as in Fig. 3C by the largest mean inner product between predicted and target P(r), in two views related by a 90° rotation. Spheres mark the labeled residues of C, and lines join their C_α_ atoms. **C**, DEER P(r) (solid) and *θ*^∗^-weighted P(r) of the resulting ensembles (dashed) at five of the 28 guided pairs, apo in blue and the ATP/Mg^2+^ ensemble in orange, with residue numbers of mouse Pgp. The apo P(r) of 607–1252 accumulates at the 80-Å limit of the measurement window, which reflects the window, not a physical peak. **D**, Extracellular (EC) and intracellular (IC) gate pairs on 7A65 (IF, left) and 7ZK4 (OF, right), extracellular side up. The gate distances *d*_EC_ and *d_IC_* are mean C_α_–C_α_ distances over these pairs (Methods). **E**, Landscapes *G* = −*RT* ln *p*, on *d*_EC_ (x) and *d*_IC_ (y) for apo (left) and ATP/Mg^2+^ (right), with p the *θ*^∗^-weighted kernel density of the N = 96 models (Methods), each zeroed at its minimum and drawn up to *RT* ln(*N* − 1) in 0.3 kcal mol^−1^ bands. Dots mark models with *θ*^∗^ > 0. **F**, *G*_state_ − *G*_OF_ read from E at the fixed positions defined in Methods, as values of a smoothed landscape, not populations. Free energies are relative and at an assumed 298 K. **G**, Model 27 of the ATP/Mg^2+^ run (the right model of B, selected by the largest mean inner product, with ATP as sticks and Mg^2+^ as spheres) on 7ZK4 chain A.

As noted previously^6,45^, the NBS pairs (511–1043 and 400–1156) show asymmetric experimental P(r) in the ADP-Vi conditions, with only one pair (511–1043) showing a short component as expected in the catalytic state of the NBS and from the cryo-EM structure. Yet such asymmetric conformations have not been observed experimentally. In contrast, guiding Boltz-2 prediction with these pairs induces the population of previously unobserved conformations that reproduce the local asymmetry captured by the experimental P(r) (Fig. 4C).

### Free energy analysis from DEER-guided ensembles

To compare the conditions quantitatively, we defined for each target collective variables that reflect the alternating access of the transporters. We projected the transporter ensembles onto the plane of two alternating access gate distances, *d*_EC_ and *d*_IC_, the mean C_α_–C_α_ distances of reported residue pairs that change upon opening of the extracellular or intracellular gate, respectively (Fig. 5A). The resulting landscapes change between conditions as expected (Fig. 5B–D). PfMATE and LmrP shift from OF at high pH toward IF at low pH, whereas MhsT narrows its extracellular gate while widening its intracellular gate with Na^+^/Trp. In addition to models near the two end states, every run contains models with intermediate gate distances. The dashed line x = y marks equal gate distances for reference.

**Figure 5.**
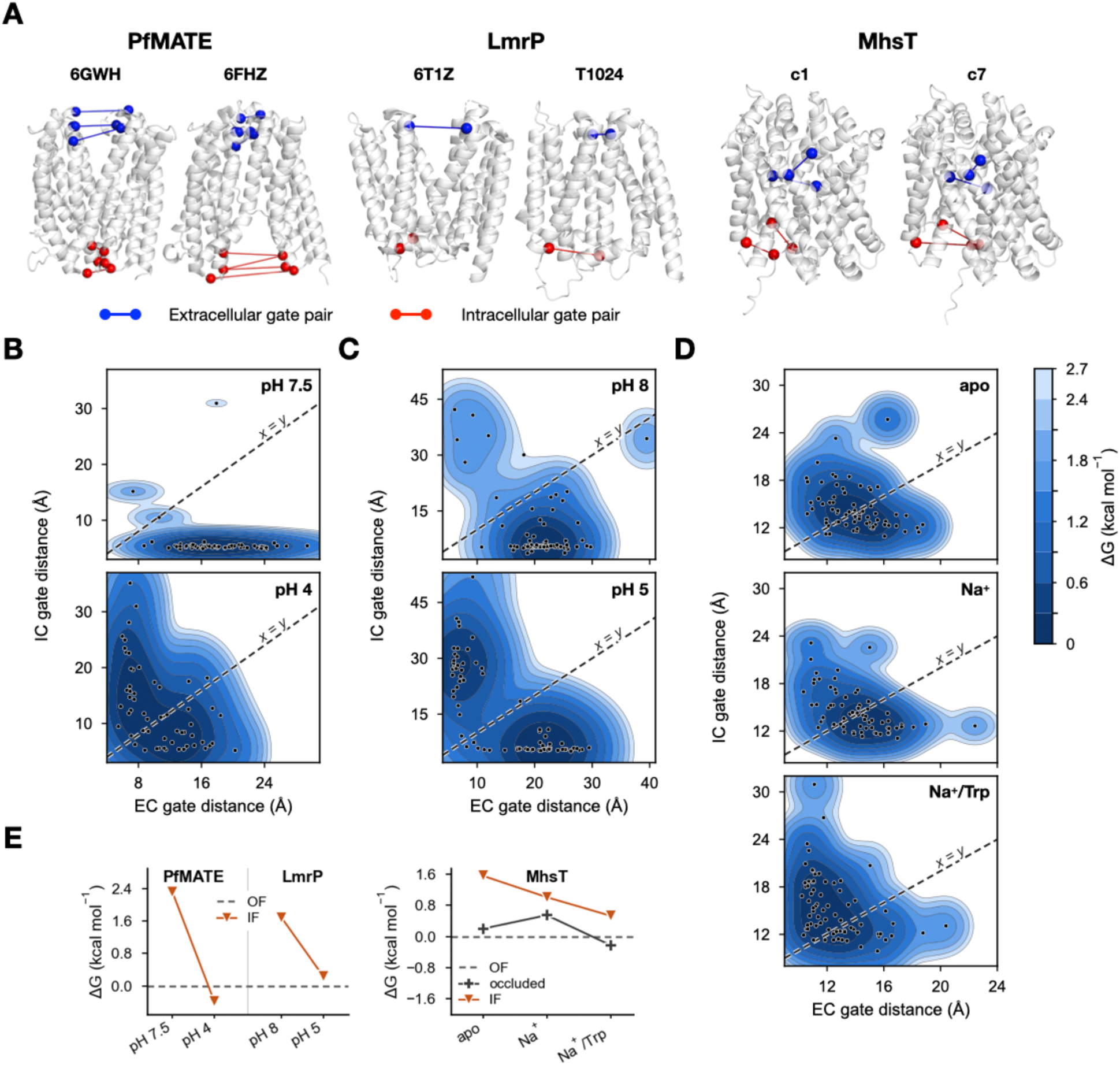
Free energies between conformational states. Extracellular (EC) and intracellular (IC) gate distances d_EC_ and d_IC_ are mean C_α_–C_α_ distances over gate pairs defined in previous work (Methods). Free energies are relative and at an assumed 298 K. **A**, Gate pairs on one outward-facing (OF, left) and one inward-facing (IF, right) reference per protein, extracellular side up. T1024 is a CASP14 prediction, and c1 and c7 are MhsT candidate cluster models. **B–D**, Landscapes *G* = −*RT* ln *p* on d_EC_ (x) and d_IC_ (y), with p the *θ*^∗^-weighted kernel density of the N = 96 models (Methods). Each landscape is zeroed at its minimum and drawn up to *RT* ln(*N* − 1) in 0.3 kcal mol^−1^ bands. Dots mark models with *θ*^∗^ > 0. **B**, PfMATE runs fitted to the tail-truncated targets (Fig. 3B). **C**, LmrP. **D**, MhsT in detergent micelles. **E**, Free energies relative to OF, with lines joining conditions. Left, PfMATE and LmrP runs in B and C. Right, MhsT. *G*_state_ − *G*_OF_ read from B–D at the fixed positions defined in Methods, as values of a smoothed landscape, not populations.

To place the Pgp transition on the gate coordinates used for the other transporters, we projected the Pgp ensembles onto the same two gate distances, computed from reported Pgp gate pairs (Fig. 4D, Methods). In the ATP/Mg^2+^ run, the NBDs approach each other (Fig. 4E). Figure 4F reports the free energies of the occluded and IF conformations relative to OF, obtained as values of the smoothed landscapes at fixed positions (Methods). Relative to apo, the ATP/Mg^2+^ run raised the occluded and IF values by 0.82 and 2.10 kcal mol^−1^, respectively, favoring OF. The ADP-Vi run without ligands shifted in the same direction, by 1.06 and 2.48 kcal mol^−1^ (Supplementary Table 2). The model of the ATP/Mg^2+^ run selected as in Fig. 3C, by the largest mean inner product between predicted and target P(r), is also the one closest to 7ZK4, with ATP–Mg^2+^ occupying both NBSs (Fig. 4B,G).

As for Pgp, Fig. 5E reports the free energies relative to OF as values of the free energy landscapes at fixed positions, with positive values favoring OF (Methods). For PfMATE and LmrP, these positions are the deepest minima nearer an OF or an IF reference, whereas for MhsT they are the gate coordinates of the candidate models c1 (OF), c5 (occluded) and c7 (IF).

At low pH, the free energy difference changed by −2.70 kcal mol^−1^ for PfMATE in the runs of Fig. 3B and by −1.45 kcal mol^−1^ for LmrP, favoring IF in both. For MhsT, Na^+^ raised the occluded free energy by 0.35 kcal mol^−1^ and lowered the IF value by 0.55 kcal mol^−1^, whereas the addition of Trp lowered both, by 0.77 and 0.48 kcal mol^−1^, respectively, shifting the ensemble from OF toward the occluded and IF conformations. Across all runs of the four proteins (Supplementary Table 2), the free energy differences changed in the expected direction for each of the 12 pairs of conditions for which the direction of the experimental conformational shift is known.

### Templates aid local conformational sampling without changing ensemble fits

To test whether prior structural information can seed an ensemble without restraining it, we harnessed the TemplateModule of Boltz-2, which incorporates template information into the pair representation without restraining coordinates^26^ (Supplementary Fig. 18A). A subset of the models of each ensemble was seeded with template-informed pair representations, using the references of PfMATE, LmrP and MhsT as templates. Each of the ten template runs also contains de novo models, equal in number to each template group, to avoid confining the ensemble to conformations near the templates. The PfMATE template runs and the de novo runs compared with them were fitted to the tail-truncated target P(r) of Fig. 3B.

Within a run, the final conformation of each predicted model reflects its starting template. In PfMATE at pH 4, 6GWH-templated models resemble de novo models, which initiate near 6GWH, whereas 6FHZ-templated models converge closer to 6FHZ than de novo models (Supplementary Fig. 18B). In LmrP at pH 5, T1024-templated models remain near T1024, whereas de novo and 6T1Z-templated models converge near 6T1Z (Supplementary Fig. 18C). In MhsT with the DM Na^+^/Trp target, each template group resides nearest its own template, while de novo models overlap the c1 group (Supplementary Fig. 18D). In template runs for the remaining seven targets, template groups likewise converged closer to their own template than de novo models (Supplementary Fig. 19).

From the initial state before DEER guidance to the relaxed final state, models of the template groups in Supplementary Fig. 18B–D move both closer to and farther from their templates (Supplementary Fig. 18E). Nevertheless, all seven groups terminate closer to their own template than the de novo models of the same run. Thus, the template acts as a starting point for sampling rather than as a restraint. Over all ten runs, the template specificity was positive for all 26 template groups (Methods).

Compared with the same-condition de novo runs, template runs exhibited similar 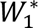. Supplementary Fig. 18F presents the RMSD between their *θ*^∗^ -weighted average structures, a metric that reflects the location of an ensemble. Same-condition differences fall among the smallest values of this metric, whereas cross-condition differences are larger. However, the MhsT apo and Na^+^ runs are not fully resolved (Supplementary Fig. 20). Overall, templates imprint a memory on individual models without a detectable difference at the ensemble level relative to de novo runs.

### Boltz-2 places ligands in DEER-guided ensembles

Finally, to examine whether ligands can be carried through DEER-guided generation, we supplied two Na^+^ ions, alone or with Trp, to the MhsT input of Boltz-2^26^, without positional restraints (Supplementary Fig. 21A). For Pgp, two ATP and two Mg^2+^ were supplied in the same manner (the ATP/Mg^2+^ run of Fig. 4). Relative to the de novo runs, 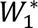 changed by −0.04 to 0.08 Å, indicating that the ligands had no systematic effect on the fit (Supplementary Table 1 and Supplementary Figs. 17, 22 and 23).

The MhsT ligand run targeted to the DM Na^+^/Trp P(r) resides between c1 and c5 on the ξ–η plane, extending toward c7 (Supplementary Fig. 21B). The model in Supplementary Fig. 21C, selected as in Fig. 3C (Methods), positions its Na^+^ ions and Trp within 0.24–0.49 Å of their locations in the 4US3 crystal structure^43^.

In control runs of Boltz-2 inference alone on the same inputs, the MhsT Trp distribution was centered within 0.12 Å of its crystallographic position with little scatter, whereas under guidance it was displaced by 0.59 Å, with concomitant broadening (Supplementary Fig. 21D). Of the 96 unguided Boltz-2 models, 73 positioned the first Na^+^ ion (Na1) within 2 Å of its binding site, compared with 67 of the 96 guided models. Among these, the Na1 offset was identical in both runs (0.34 Å), whereas the spread was wider under guidance (0.52 versus 0.05 Å). Most models of both runs positioned the second Na^+^ ion (Na2) outside its binding site. For Pgp, each of the four ligands resided within 2 Å of its 7ZK4 binding site in all 96 inference samples but in only 40–68 of the 96 guided models (Supplementary Fig. 21E). Ligand placement accuracy therefore originates from Boltz-2 itself, whereas guidance broadens the ligand distributions.

Supplementary Fig. 21F compares ligand versus de novo runs targeted to the same DEER P(r) by the RMSD between their *θ*^∗^-weighted average structures. For both proteins, this RMSD is smaller than any RMSD in the panel between runs of the same protein targeted to different P(r). Thus, inclusion of the ligands perturbed the DEER-guided ensembles less than a change in the DEER target (Supplementary Fig. 21F).

## Discussion

The premise of the work presented here is that DEER P(r) can guide a generative structure model toward the underlying conformational ensemble of a protein, provided that the contribution of the spin label rotamers to the distribution width is accounted for. SC diffusion provides such a spin label model, differentiable with respect to the structure, whereas the frozen Boltz-2 generator supplies the structures. We establish this method through application to four transporters with experimentally reported P(r). The resulting inference-time pipeline enables the translation of the P(r) into condition-specific ensembles that shift in the expected direction, from which free energy differences between conformational states are determined, while accommodating templates or ligands as inputs.

As outlined in the Introduction, existing approaches are bounded by the content of a precomputed pool or, for DEERFold, by a single-state P(r) input. The pool limit is quantifiable. Even optimal weighting of the 500-model PfMATE DEERFold pool yielded a 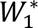 of 6.1–6.6 Å to the full experimental targets, compared with approximately 0.7 Å for de novo guided ensembles fitted to the same pairs. The same dependence applies to the refinement of one structure per state (Rosetta^65^, DEERefiner^66^), to reweighting^67^ and to AlphaFold2 pools generated by MSA perturbation^68–70^. Experimental data can also enter a network through fine-tuning (AlphaLink^71^), per-target weight updates (Distance-AF^72^) or guidance of a frozen network, through its inputs or representations^33,34,73,74^ or the noisy coordinates of each diffusion step^31,32,53,75^. Our pipeline backpropagates the loss into the single and pair representations of each model in a frozen Boltz-2 network (Methods). Unlike DEERFold, the generator is therefore neither fine-tuned nor retrained. We did not compare the pipeline with reMD or EBMetaD, whose simulated ensembles conflate conformational heterogeneity with label flexibility, whereas ours represent conformational heterogeneity alone.

Rotamer library and accessible-volume models such as MMM or MTSSLwizard predict a width that depends on how the label rotamers are sampled^11,76^. In our benchmark, both predicted broader distributions than measured for most pairs (Fig. 1A), which obscure the width attributed to conformational heterogeneity, consistent with the overestimation of spin label flexibility by in silico labeling^12^. In SC diffusion, the temperature τ sets the width assigned to the spin label, with τ = 0.2 reflecting a design choice rather than a measured value. The remaining width, including any additional label flexibility, is consequently attributed to conformational heterogeneity. Tessmer et al.^76^ adopted the opposite convention. They calibrated the off-rotamer dihedral spread of their rotamer library model to reproduce experimental P(r) from the crystal structures. This convention attributes the entire measured width to the label and subsumes any conformational contribution. In PfMATE and MsbA, widths tend to be larger on the side toward which each transporter opens (Fig. 1G), suggesting that part of the remaining width under our convention is conformational in origin.

Calibrating τ by performing DEER on essentially static proteins, i.e., devoid of conformational heterogeneity, offers an alternative. Because spin labels are typically introduced at solvent-exposed helix and strand sites of similar environment, on which SC diffusion was trained, such a τ should be transferable between proteins. The width in excess of the calibrated label contribution would then estimate the conformational heterogeneity of other proteins, provided that the two contributions are independent, as assumed in Results.

Because SC diffusion is differentiable with respect to the structure, a single backpropagation through it into Boltz-2 yields the gradient for all optimized variables (single and pair representations). These variables number approximately 2 × 10^7^ (LmrP) to 2 × 10^8^ (Pgp) per model, or 2 × 10^9^ to 2 × 10^10^ per ensemble, which renders gradient-free approaches impractical. Each guided step incurs a computational cost linear in ensemble size and cubic in sequence length, whereas the optimized representations require memory quadratic in sequence length. A guided Pgp ATP/Mg^2+^ ensemble required approximately 250 H100 GPU-hours, whereas the PfMATE, LmrP and MhsT ensembles required approximately 15–60 L40S GPU-hours (Methods).

The generator, in turn, shapes what guidance can produce. In development runs with earlier settings, the same SC diffusion model guided Boltz-2 and OpenFold, an AlphaFold2 reimplementation, from distinct default ensembles toward the same PfMATE pH 7.5 P(r). Their weighted ensembles attained nearly identical TM-scores to 6GWH (OpenFold 0.749, Boltz-2 0.737) and to 6FHZ (OpenFold 0.613, Boltz-2 0.607), suggesting that the data and the forward model primarily determined the ensemble. The generator nevertheless remains consequential, because guidance yields realistic models only for conformations it is capable of constructing. For the synthetic targets, the ensembles overpopulated the center of PC1 in every replicate and over the full range of 3 to 40 pairs, which points to generator bias rather than an insufficient number of pairs (Supplementary Figs. 8 and 9). A differentiable all-atom generator whose unguided samples accurately cover the experimentally populated states could replace Boltz-2 with reduced guidance.

Experimental design is nonetheless critical. Three pairs selected by their predicted end-state distance difference recovered a bimodal ensemble, whereas 40 poorly discriminating pairs collapsed it onto a single mode (Supplementary Fig. 9). Because highly ranked pairs can report on the same motion, candidate pairs, rather than single sites, should be ranked by the incremental information about conformational heterogeneity that each adds to the pairs already selected. Mutual-information criteria accomplish this for DEER and Förster resonance energy transfer (FRET)^77,78^, as earlier restraint selection did for single folds^79^.

When restrained with experimental P(r), the ensembles shifted between conditions in the expected direction, albeit partially, largely reflecting the mixture of states encoded in the experimental P(r). At pH 5, the LmrP P(r) with resolved OF and IF peaks partition almost evenly (OF fractions of 54–58% at three pairs, compared with 58% of the ensemble weight closer to 6T1Z). The MhsT apo and Na^+^ targets in DM differed by a median per-pair *W*_1_ of only 0.7 Å, which accounts for the overlap of their ensembles. Among the 28 fitted pairs of Pgp, the ADP-Vi ensembles reproduced the known asymmetry of the two NBSs^6,45^, with a short-distance component at the NBS2 pair 511– 1043 but not at the NBS1 pair 400–1156 (Fig. 4C and Supplementary Fig. 17).

Because the pipeline reproduces whatever the target P(r) contains, it is sensitive to the shape of P(r). The ensembles fitted to the raw PfMATE targets contained numerous distorted structures associated with the long tails of these P(r), which we attribute to aggregation^80^ because long-distance DEER components frequently arise from it, although a conformational origin cannot be excluded. The tail-free synthetic targets yielded none (Supplementary Fig. 6), indicating that low-noise P(r) yield the most interpretable ensembles. For the same reason, the loss penalizes invalid chemistry rather than unfamiliar conformations. A loss term rewarding high Boltz-2 confidence (pLDDT) biased models back toward the generator’s own prediction, where pLDDT is highest, thereby degrading the fit. Because experimental P(r) can contain inherently low-confidence contributions, such as low-population states, only the MolProbity surrogate and the CCD bound violation term were retained (Supplementary Table 1).

To our knowledge, neither conformational free energies nor tabulated OF and IF populations have been published for PfMATE, LmrP, MhsT or Pgp. The closest anchors are protonation midpoints, pK 5.1 for the PfMATE OF-to-IF transition^38^ and pK 4.7 ± 1.3 for an LmrP outward-open to outward-closed transition^81^. Because the free energy differences depend on how the conformations are defined, alternative definitions of the states, such as assignment to the nearest reference structure, were tested. Under these definitions, free energy shifts of PfMATE, LmrP and MhsT remained in the expected direction. The low-pH shifts of PfMATE and LmrP (Fig. 5E) are of the same order as spectroscopically derived free energies of other proteins. These include approximately 0.5 kcal mol^−1^ for Glt_Ph_ (detergent versus lipid)^82^, 1.6 kcal mol^−1^ for the HCN2 cyclic NBD (ligand binding)^83^ and 1.15 kcal mol^−1^ for LeuT (Na^+^ stabilization of the closed intracellular gate)^84^.

Two caveats limit these values. The landscapes are built from *θ*^∗^ weights over models whose frequencies reflect generator sampling rather than equilibrium populations, and reweighting cannot populate unsampled states^85,86^. We also assumed 298 K, following the convention that a frozen DEER sample reports the equilibrium at the temperature prior to freezing^87^, a premise supported only indirectly^88,89^. Van’t Hoff analysis of samples equilibrated at several temperatures before freezing, as carried out for the ABC exporter TmrAB^90^, could separate enthalpy and entropy.

Templates, supplied as inputs rather than restraints, imprinted a memory on the models initiated from them without altering the ensemble fits, whereas ligand poses originated from Boltz-2 rather than from the DEER data. Cryo-EM and DEER have been combined to investigate ABC transporters^49,91^, while cryo-EM models can serve as AlphaFold templates^92^. A cryo-EM structure could therefore serve as a template for models of a DEER-guided ensemble, which would reveal the mechanistic identity of a state and its relative weight.

DEER-guided ensembles should be interpreted as hypotheses consistent with the fitted distributions rather than as determined structures. Each model carries far more optimized variables than there are fitted distributions, leading to differences between repeated runs. Although three replicates for the synthetic bimodal target differed little (Supplementary Fig. 8), each experimental condition was run only once. Because we fitted the targets as exact distributions, forward-model errors in mean distance, both a global offset and per-pair errors, propagate into the structures. Rigorous testing therefore requires additional pairs not used in the fit, together with comparison to other methods.

Any data whose forward model is differentiable with respect to the structure, such as NMR observables, crosslinks, hydrogen–deuterium exchange or single-molecule FRET, could enter the loss alongside the P(r). A single ensemble could then be fitted simultaneously to several types of data, as in integrative ensemble determination^19^. As structure generators improve, DEER-guided generation could render candidate ensembles a routine product of DEER studies.

## Methods

### Spin-center diffusion model

Spin centers are the nitroxide N1–O1 bond midpoints. The model (Fig. 1B) is an improved denoising diffusion probabilistic model^27,55^ with T = 4,000 steps, conditioned on the environment of the site, which comprises at most 512 heavy atoms of standard residues within 12 Å of its C_α_, excluding the labeled side chain. Spin centers and environment atoms were expressed in a local N–C_α_–C frame centered on C_α_. The temperature τ is the per-axis standard deviation of the initial noise x_T_ in Å, an isotropic Gaussian centered on C_α_. For a fixed atom selection, spin centers are differentiable with respect to the environment atom coordinates and to the N, C_α_ and C atoms of the site.

#### Training data

Labeling sites were non-Gly, non-Pro residues in helices or strands (DSSP^93^ code H, G, I or E) with a solvent-accessible surface area above 50 Å^2^ in 17,592 single protein chains (each chain A of a PDB entry^94^) from the OpenFold training set^57,58^. At each site, chiLife^36^ version 1.2.3 generated up to 10,000 uniformly weighted spin label rotamers using its reimplementation of MTSSLwizard^11^ for the spin label R1M.

#### Training

The noise schedule is linear, with β_t_ from 10^−4^ to 0.02. A PointNet encoder^56^ reduces the environment to a 1024-dimensional max-pooled feature, which is concatenated with x_t_ and a learned timestep embedding and passed through seven Linear–ReLU layers, each of width 512. The model was optimized with AdamW^95^ and gradient clipping at norm 1.0, with one rotamer drawn per site in each epoch (Supplementary Fig. 1).

### DEER data and targets

For T4 lysozyme (T4L), the experimental values in Fig. 1 (Supplementary Fig. 3) are the means and s.d. of 50 pairs^14^. For PfMATE (25 pairs per state)^38^ and for MsbA in the nucleotide-bound (“ATP” in Fig. 1) and apo states^4^, they are the single-Gaussian major-peak means and s.d. High-pH PfMATE data were compared with the outward-facing (OF) structure 6GWH and low-pH data with the inward-facing (IF) structure 6FHZ. The T4L and MsbA data were compared with 3LZM, AMP-PNP-bound 7PH4 and apo 9FV9. All experimental reference structures of the guided runs were released before 1 June 2023, the cutoff for the PDB structures in the training data of Boltz-2^26^. For the tail-truncated targets, each of the 84 PfMATE P(r) (42 pairs at pH 7.5 and pH 4) was restricted to bins of its native 1-Å grid. The retained bins are within 20 Å of the mode of the P(r) and have a probability of at least 0.10 times its peak. Each truncated P(r) was then renormalized. To associate distorted PfMATE models with long target tails (Results), the tail bins of each pair were defined as those in which the target P(r) is below 0.10 times its peak and more than 5 Å from its mode. The tail probability of a structure is the mean over pairs of its predicted P(r) mass in these bins.

### MMM and MTSSLwizard baselines

The 141 pairs comprise 50 T4L pairs, 25 PfMATE pairs at pH 7.5, 25 PfMATE pairs at pH 4, 20 MsbA-ATP pairs and 21 MsbA-apo pairs. MMM^9^ and MTSSLwizard^11^ predictions used the chiLife^36^ reimplementations, version 1.2.3, with label R1M and default settings. For MMM and MTSSLwizard, per-pair μ and σ are the mean and s.d. of all pairwise spin-spin distances, weighted by the products of the rotamer weights. For SC diffusion, 300 spin centers were drawn at each site by ancestral sampling at 11 values of τ from 0.1 to 3 (Supplementary Fig. 2). Ancestral sampling, used throughout Fig. 1C–G, runs all 4,000 stochastic reverse steps of the model, with τ scaling both its initial noise and the new noise added at every step. The predicted mean DEER distance μ_pred_ is the mean of all 300 × 300 pairwise spin-spin distances, with σ_pred_ their standard deviation. In Fig. 1G, the membrane side of a pair is the sign of its mean membrane-normal coordinate in OPM-oriented^96^ 6GWH and 6FHZ or, for MsbA, after superposition onto OPM-oriented 6UZL^97^, with loops predicted as cytoplasmic by DeepTMHMM^98^ on the negative side.

#### Shell center

The shell center (Fig. 1E) is the point at the mean C_α_-nitroxide distance from the C_α_ along the normalized mean of the C_α_-nitroxide unit vectors. Distances between the SC diffusion and MTSSLwizard shell centers are reported as the median over the 152 sites of the 141 pairs. The drift of the SC diffusion shell center with τ was measured at the same 152 sites, with 1,500 ancestral samples per site generated from the same noise draws at every τ.

### DEER-guided ensemble generator

#### Generator and representations

Structures were generated with Boltz-2^26^, whose diffusion sampler was replaced by the guided loop described below. Boltz-2 version 2.2.1 was run with three recycling steps and without its inference-time steering. The inputs were the sequence and a precomputed MSA, without templates or ligands unless stated otherwise. Alignments were computed with the ColabFold MMseqs2 server^99^. For PfMATE, the input sequence was that of the 6HFB crystallization construct^39^ without its His tag. For LmrP, MhsT and Pgp, the input sequences were wild type.

All Boltz-2 and SC diffusion weights were frozen. The optimized variables are each model’s single and pair representations at the Pairformer input, of sizes L×384 and L×L×128 for L input tokens, one per residue or ligand heavy atom. Because they were captured after the last of the four trunk passes, they incorporate recycling, the MSA module and, in template runs, the template module. In de novo runs, all models start from identical copies of the same single and pair representations, whereas the initial coordinates x_T_ are random noise. Models therefore differ only through their own gradients and the diffusion noise. The size of an ensemble is 96.

#### Guided steps and outer update

In each cycle, a 50-step reverse-diffusion trajectory was run on Boltz-2’s Karras-type schedule^29^, with σ falling from 2560 Å to 0. At guided steps i=0–35 (σ≥2.85 Å), gradients of the loss terms with respect to the denoised estimate x^_0_ were backpropagated through the diffusion module and the Pairformer into the representations. The representations were then updated by one Adam step^62^ at each guided step, with learning rate 0.2 and global-norm clipping at 5. The last 14 steps, 36–49, were denoised without guidance while conditioned on the optimized representations. Each cycle ended with one outer update. For this update, a separate 50-step trajectory was run from fixed noise. Its final structures are not part of the resulting ensemble. Their loss was backpropagated through all 50 steps into the representations, which were then updated by one step of a separate Adam optimizer with learning rate 0.05. Weights for each model were not optimized during guidance but fitted only offline (*θ*^∗^).

#### SC diffusion sampling during guidance

Guidance and the evaluation of resulting ensembles used deterministic denoising diffusion implicit model (DDIM) sampling^100^ with η = 0, 100 evenly spaced timesteps, τ = 0.2 and 48 spin centers per site and structure (Supplementary Fig. 24). This sampler applies τ to the initial noise alone. Values of τ other than 0.2 were used for guidance only in the runs of Supplementary Fig. 10. Each model of the ensemble used one fixed draw of spin-center starting noise, the same at every site, guided step and cycle. Because the sampler is deterministic, the spin centers are then a differentiable function of the structure, which permits the loss to be backpropagated to the structure and ensures that changes in the predicted distances originate from changes in the structure rather than from new noise.

#### Cycles and the resulting ensemble

Runs had at most five cycles, with x_T_ redrawn every cycle while representations and optimizer states were retained between cycles. From the second cycle onward, a run stopped after the cycle’s outer update if the in-run weighted EMD (W_1_^p^, Fig. 2A) of the cycle’s final-step structures (i=49) improved by less than 0.05 Å or deteriorated. The resulting ensemble consists of the final-step structure (i=49) of every model in the last cycle, relaxed and weighted offline by θ^∗^ as described below.

### DEER loss

For each of the K pairs k, the 48^2^ spin-center distances of each structure were pooled over the ensemble with uniform weight. From this pool, 48 predicted quantiles at levels 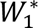 were compared with the target quantiles of the experimental P(r) at the same levels. Predicted quantiles use linear interpolation. The comparison used the one-dimensional Wasserstein-1 distance W_1,k_, the mean absolute difference between predicted and target quantiles, with each pair weighted by its own misfit, which is held constant during backpropagation and normalized to sum to one over pairs. The loss, the weighted EMD W_1_^p^=Σ_k_ W_1,k_^2^/Σ_k_ W_1,k_, is the quantity in Fig. 2A.

### MolProbity surrogate and CCD bound violation

Two reference-free terms act on x^_0_. The first, the MolProbity surrogate, adopts the form and weights of the MolProbity score^60^ without its constant, with soft versions of the clashscore c and of the percentages of rotamer outliers and of residues outside favored Ramachandran regions. Here, c is a soft count, per 1,000 heavy atoms, of heavy atom pairs more than one residue apart whose van der Waals overlap exceeds 0.40 Å, with CCP4 van der Waals radii^101^ and logistic smoothing of width 0.10 Å. The rotamer and Ramachandran terms are soft percentages of side chains with rotamer density below 0.003 and of residues with Ramachandran density below 0.02 in the Top8000 reference distributions^102,103^.

The second term, the CCD bound violation, is a linear flat-bottom penalty rather than a force field. It sums the amounts by which atom-pair distances fall below their lower or above their upper bounds. The bounds were derived from RDKit^104^ distance-bounds matrices of wwPDB Chemical Component Dictionary^61^ residues, plus peptide-junction terms. Because Boltz-2 does not construct CCD distance bounds for a single-chain protein of standard residues, the bounds of each standard residue were computed with the Boltz-2 geometry-constraint routine and evaluated with the PoseBusters^105^ potential of Boltz-2 (version 2.2.1). Peptide-junction terms were added as a bond term for the C(i)–N(i+1) distance of 1.329 Å and as angle terms for the C_α_(i)–N(i+1) and C(i)– C_α_(i+1) distances of 2.435 Å and for the O(i)–N(i+1) distance of 2.250 Å.

### Gradient-norm budget

The coordinate gradients of the DEER loss, the CCD bound violation and the MolProbity surrogate were combined by a gradient-norm budget, with norms in the ratio 85 to 5 to 10. Each regularizer gradient was rescaled to its share of the DEER gradient norm, 10/85 for the MolProbity surrogate and 5/85 for the CCD bound violation, and multiplied by the ramp w_i_=i/35 over the guided steps, with w=1 in the outer update. The resulting coefficients are the weights λ_MP_ and λ_CCD_ of Fig. 2A, which are therefore recomputed at every guided step.

### Processing after guidance

#### Relaxed ensembles

After guidance, which used no force field or energy minimization, the final structures were relaxed by restrained energy minimization as in AlphaFold2^20^. Relaxation used the AMBER ff99SB force field^106^ in OpenMM^107^ 8.2. Minimization was repeated for up to three rounds while AlphaFold2 structural violations remained.

#### Offline reweighting

The weights θ^∗^, one per model, nonnegative and summing to one, minimize the cumulative form of the mean per-pair W_1_, ℒ(θ). This loss is the mean over the K pairs of the area between the θ-weighted mean of the models’ empirical cumulative distribution functions of their 48^2^ distances and the target cumulative distribution function. Fits are reported as W_1_*, the mean over pairs of the 48-quantile W_1_ between the θ^∗^-weighted pooled predicted distances and the target quantiles.

### Synthetic recovery benchmark

The pool consisted of 500 PfMATE models generated with DEERFold^23^ by distance restrained subsampling at N_eff_=5, with restraint distances from PfMATE DEER data (Supplementary Fig. 5). The models were ranked from OF to IF by Δ_RMSD_, their C_α_ RMSD to 6GWH, an OF crystal structure, minus that to the IF crystal structure 6FHZ. The three imposed population profiles over the rank i=0,…,499, each normalized to unit sum, were OF- and IF-weighted ramps, linear between 1.0 and 0.01, and a bimodal profile, 0.5 [1+cos(2π i/500)]+0.01. For each pair, the target consisted of 48 weighted quantiles of the profile-weighted mixture of all pairwise spin-spin distances. These distances were computed by applying the SC diffusion model to each pool model with the DDIM settings used for guidance and drawing 48 spin centers per site. Targets were built for the 40 pairs whose mean distance, predicted with SC diffusion, differed most between the OF and IF reference structures. PC1 and PC2 are the first two principal components of the pool’s C_α_ coordinates at 97 sites after superposition^108^ onto the pool mean, onto which the models of the resulting ensembles were also superposed before projection. The PC1 densities in Fig. 2D are Gaussian kernel density estimates, which for resulting ensembles are unweighted and for targets weight the 500 pool models by the imposed profile, with one kernel s.d. of 4.47 PC units for all six curves.

### Ensemble analyses

TM-scores^63^ were computed with TM-align^109^ by comparing all C_α_ atoms of a structure with chain A of the reference and normalizing by the reference length. The θ^∗^ mass closer to a reference is the summed θ^∗^ of the models whose TM-score to it is higher than to the other reference.

The individual structures in Figs. 3C,E and 4B,G and Supplementary Fig. 21C were selected without reference structures. Among models with nonzero *θ*^∗^ that pass reference-free validity checks, the model shown is the one with the largest mean over pairs of the inner product between its predicted P(r) and the target P(r), both in 1-Å bins. These checks required backbone C–N distances of at most 2.0 Å and no non-bonded heavy atom contact closer than 2.2 Å. Each Na^+^ or Mg^2+^ ion was required to be coordinated by at least four O or N atoms at 2.0–3.0 Å, with none closer than 1.9 Å. Each ATP or Trp was required to contact at least ten protein heavy atoms within 4 Å.

#### Conformational coordinates

To locate each PfMATE and LmrP model along the OF-to-IF transition, the model and the second reference (6FHZ or T1024) were superposed onto the first reference (6GWH or 6T1Z) over their common C_α_ atoms. The coordinate t is the projection of the displacement of the model from the first reference onto the line d connecting the two references. It is expressed as a fraction of the length of d (t = 0 at the first reference and t = 1 at the second). Equivalently, t = ½ + (RMSD_1_^2^ − RMSD_2_^2^)/(2·RMSD_12_^2^), where RMSD_1_ and RMSD_2_ are the RMSDs of the model to the first and second reference and RMSD_12_ is that between the references, all in the superposition onto the first reference. For MhsT, the ensemble models and the candidate models c1 (OF), c5 (occluded) and c7 (IF)^44^ were superposed onto c1. The coordinate ξ is defined in the same way as t, with d running from c1 to c7, whereas η is the displacement of the model toward c5 perpendicular to d, also expressed as a fraction of the length of d.

#### RMSDs

For the RMSD between average structures, the models of each run were superposed iteratively onto their *θ*^∗^ -weighted mean by four rounds of C_α_ Kabsch superposition^108^. The weighted mean structures of two runs were then compared by C_α_ RMSD after Kabsch superposition.

### Free energies on gate coordinates

#### Gate coordinates

The gate distances *d*_EC_ and *d*_IC_ are mean C_α_–C_α_ distances over fixed extracellular and intracellular residue pairs derived from published descriptions of each protein’s gates or of homologous gates (Figs. 4D and 5A). No gate pair was among the guided DEER pairs. For PfMATE, the extracellular pairs were Ala54–Ala277, Leu53–Ala277, Pro50–Ile358 and Leu58–Val276 and the intracellular pairs Arg88–Glu310, Ala82–Ala306, Gly86–Ala306 and Gly86–Gly309^39,110^. For LmrP, the extracellular pair was Ile34–Val240^111^ and the intracellular pair Asp68–Gly348^5,41,64,112^. For MhsT, the extracellular pairs were Arg34–Asp385 and Tyr108– Phe230 and the intracellular pairs Glu10–Arg344 and Tyr245–Glu330^43^, with all but Glu10– Arg344 mapped from the LeuT gate pairs^113–115^. For Pgp, in mouse numbering, the extracellular pairs were Ala79–Thr736^116,117^ and Tyr306–Tyr949^118^ and the intracellular pairs Lys429–Ser1173 and Ser528–Lys1072^116,117^.

#### Free energies

The landscapes of Figs. 4E and 5B–D were computed as *G* = −*RT* ln *p*, where p is the θ^∗^-weighted Gaussian kernel density of the models of each resulting ensemble on (*d*_EC_, *d*_IC_). Figures 2E, 4F and 5E and Supplementary Table 2 report *G*_state_ − *G*_OF_ of these landscapes at fixed positions (d_EC_, d_IC_) at an assumed 298.15 K, with positive values favoring OF. The positions are shared by all runs of a data set (the experimental runs of one protein, or the synthetic runs). For PfMATE and LmrP, the OF positions were the deepest minima nearer an OF reference in the landscapes obtained by averaging the densities of the two runs in Fig. 5B or 5C with equal weight, at (16.4, 5.3) and (21.3, 5.7) Å, respectively. The IF positions were the deepest minima nearer an IF reference in the low-pH landscapes of PfMATE and LmrP, at (7.5, 13.8) and (7.0, 28.6) Å, respectively, because the global minima of these landscapes reside on the OF side. For the synthetic targets of Fig. 2, the OF-weighted and IF-weighted resulting ensembles replaced the two conditions, yielding (13.5, 5.1) and (6.2, 16.2) Å. For MhsT, the positions were the gate coordinates of the candidate models c1 (OF), c5 (occluded) and c7 (IF), (13.7, 11.2), (11.0, 14.2) and (11.1, 22.5) Å. For Pgp, the OF position was the minimum nearer 7ZK4 in the landscape obtained by averaging the apo and ATP/Mg^2+^ densities with equal weight, at (23.8, 15.8) Å. The IF and occluded positions were the minima of the apo and ATP/Mg^2+^ landscapes, at (16.5, 45.1) and (15.7, 31.6) Å, respectively.

### Template runs

Templates act solely through the Boltz-2 template module. The templates were 6GWH and 6FHZ for PfMATE, 6T1Z and T1024 for LmrP, and c1, c5 and c7 for MhsT. The 6T1Z template file was rebuilt to the full sequence, whereas distances to 6T1Z were measured on the experimental structure. Models were assigned cyclically to the trunk representations computed without a template (de novo) or with one of the templates. Each run therefore contains its own de novo group.

The gain of template T is the unweighted median C_α_ RMSD to T of the run’s de novo models minus that of its T-seeded models. Its specificity is this gain minus the mean of the same difference measured to each other template S of the run. The gain, the specificity and Supplementary Fig. 18E use C_α_ RMSDs to the templates over residues common by number. In Supplementary Fig. 18F, the 14 runs are the de novo and template runs of the two PfMATE conditions with tail-truncated targets, the two LmrP conditions and the three MhsT DM conditions.

### Ligand runs

Ligands were unconstrained input entities specified by Chemical Component Dictionary codes^61^. They were two Na^+^ with or without Trp for MhsT, and two ATP and two Mg^2+^ for Pgp. The loss was computed on protein atoms only. The inference-only controls were 96 unconstrained Boltz-2 predictions each from the inputs of the MhsT Na^+^/Trp ligand runs and the Pgp ATP/Mg^2+^ run, with the 50 steps of the guided runs.

### Ligand placement metrics

Ligand placement errors, measured in C_α_-fitted pocket frames of 4US3 (MhsT) or 7ZK4 (Pgp), are heavy atom RMSDs computed in these frames without further superposition (ATP phosphate oxygens not symmetrized) or, for ions, single atom distances. The pocket frames used the residues within 8.0 Å of the crystal ligands for MhsT and, for Pgp, of each site’s ligands in the NBD that contributes its Walker A motif. Pgp ligand copies were assigned to sites by pocket. The two Na^+^ ions were assigned to Na1 (which coordinates the Trp carboxylate in 4US3) and Na2 by minimum summed distance. The centroid offset, the systematic error of the distribution of ligand positions, is the distance between the mean ligand centroid and the reference-structure centroid. For an ion, the spread is the median distance of its positions from their mean. All placement statistics are unweighted.

### Computational costs

Runs of Supplementary Table 1 used 8 or 16 NVIDIA L40S GPUs, except the Pgp runs, which used 8 H100 GPUs. A guided ensemble required approximately 15–60 L40S GPU-hours for PfMATE, LmrP and MhsT, compared with approximately 250 H100 GPU-hours for the Pgp ATP/Mg^2+^ ensemble.

## Data availability

The resulting ensembles, the DEER targets, the MhsT candidate models c1–c7 and the data underlying the figures are available at [https://github.com/CAAIPD/SC_diffusion]. The DEER data were obtained from published studies^4–6,14,38,40,44,45^. The reference structures are available from the Protein Data Bank under accession codes 3LZM, 6GWH, 6FHZ, 6HFB, 7PH4, 9FV9, 6UZL, 6T1Z, 4US3, 7A65 and 7ZK4.

## Code availability

The code for SC diffusion, DEER-guided generation and all analyses, together with the trained SC checkpoint, is available at [https://github.com/CAAIPD/SC_diffusion].

## Author contributions

T.-Y.K. and H.S.M. conceived the study. T.-Y.K. developed SC diffusion and the DEER-guided pipeline, performed the computations and analyzed the data. H.S.M. supervised the project. T.-Y.K. and H.S.M. wrote the manuscript. J.M. discussed and edited the manuscript.

