## Supplementary for "Differentiable Spin-Center Diffusion Model Enables DEER-Guided Generative Pipeline of Weighted Protein Ensembles"

[1]Department of Molecular Physiology and Biophysics, Vanderbilt University, Nashville, TN, USA. [2]Center for Applied Artificial Intelligence in Protein Dynamics, Vanderbilt University, Nashville, TN, USA. [3]Department of Chemistry, Vanderbilt University, Nashville, TN, USA. [4]Center for Structural Biology, Vanderbilt University, Nashville, TN, USA. [5]Institute of Drug Discovery, Faculty of Medicine, Leipzig University, Leipzig, Germany.

*

### Supplementary Tables

#### Supplementary Table 1. Properties of DEER-guided ensembles

The clashscore, rotamer outliers, Ramachandran outliers and MolProbity score were obtained with the MolProbity surrogate for the 96 structures of each resulting ensemble after relaxation (Methods). Each entry reports the median and, in parentheses, the 5th–95th percentile over the 96 structures, without weights. Hydrogens were added before scoring. Only protein atoms were scored, not ligands or ions. The clashscore is the number of all-atom overlaps of at least 0.4 Å per 1,000 atoms, whereas rotamer and Ramachandran outliers are expressed as percentages of the evaluated residues. OF and IF denote the outward-facing and inward-facing conformations.

| DEER target | Run type | Pair number | $W_1^*$ (Å) | Clashscore | Rotamer outliers (%) | Ramachandran outliers (%) | MolProbity score |
|---|---|---|---|---|---|---|---|
| *PfMATE* | | | | | | | |
| pH 7.5 | de novo | 42 | 0.7138 | 5.3 (1.8–13.9) | 1.07 (0.27–2.94) | 0.32 (0.00–1.48) | 1.45 (1.00–2.37) |
| pH 4 | de novo | 42 | 0.7140 | 5.5 (1.5–12.2) | 1.07 (0.20–3.28) | 0.21 (0.00–1.11) | 1.45 (0.88–2.26) |
| pH 7.5, tail-truncated | de novo | 42 | 0.6808 | 2.3 (0.7–6.3) | 0.53 (0.00–1.60) | 0.00 (0.00–0.42) | 1.03 (0.72–1.49) |
| pH 4, tail-truncated | de novo | 42 | 0.6388 | 2.7 (0.8–9.0) | 0.53 (0.00–1.67) | 0.21 (0.00–0.69) | 1.11 (0.75–1.71) |
| pH 7.5 | template (de novo, 6GWH, 6FHZ) | 42 | 0.6868 | 6.0 (2.0–14.0) | 1.07 (0.27–2.94) | 0.21 (0.00–1.27) | 1.48 (0.98–2.36) |
| pH 4 | template (de novo, 6GWH, 6FHZ) | 42 | 0.7647 | 5.6 (1.6–13.1) | 1.07 (0.27–3.21) | 0.21 (0.00–1.27) | 1.47 (1.00–2.33) |
| pH 7.5, tail-truncated | template (de novo, 6GWH, 6FHZ) | 42 | 0.6574 | 2.5 (0.5–10.1) | 0.80 (0.00–1.87) | 0.00 (0.00–0.42) | 1.05 (0.69–1.92) |
| pH 4, tail-truncated | template (de novo, 6GWH, 6FHZ) | 42 | 0.6565 | 3.0 (0.8–10.1) | 0.80 (0.00–1.94) | 0.21 (0.00–0.63) | 1.12 (0.75–1.95) |
| *LmrP* | | | | | | | |
| pH 5 | de novo | 9 | 0.1512 | 1.4 (0.3–4.5) | 0.29 (0.00–1.17) | 0.00 (0.00–0.99) | 0.92 (0.61–1.53) |
| pH 8 | de novo | 9 | 0.1367 | 0.6 (0.1–2.1) | 0.29 (0.00–0.59) | 0.00 (0.00–0.49) | 0.70 (0.55–1.02) |
| pH 5 | template (de novo, 6T1Z, T1024) | 9 | 0.1759 | 1.0 (0.2–4.7) | 0.29 (0.00–1.17) | 0.00 (0.00–0.74) | 0.83 (0.56–1.54) |
| pH 8 | template (de novo, 6T1Z, T1024) | 9 | 0.1499 | 0.8 (0.0–4.2) | 0.29 (0.00–1.17) | 0.00 (0.00–0.49) | 0.74 (0.50–1.26) |
| *MhsT, detergent (DM, 22 pairs) and nanodiscs (ND, 20 pairs, a subset of the DM pairs)* | | | | | | | |
| DM apo | de novo | 22 | 0.3481 | 2.7 (1.0–6.6) | 0.58 (0.00–1.44) | 0.22 (0.00–0.67) | 1.09 (0.80–1.68) |
| DM $Na^+$ | de novo | 22 | 0.3052 | 3.5 (1.1–7.9) | 0.58 (0.00–1.73) | 0.22 (0.00–0.73) | 1.22 (0.82–1.75) |
| DM $Na^+$/Trp | de novo | 22 | 0.2693 | 3.5 (1.4–9.2) | 0.58 (0.00–1.44) | 0.22 (0.00–0.67) | 1.21 (0.88–1.84) |
| DM apo | template (de novo, c1, c5, c7) | 22 | 0.2818 | 2.9 (1.1–9.1) | 0.58 (0.00–1.51) | 0.22 (0.00–0.67) | 1.18 (0.85–1.84) |
| DM $Na^+$ | template (de novo, c1, c5, c7) | 22 | 0.2961 | 2.5 (1.0–7.7) | 0.58 (0.00–1.73) | 0.22 (0.00–0.67) | 1.12 (0.82–1.80) |
| DM $Na^+$/Trp | template (de novo, c1, c5, c7) | 22 | 0.2534 | 3.2 (1.2–8.6) | 0.58 (0.00–1.44) | 0.22 (0.00–0.67) | 1.18 (0.85–1.85) |
| ND apo | de novo | 20 | 0.2719 | 3.6 (1.0–8.9) | 0.58 (0.00–1.44) | 0.22 (0.00–0.67) | 1.26 (0.84–1.82) |
| ND $Na^+$ | de novo | 20 | 0.2769 | 3.1 (0.9–7.7) | 0.58 (0.00–1.51) | 0.22 (0.00–0.67) | 1.17 (0.79–1.75) |
| ND $Na^+$/Trp | de novo | 20 | 0.2921 | 3.0 (1.0–7.8) | 0.58 (0.00–1.73) | 0.00 (0.00–0.67) | 1.11 (0.81–1.74) |

| DEER target | Run type | Pair number | $W_1^*$ (Å) | Clashscore | Rotamer outliers (%) | Ramachandran outliers (%) | MolProbity score |
|---|---|---|---|---|---|---|---|
| ND apo | template (de novo, c1, c5, c7) | 20 | 0.2746 | 3.7 (1.3–8.5) | 0.58 (0.00–1.51) | 0.22 (0.00–0.67) | 1.22 (0.92–1.84) |
| ND $Na^+$ | template (de novo, c1, c5, c7) | 20 | 0.2714 | 3.3 (0.8–7.8) | 0.58 (0.00–1.15) | 0.00 (0.00–0.67) | 1.17 (0.76–1.62) |
| ND $Na^+$/Trp | template (de novo, c1, c5, c7) | 20 | 0.2631 | 2.5 (1.0–8.0) | 0.58 (0.00–1.15) | 0.00 (0.00–0.44) | 1.08 (0.82–1.64) |
| DM $Na^+$ | ligand (2 $Na^+$) | 22 | 0.2936 | 3.4 (1.0–8.9) | 0.86 (0.00–1.51) | 0.22 (0.00–0.89) | 1.25 (0.84–1.78) |
| DM $Na^+$/Trp | ligand (2 $Na^+$, Trp) | 22 | 0.3021 | 3.6 (1.3–7.3) | 0.58 (0.00–1.73) | 0.22 (0.00–0.67) | 1.21 (0.85–1.86) |
| ND $Na^+$ | ligand (2 $Na^+$) | 20 | 0.2919 | 3.6 (1.3–8.0) | 0.58 (0.00–1.88) | 0.22 (0.00–0.73) | 1.25 (0.94–1.90) |
| ND $Na^+$/Trp | ligand (2 $Na^+$, Trp) | 20 | 0.2472 | 3.2 (1.1–7.5) | 0.58 (0.00–1.73) | 0.22 (0.00–0.67) | 1.18 (0.83–1.74) |
| *Pgp* | | | | | | | |
| apo | de novo | 28 | 0.4761 | 3.1 (1.2–10.1) | 0.57 (0.26–3.38) | 0.24 (0.00–1.14) | 1.18 (0.89–2.24) |
| ADP-Vi | de novo | 28 | 0.4597 | 5.2 (2.4–16.3) | 0.95 (0.28–4.00) | 0.47 (0.08–2.02) | 1.53 (1.11–2.44) |
| ADP-Vi | ligand (2 ATP, 2 $Mg^{2+}$) | 28 | 0.5358 | 5.0 (2.5–16.9) | 1.09 (0.26–7.27) | 0.47 (0.16–3.46) | 1.60 (1.07–3.08) |
| *PfMATE, synthetic targets from the 500-model pool* | | | | | | | |
| OF-weighted ramp | de novo | 40 | 0.3321 | 0.5 (0.1–1.5) | 0.00 (0.00–0.53) | 0.00 (0.00–0.21) | 0.70 (0.55–0.89) |
| IF-weighted ramp | de novo | 40 | 0.2850 | 0.4 (0.0–1.9) | 0.14 (0.00–0.60) | 0.00 (0.00–0.21) | 0.69 (0.50–0.95) |
| bimodal | de novo | 40 | 0.3470 | 0.5 (0.1–2.0) | 0.00 (0.00–0.80) | 0.00 (0.00–0.21) | 0.69 (0.55–0.97) |
| bimodal | de novo, seed replicate | 40 | 0.3073 | 0.5 (0.1–1.1) | 0.00 (0.00–0.53) | 0.00 (0.00–0.21) | 0.69 (0.54–0.81) |
| bimodal | de novo, seed replicate | 40 | 0.2992 | 0.5 (0.1–2.0) | 0.00 (0.00–0.60) | 0.00 (0.00–0.42) | 0.69 (0.55–1.00) |

#### Supplementary Table 2. Free energies from gate distances

**a** $\Delta G$ of each run (kcal $mol^{-1}$), obtained as the landscape values $G_{IF} - G_{OF}$ and, for MhsT and Pgp, $G_{occ} - G_{OF}$ (occ, occluded conformation) at the positions of Figs. 4F and 5E (Methods).

| DEER target | Run type | $G_{IF} - G_{OF}$ | $G_{occ} - G_{OF}$ |
|---|---|---|---|
| *PfMATE, landscape values* | | | |
| pH 7.5 | de novo | +0.90 | – |
| pH 4 | de novo | −0.15 | – |
| pH 7.5, tail-truncated | de novo | +2.32 | – |
| pH 4, tail-truncated | de novo | −0.38 | – |
| pH 7.5, tail-truncated | template | > +2.53 | – |
| pH 4, tail-truncated | template | −0.20 | – |
| *LmrP, landscape values* | | | |
| pH 5 | de novo | +0.24 | – |
| pH 8 | de novo | +1.69 | – |
| pH 5 | template | +0.14 | – |
| pH 8 | template | +1.72 | – |
| *MhsT in detergent (DM) and nanodiscs (ND), landscape values* | | | |

| DEER target | Run type | $G_{IF} - G_{OF}$ | $G_{occ} - G_{OF}$ |
|---|---|---|---|
| DM apo | de novo | +1.56 | +0.20 |
| DM $Na^+$ | de novo | +1.01 | +0.55 |
| DM $Na^+$/Trp | de novo | +0.53 | −0.22 |
| DM apo | template | +1.05 | +0.28 |
| DM $Na^+$ | template | +1.11 | +0.04 |
| DM $Na^+$/Trp | template | −0.05 | +0.09 |
| ND apo | de novo | +0.26 | −0.59 |
| ND $Na^+$ | de novo | +0.90 | −0.42 |
| ND $Na^+$/Trp | de novo | −0.67 | −1.35 |
| ND apo | template | +0.56 | −0.04 |
| ND $Na^+$ | template | +0.40 | −0.10 |
| ND $Na^+$/Trp | template | −0.61 | −0.97 |
| DM $Na^+$ | ligand | > +2.32 | +0.24 |
| DM $Na^+$/Trp | ligand | +0.38 | −0.95 |
| ND $Na^+$ | ligand | +1.01 | −0.01 |
| ND $Na^+$/Trp | ligand | < −1.57 | < −2.67 |
| *Pgp, landscape values* | | | |
| apo | de novo | −1.77 | −1.14 |
| ADP-Vi | de novo | +0.71 | −0.08 |
| ADP-Vi | ligand | +0.33 | −0.32 |

**b** ΔΔ*G* between conditions (kcal $mol^{-1}$), computed as the condition named first minus the other from the unrounded values of part a.

| Contrast | Run type | ΔΔ*G*, IF − OF | ΔΔ*G*, occ − OF |
|---|---|---|---|
| *PfMATE, landscape values* | | | |
| pH 4 − pH 7.5 | de novo | −1.05 | – |
| pH 4 − pH 7.5, tail-truncated | de novo | −2.70 | – |
| pH 4 − pH 7.5, tail-truncated | template | < −2.73 | – |
| *LmrP, landscape values* | | | |
| pH 5 − pH 8 | de novo | −1.45 | – |
| pH 5 − pH 8 | template | −1.58 | – |
| *MhsT, landscape values* | | | |
| DM: $Na^+$ − apo | de novo | −0.55 | +0.35 |
| DM: $Na^+$/Trp − apo | de novo | −1.03 | −0.42 |
| DM: $Na^+$/Trp − $Na^+$ | de novo | −0.48 | −0.77 |
| ND: $Na^+$ − apo | de novo | +0.63 | +0.17 |

| Contrast | Run type | $\Delta\Delta G$, IF − OF | $\Delta\Delta G$, occ − OF |
|---|---|---|---|
| ND: $Na^+$/Trp − apo | de novo | −0.93 | −0.76 |
| ND: $Na^+$/Trp − $Na^+$ | de novo | −1.56 | −0.94 |
| DM: $Na^+$ − apo | template | +0.05 | −0.25 |
| DM: $Na^+$/Trp − apo | template | −1.11 | −0.20 |
| DM: $Na^+$/Trp − $Na^+$ | template | −1.16 | +0.05 |
| ND: $Na^+$ − apo | template | −0.16 | −0.06 |
| ND: $Na^+$/Trp − apo | template | −1.17 | −0.93 |
| ND: $Na^+$/Trp − $Na^+$ | template | −1.01 | −0.87 |
| DM: $Na^+$/Trp − $Na^+$ | ligand | < −1.94 | −1.19 |
| ND: $Na^+$/Trp − $Na^+$ | ligand | < −2.58 | < −2.66 |
| apo: ND − DM | de novo | −1.29 | −0.79 |
| $Na^+$: ND − DM | de novo | −0.11 | −0.96 |
| $Na^+$/Trp: ND − DM | de novo | −1.20 | −1.13 |
| apo: ND − DM | template | −0.49 | −0.33 |
| $Na^+$: ND − DM | template | −0.71 | −0.14 |
| $Na^+$/Trp: ND − DM | template | −0.56 | −1.06 |
| *Pgp, landscape values* | | | |
| ADP-Vi − apo | de novo | +2.48 | +1.06 |
| ATP/$Mg^{2+}$ run − ADP-Vi | ligand − de novo | −0.38 | −0.24 |
| ATP/$Mg^{2+}$ run − apo | ligand − de novo | +2.10 | +0.82 |

#### Supplementary Figures

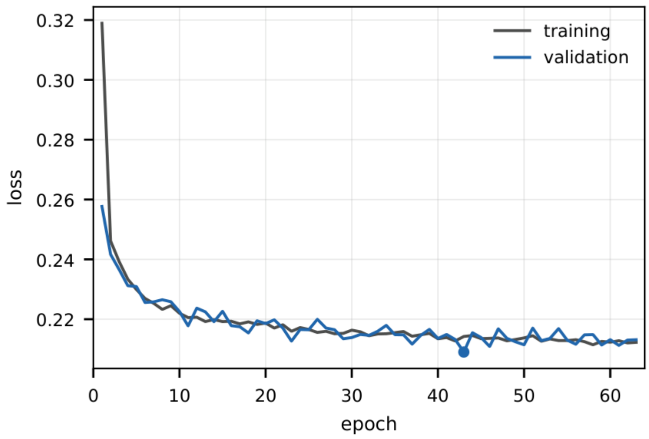


**Supplementary Fig. 1. Training and validation loss of the spin-center (SC) diffusion model.** The loss (y axis) is MSE(ϵ) + $\lambda L_{vlb}$ with $\lambda = 10^{-3}$, the hybrid objective of the improved denoising diffusion probabilistic model[1]. MSE(ϵ) is the mean squared error between the added and the predicted noise ϵ, whereas $L_{vlb}$ is the variational lower-bound term, which trains the learned variances. The proteins were split at random, without sequence-similarity clustering, into 15,833 for training and 1,759 (10%) for validation, which ensures that no validation protein contributes training sites. Training used the AdamW optimizer with a learning rate of $5 \times 10^{-4}$, cosine annealing over a nominal 100 epochs and a batch size of 512. Because the training loss is the mean over the minibatches of each epoch, it lags the model at the end of that epoch, whereas the validation loss is evaluated on the held-out proteins at the end of each epoch. Training was stopped early after 20 epochs without improvement of the validation loss (epoch 63). The checkpoint with the lowest validation loss (dot, epoch 43, validation loss 0.2090, training loss 0.2142) was retained and is the model used for all SC diffusion predictions in this work. The loss values are not used to measure how well the model reproduces MTSSLwizard. That comparison is presented in Fig. 1C–E.

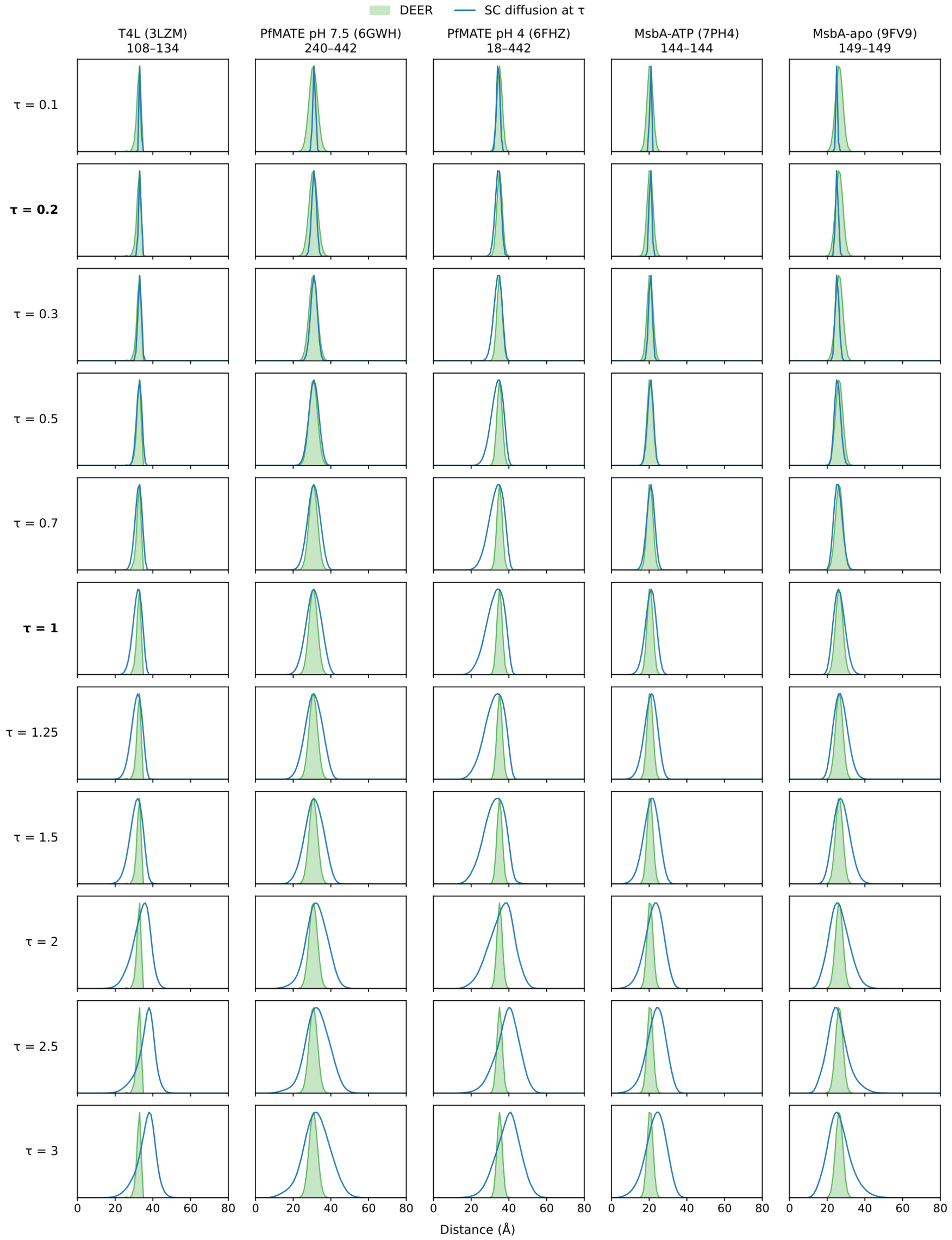


**Supplementary Fig. 2. Representative SC diffusion distance distributions across the sampling temperature τ.** Rows correspond to the eleven τ values of Fig. 1D (0.1–3), with the two temperatures of Fig. 1C, τ = 0.2 and 1, in bold. Columns correspond to the five structures: T4L (3LZM), PfMATE pH 7.5 (6GWH), PfMATE pH 4 (6FHZ), MsbA-ATP (7PH4) and MsbA-apo (9FV9). Each MsbA pair was labeled at the same residue on chains A and B. Blue, SC diffusion at the τ of the row, with 1,500 spin centers per site drawn by full 4,000-step ancestral sampling as in Fig. 1D, independently of Supplementary Fig. 3. Green, DEER, as in

Supplementary Fig. 3, repeated in every row. As in that figure, the PfMATE and MsbA DEER curves are single Gaussians for the major peak only (Methods). All curves use 1-Å bins and are drawn as lines through the bin centers without smoothing, over 0–80 Å in every panel. Each curve is scaled to the maximum of its full distribution, which permits comparison of peak positions and widths but not of areas.

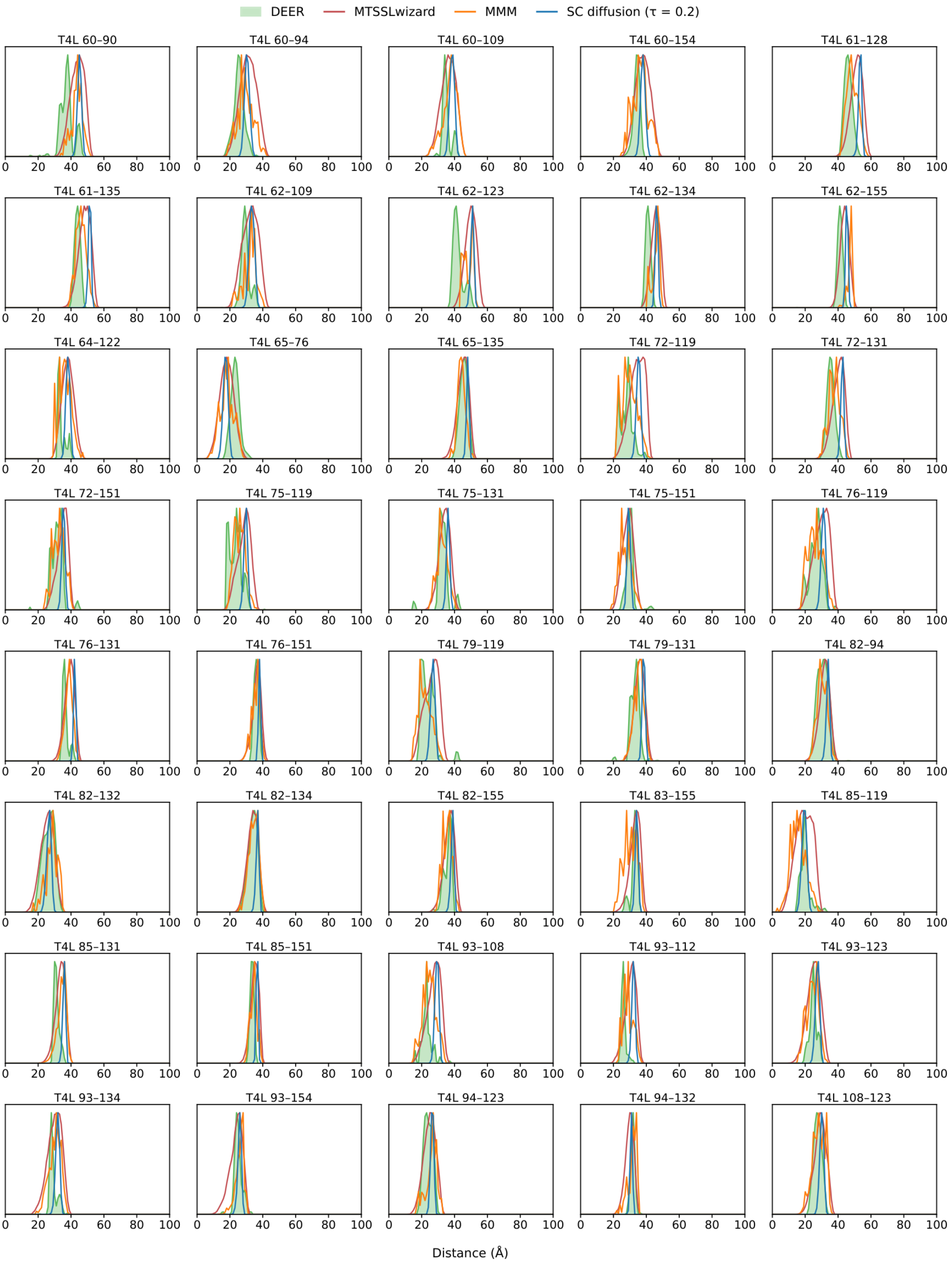
DEER
MTSSLwizard
MMM
SC diffusion (τ = 0.2)
T4L 60–90
T4L 60–94
T4L 60–109
T4L 60–154
T4L 61–128
T4L 61–135
T4L 62–109
T4L 62–123
T4L 62–134
T4L 62–155
T4L 64–122
T4L 65–76
T4L 65–135
T4L 72–119
T4L 72–131
T4L 72–151
T4L 75–119
T4L 75–131
T4L 75–151
T4L 76–119
T4L 76–131
T4L 76–151
T4L 79–119
T4L 79–131
T4L 82–94
T4L 82–132
T4L 82–134
T4L 82–155
T4L 83–155
T4L 85–119
T4L 85–131
T4L 85–151
T4L 93–108
T4L 93–112
T4L 93–123
T4L 93–134
T4L 93–154
T4L 94–123
T4L 94–132
T4L 108–123
0
20
40
60
80
100
Distance (Å)

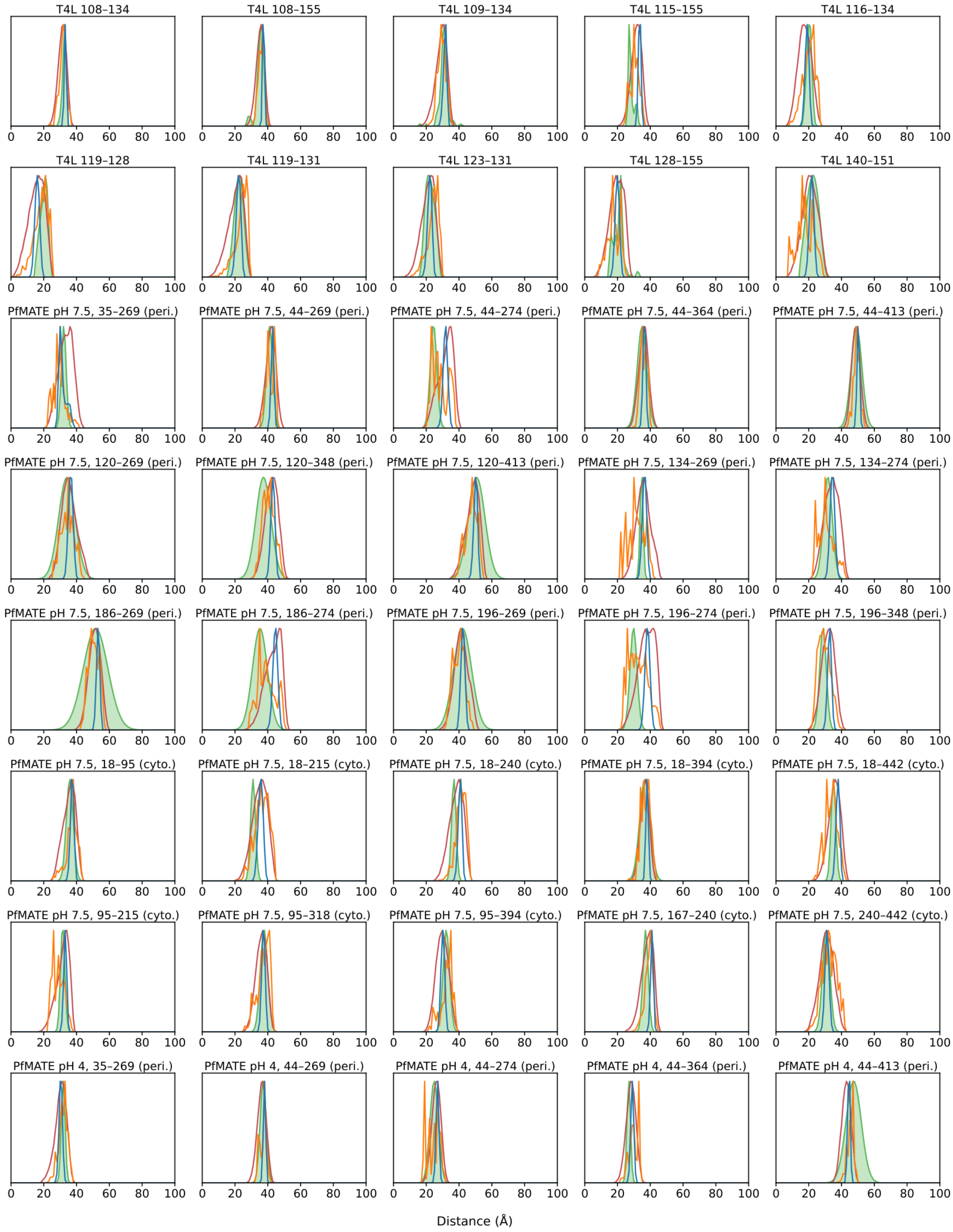
T4L 108–134
T4L 108–155
T4L 109–134
T4L 115–155
T4L 116–134
T4L 119–128
T4L 119–131
T4L 123–131
T4L 128–155
T4L 140–151
PfMATE pH 7.5, 35–269 (peri.)
PfMATE pH 7.5, 44–269 (peri.)
PfMATE pH 7.5, 44–274 (peri.)
PfMATE pH 7.5, 44–364 (peri.)
PfMATE pH 7.5, 44–413 (peri.)
PfMATE pH 7.5, 120–269 (peri.)
PfMATE pH 7.5, 120–348 (peri.)
PfMATE pH 7.5, 120–413 (peri.)
PfMATE pH 7.5, 134–269 (peri.)
PfMATE pH 7.5, 134–274 (peri.)
PfMATE pH 7.5, 186–269 (peri.)
PfMATE pH 7.5, 186–274 (peri.)
PfMATE pH 7.5, 196–269 (peri.)
PfMATE pH 7.5, 196–274 (peri.)
PfMATE pH 7.5, 196–348 (peri.)
PfMATE pH 7.5, 18–95 (cyto.)
PfMATE pH 7.5, 18–215 (cyto.)
PfMATE pH 7.5, 18–240 (cyto.)
PfMATE pH 7.5, 18–394 (cyto.)
PfMATE pH 7.5, 18–442 (cyto.)
PfMATE pH 7.5, 95–215 (cyto.)
PfMATE pH 7.5, 95–318 (cyto.)
PfMATE pH 7.5, 95–394 (cyto.)
PfMATE pH 7.5, 167–240 (cyto.)
PfMATE pH 7.5, 240–442 (cyto.)
PfMATE pH 4, 35–269 (peri.)
PfMATE pH 4, 44–269 (peri.)
PfMATE pH 4, 44–274 (peri.)
PfMATE pH 4, 44–364 (peri.)
PfMATE pH 4, 44–413 (peri.)
0 20 40 60 80 100
Distance (Å)

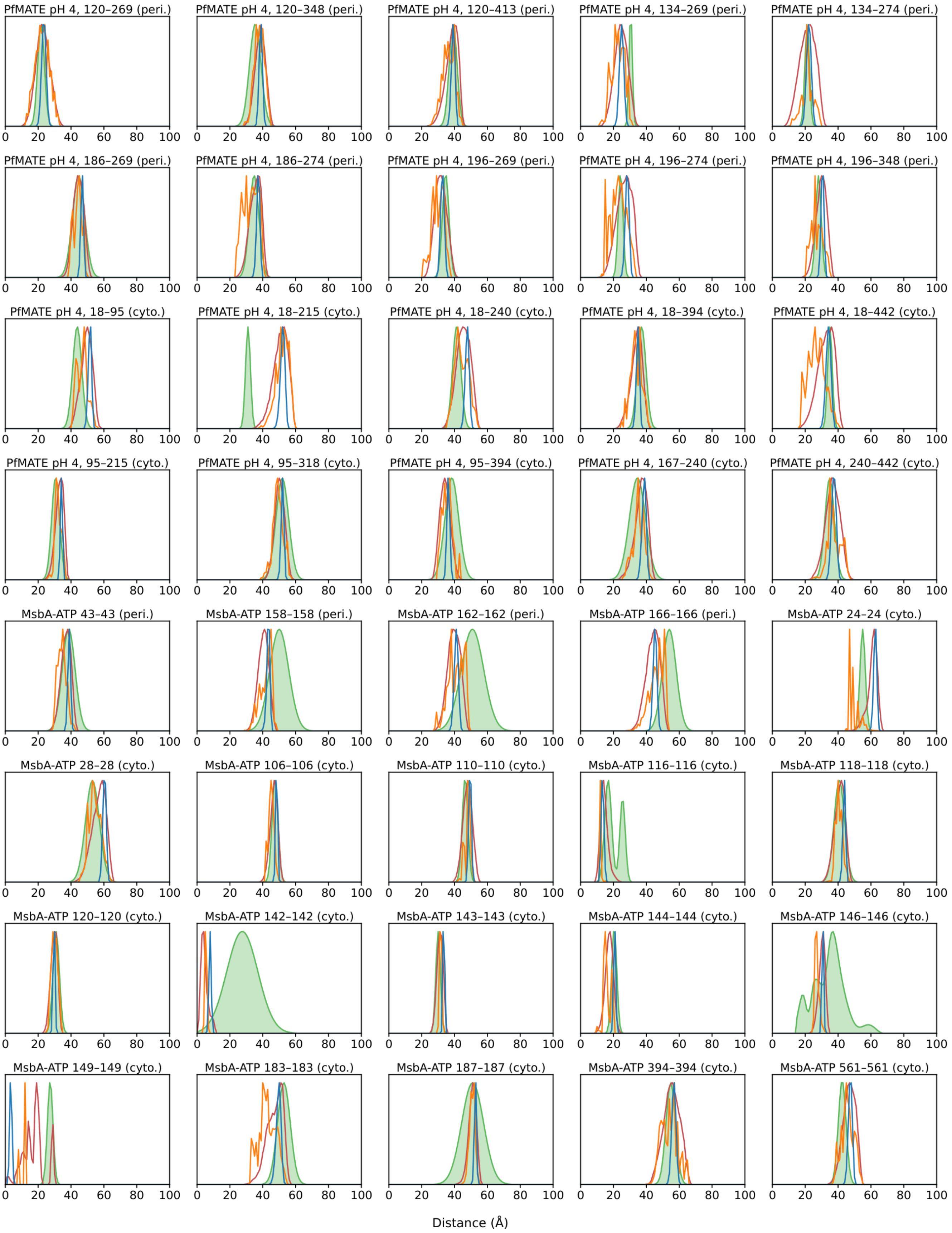

PfMATE pH 4, 120–269 (peri.)
PfMATE pH 4, 120–348 (peri.)
PfMATE pH 4, 120–413 (peri.)
PfMATE pH 4, 134–269 (peri.)
PfMATE pH 4, 134–274 (peri.)
PfMATE pH 4, 186–269 (peri.)
PfMATE pH 4, 186–274 (peri.)
PfMATE pH 4, 196–269 (peri.)
PfMATE pH 4, 196–274 (peri.)
PfMATE pH 4, 196–348 (peri.)
PfMATE pH 4, 18–95 (cyto.)
PfMATE pH 4, 18–215 (cyto.)
PfMATE pH 4, 18–240 (cyto.)
PfMATE pH 4, 18–394 (cyto.)
PfMATE pH 4, 18–442 (cyto.)
PfMATE pH 4, 95–215 (cyto.)
PfMATE pH 4, 95–318 (cyto.)
PfMATE pH 4, 95–394 (cyto.)
PfMATE pH 4, 167–240 (cyto.)
PfMATE pH 4, 240–442 (cyto.)
MsbA-ATP 43–43 (peri.)
MsbA-ATP 158–158 (peri.)
MsbA-ATP 162–162 (peri.)
MsbA-ATP 166–166 (peri.)
MsbA-ATP 24–24 (cyto.)
MsbA-ATP 28–28 (cyto.)
MsbA-ATP 106–106 (cyto.)
MsbA-ATP 110–110 (cyto.)
MsbA-ATP 116–116 (cyto.)
MsbA-ATP 118–118 (cyto.)
MsbA-ATP 120–120 (cyto.)
MsbA-ATP 142–142 (cyto.)
MsbA-ATP 143–143 (cyto.)
MsbA-ATP 144–144 (cyto.)
MsbA-ATP 146–146 (cyto.)
MsbA-ATP 149–149 (cyto.)
MsbA-ATP 183–183 (cyto.)
MsbA-ATP 187–187 (cyto.)
MsbA-ATP 394–394 (cyto.)
MsbA-ATP 561–561 (cyto.)
0 20 40 60 80 100
Distance (Å)

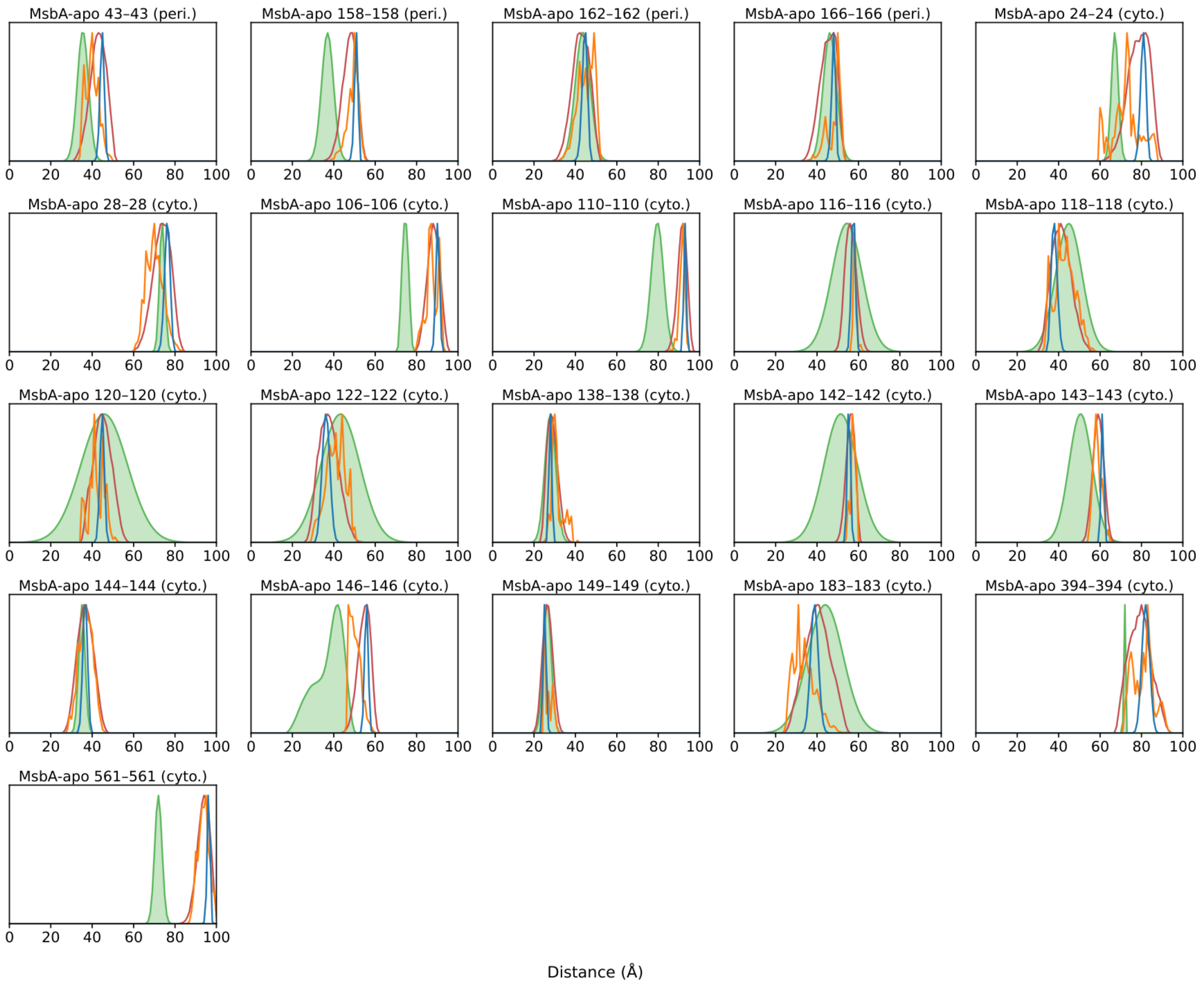


**Supplementary Fig. 3. Per-pair distance distributions for the DEER pairs of Fig. 1: DEER, MTSSLwizard, MMM and SC diffusion.** Each panel shows one pair of Fig. 1. The 141 pairs comprise 50 T4L pairs, 25 PfMATE pairs per state, 20 MsbA-ATP pairs and 21 MsbA-apo pairs. Panel titles list the protein, the labeled residues and, for membrane proteins, the side (peri., periplasmic, or cyto., cytoplasmic) assigned as in Fig. 1G. Each MsbA pair was labeled at the same residue on chains A and B. Each panel overlays the experimental $P(r)$ (green, filled), MTSSLwizard (red), MMM (orange) and SC diffusion at $\tau = 0.2$ (blue, 300 spin centers per site, drawn with the 4,000-step ancestral sampler of Fig. 1 independently of the samples in that figure). Curves use 1-Å bins without smoothing and are scaled to their own maximum, which precludes a comparison of areas. Every panel spans 0–100 Å. DEER data are from T4L[2], PfMATE[3] and MsbA[4].

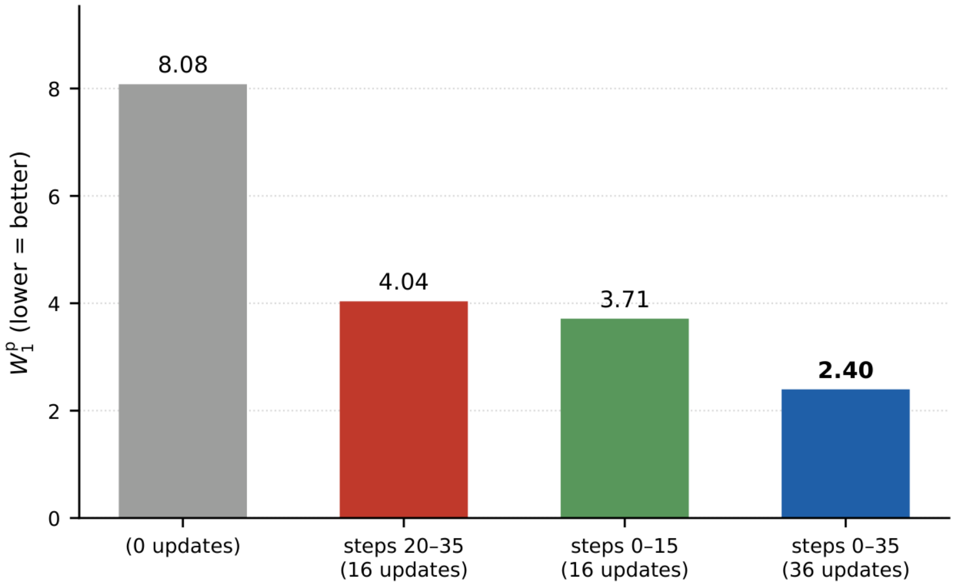


**Supplementary Fig. 4. Guiding at high-noise diffusion steps improves the DEER fit efficiently.** Four PfMATE de novo runs against the pH 7.5 target (42 pairs), each with $N = 8$ structures, one 50-step trajectory and the same seed, differ only in the diffusion steps at which guidance updates were applied. The labels below the bars indicate the guided steps and the number of updates, with step 0 the first, highest-noise step. The four runs used no guidance, guidance at the lower-noise steps 20–35 (16 updates), guidance at the high-noise steps 0–15 (16 updates) and guidance throughout the high- and lower-noise steps (0–35, 36 updates), as in the main runs. Bar heights and the numbers above the bars report the $W_1{}^p$ logged by each run (Å, lower is better). These values compare where guidance is applied at equal $N$ and seed but do not measure ensemble quality.

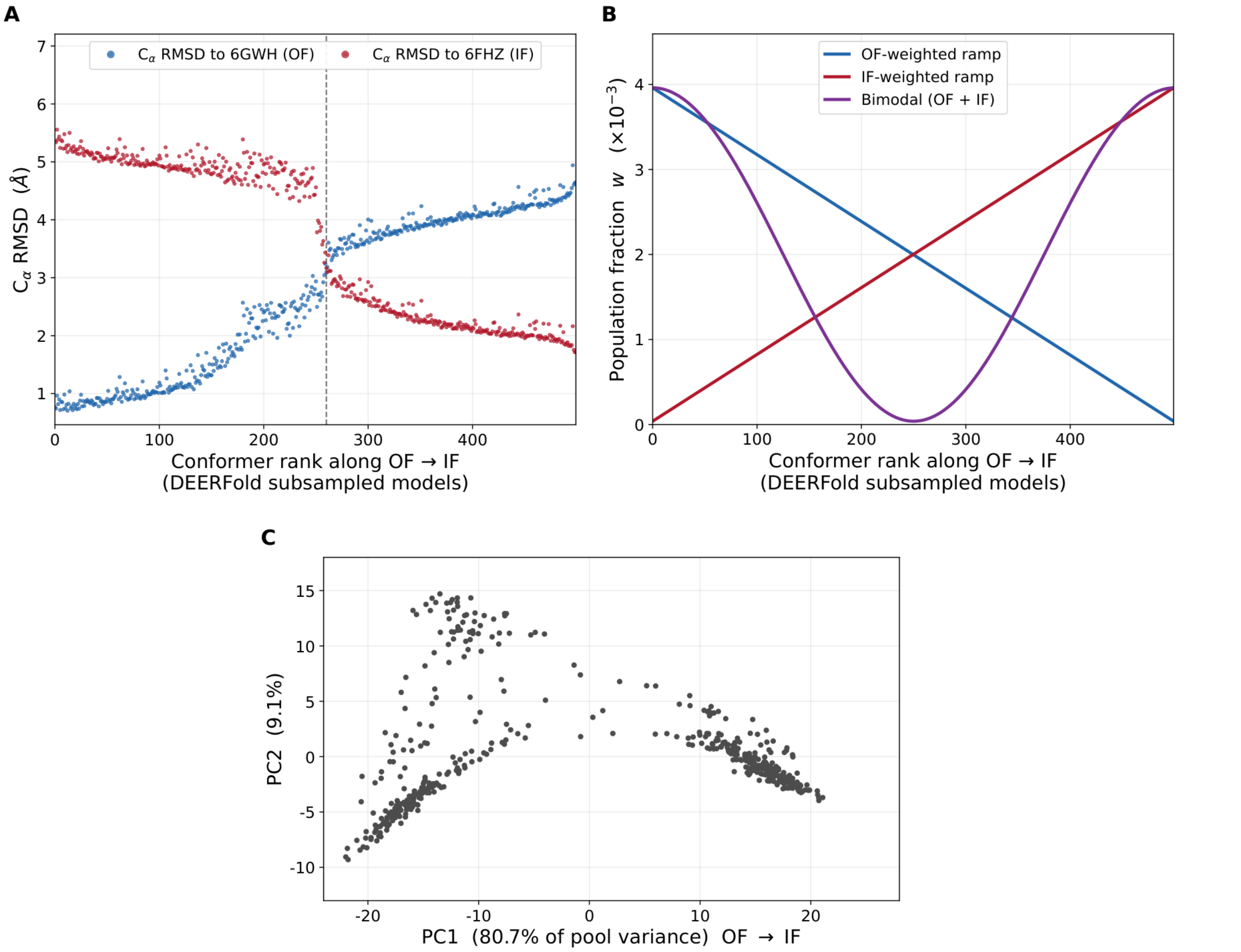


**Supplementary Fig. 5. Construction of the ground-truth ensembles for the synthetic recovery benchmark, and the conformation pool in the plane of Fig. 2B.** The pool consists of 500 PfMATE models generated with DEERFold[5] by distance restrained subsampling at $N_{eff}$ = 5. (A) $C_\alpha$ RMSD of each of the 500 models to the outward-facing (OF, 6GWH) and inward-facing (IF, 6FHZ) crystal structures. Models are ordered by $\Delta_{RMSD}$ = RMSD(6GWH) − RMSD(6FHZ). The two curves cross at rank 260 ($\Delta_{RMSD}$ = 0, dashed line). Panels A and B share the horizontal axis, the model rank from OF to IF. (B) The three population profiles imposed on this order, each normalized to sum to 1: the OF-weighted and IF-weighted ramps, linear in rank, and the bimodal OF + IF profile, a raised cosine in rank. (C) The same 500 models projected onto the plane of Fig. 2B. Each model is one point at its projection on PC1 and PC2 of the pool (80.7% and 9.1% of pool variance), with the same color and size for all points. Unlike Fig. 2B, this panel encodes no imposed population weight, because the weights are defined on rank (B), not on PC1. PC1 runs from outward-facing (6GWH) to inward-facing (6FHZ). Axis ranges are identical to those in the left column of Supplementary Fig. 6.

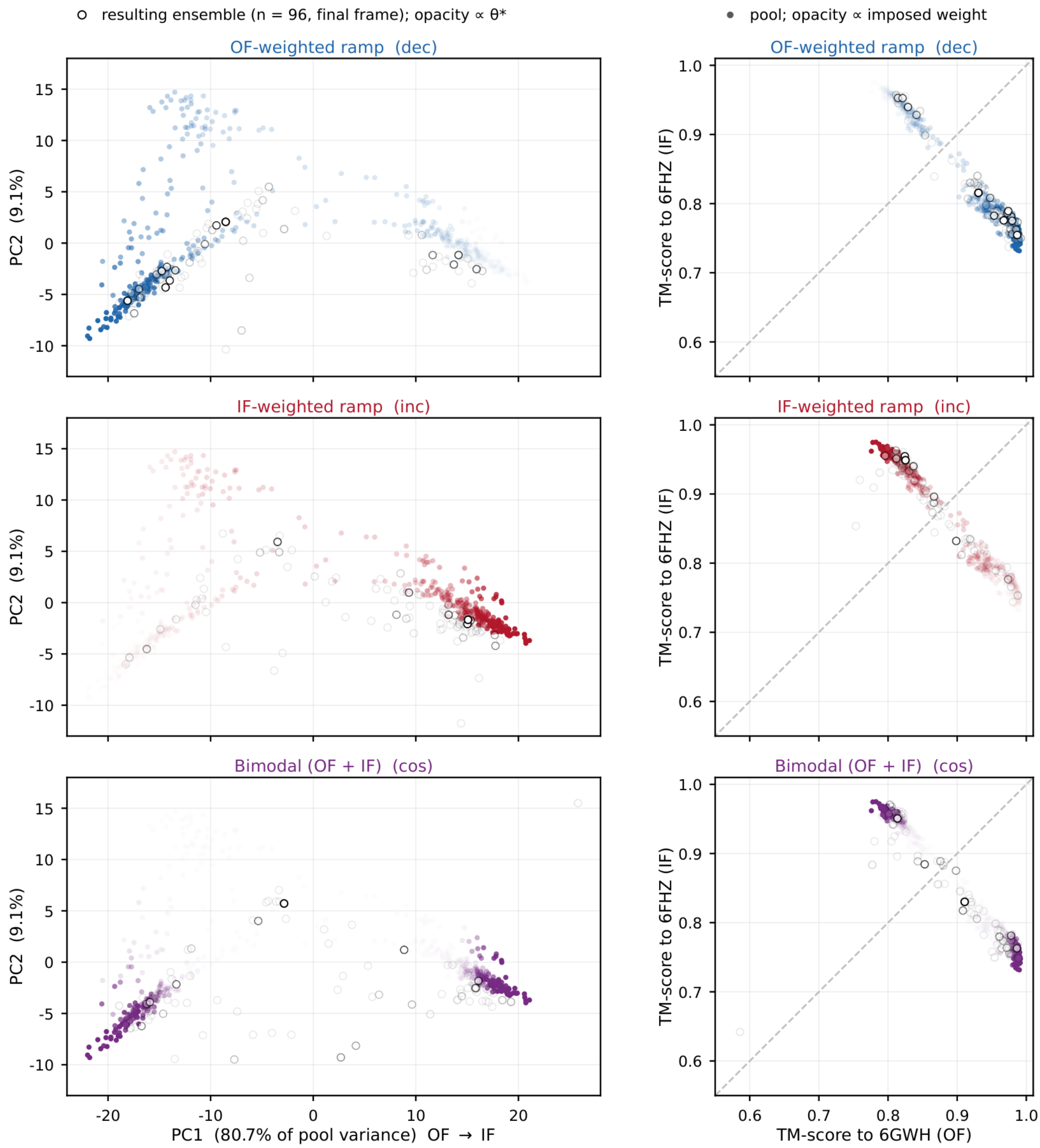


**Supplementary Fig. 6. θ*-weighted resulting ensembles of the synthetic runs in the pool plane of Fig. 2B (left) and in the plane of TM-scores to 6GWH and 6FHZ (right).** Left, open circles: the 96 structures of the resulting ensembles of the OF-weighted (top), IF-weighted (middle) and bimodal (bottom) runs on PC1 and PC2 of the 500-model pool, with opacity proportional to $\theta^*$. Along PC1 (PC units), the $\theta^*$-weighted means of the resulting ensembles versus the weighted target means are −7.06 versus −8.18 (OF-weighted), +7.36 versus +8.18 (IF-weighted) and −0.34 versus −0.003 (bimodal). Panel tags dec, inc and cos denote the OF-weighted, IF-weighted and bimodal targets. Right, the same structures and weights in the plane of TM-scores to 6GWH (OF, *x*) and 6FHZ (IF, *y*), as in Fig. 3A. Axes span 0.55–1.01, the gray dashed line marks the diagonal, and points are drawn as on the left. None of the 288 structures of the resulting ensembles has a minimum TM-score below 0.55, the distortion threshold of Fig. 3A.

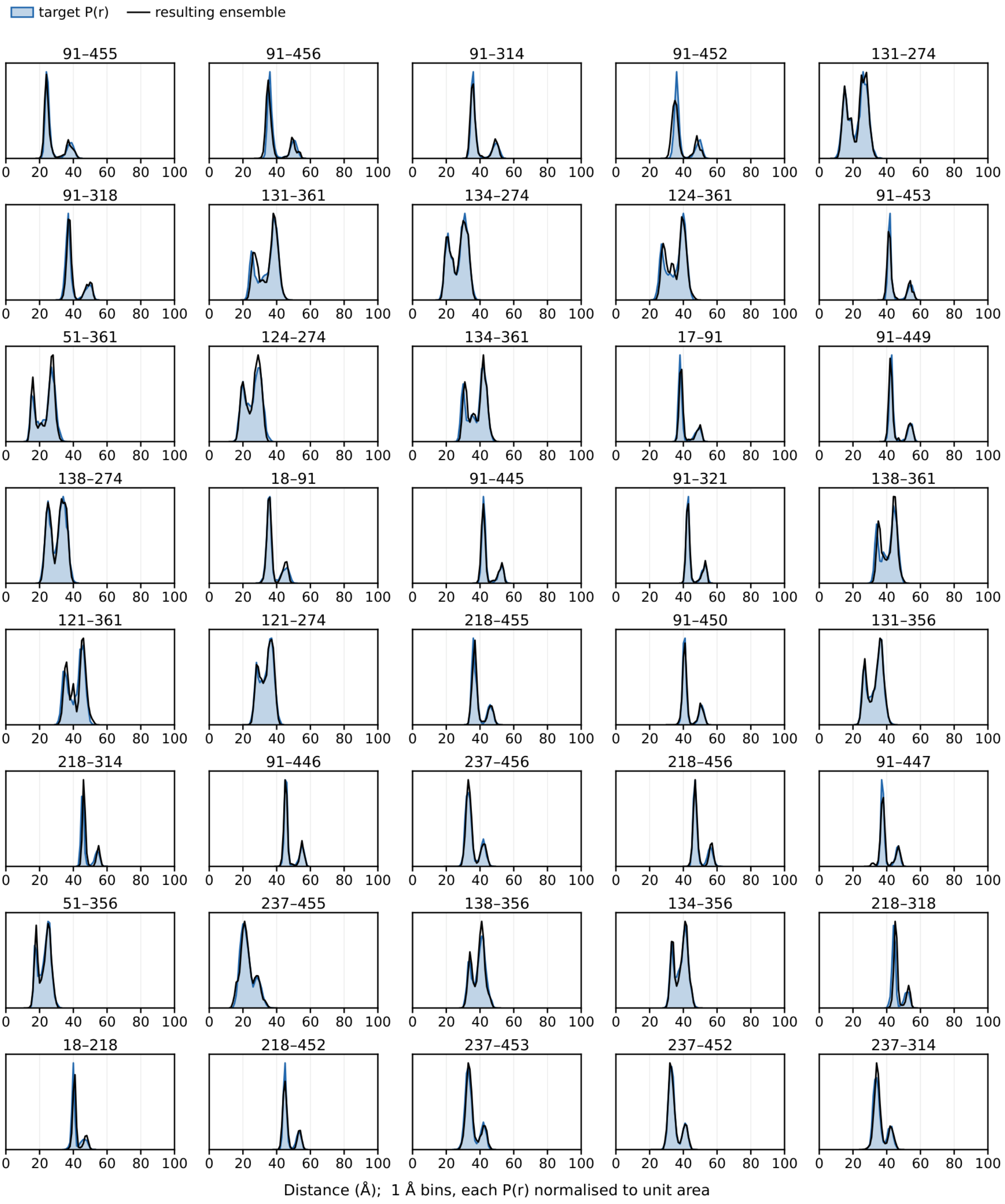

target P(r)
resulting ensemble
91–455
91–456
91–314
91–452
131–274
91–318
131–361
134–274
124–361
91–453
51–361
124–274
134–361
17–91
91–449
138–274
18–91
91–445
91–321
138–361
121–361
121–274
218–455
91–450
131–356
218–314
91–446
237–456
218–456
91–447
51–356
237–455
138–356
134–356
218–318
18–218
218–452
237–453
237–452
237–314
0 20 40 60 80 100
Distance (Å); 1 Å bins, each P(r) normalised to unit area

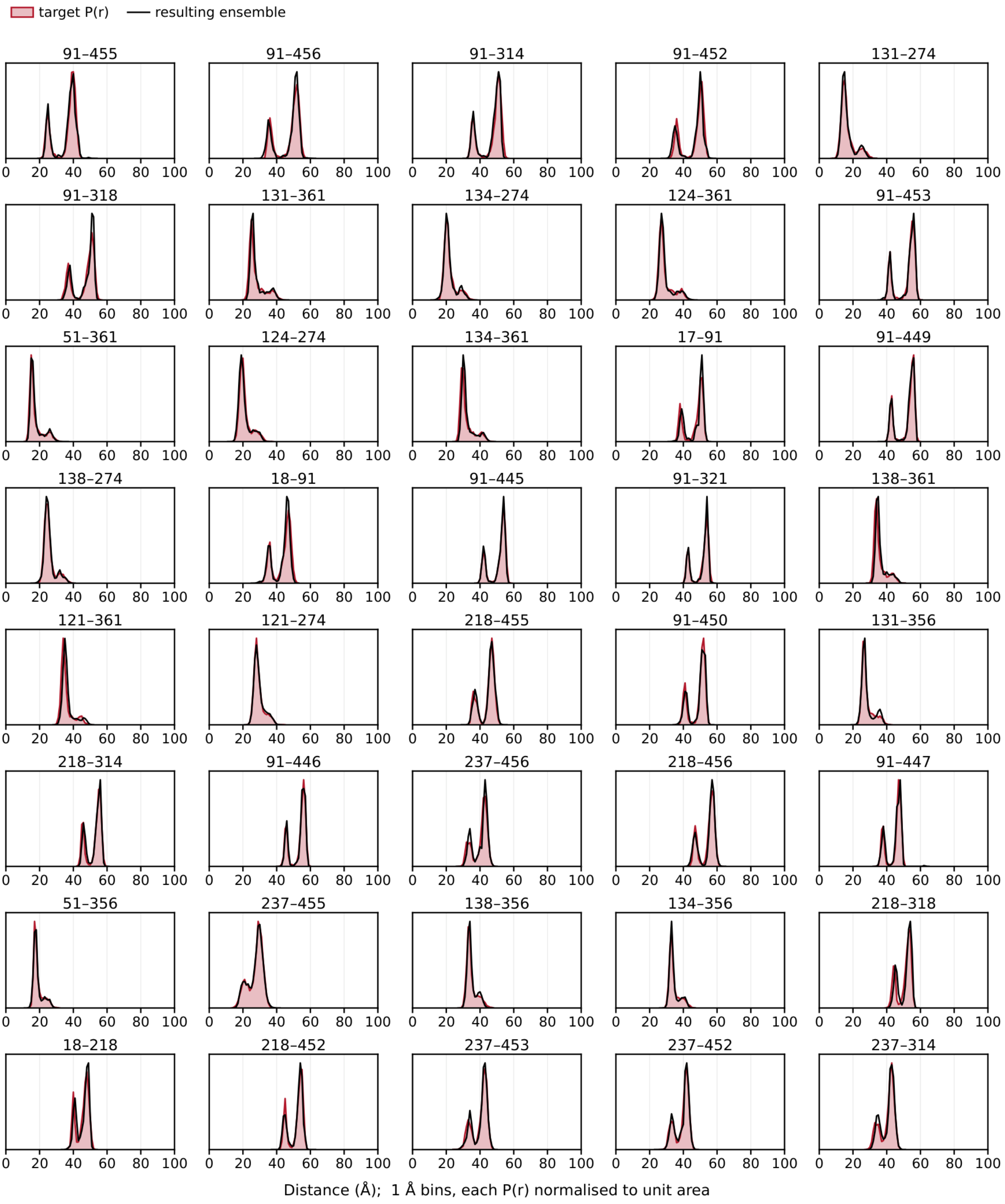
target P(r)
resulting ensemble
91–455
91–456
91–314
91–452
131–274
91–318
131–361
134–274
124–361
91–453
51–361
124–274
134–361
17–91
91–449
138–274
18–91
91–445
91–321
138–361
121–361
121–274
218–455
91–450
131–356
218–314
91–446
237–456
218–456
91–447
51–356
237–455
138–356
134–356
218–318
18–218
218–452
237–453
237–452
237–314
0 20 40 60 80 100
Distance (Å); 1 Å bins, each P(r) normalised to unit area

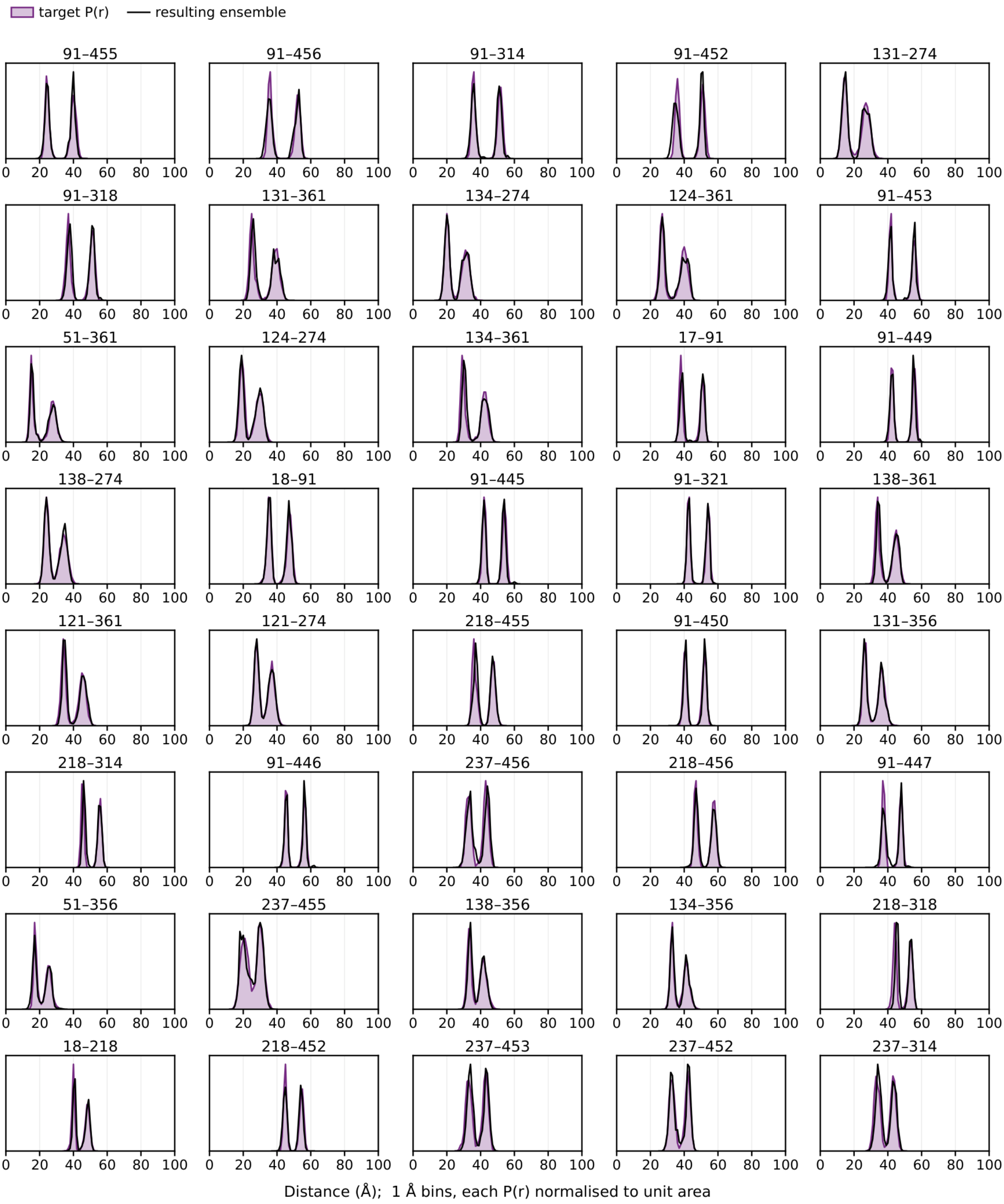


**Supplementary Fig. 7. Distance distributions of the three synthetic targets, with the resulting ensembles overlaid.** All 40 pairs of the OF-weighted ramp (blue), the IF-weighted ramp (red) and the bimodal target (purple) are shown, with the labeled residue pair in each panel title. Filled curves ("target *P*(*r*)"), in the colors of these targets in Fig. 2D, are the imposed target *P*(*r*) drawn from the weighted mixture over the 500-model pool. Black lines ("resulting ensemble") are the $\theta^*$-weighted *P*(*r*) of the resulting ensemble of the matching run, computed with environment atoms selected in each structure (*k* = 40 pairs, 96 structures of the final frame × $48^2$ spin-center samples, each distance carrying the weight $\theta_i^*/48^2$ of its structure *i*). The corresponding $W_1^*$ ($\theta^*$ weights) are 0.33 Å (OF-weighted), 0.28 Å (IF-weighted) and 0.35 Å (bimodal). All distributions are histogrammed in 1-Å bins, normalized to unit area and shown over 0–100 Å in every panel. Each panel is scaled to its own maximum, without a labeled y axis.

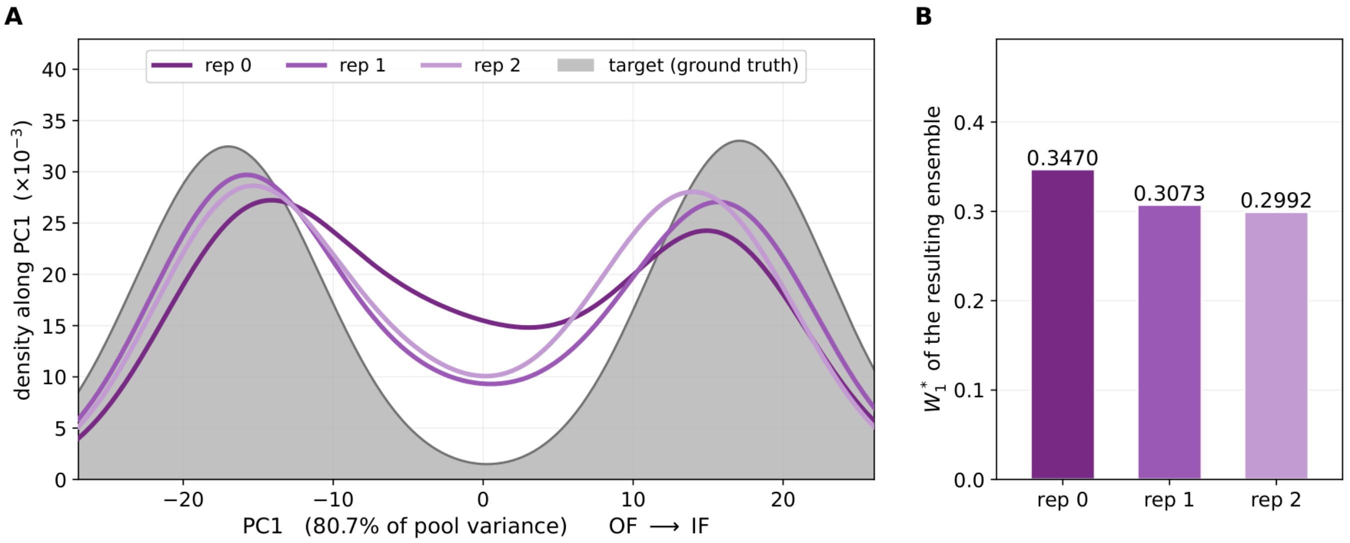


**Supplementary Fig. 8. Run-to-run reproducibility of the bimodal recovery.** Three runs (rep 0–2) were guided toward the same synthetic bimodal target with identical settings (40 DEER pairs) and differed only in the random seeds of Boltz-2 and of SC diffusion. (A) Density of each resulting ensemble (unweighted, 96 structures each) projected on the same PC1 axis of the 500-model pool (80.7% of pool variance, common Gaussian kernel width 5.73 PC units), with the ground-truth target in gray. (B) $W_1^*$ of the resulting ensemble of each run (offline $\theta^*$): 0.3470, 0.3073 and 0.2992 Å (mean 0.3178 Å, s.d. 0.0256 Å, i.e., 8.1% of the mean). This run-to-run spread serves as the resolution of $W_1^*$ comparisons. However, it was measured for one synthetic target and does not constitute a general noise floor.

A

B

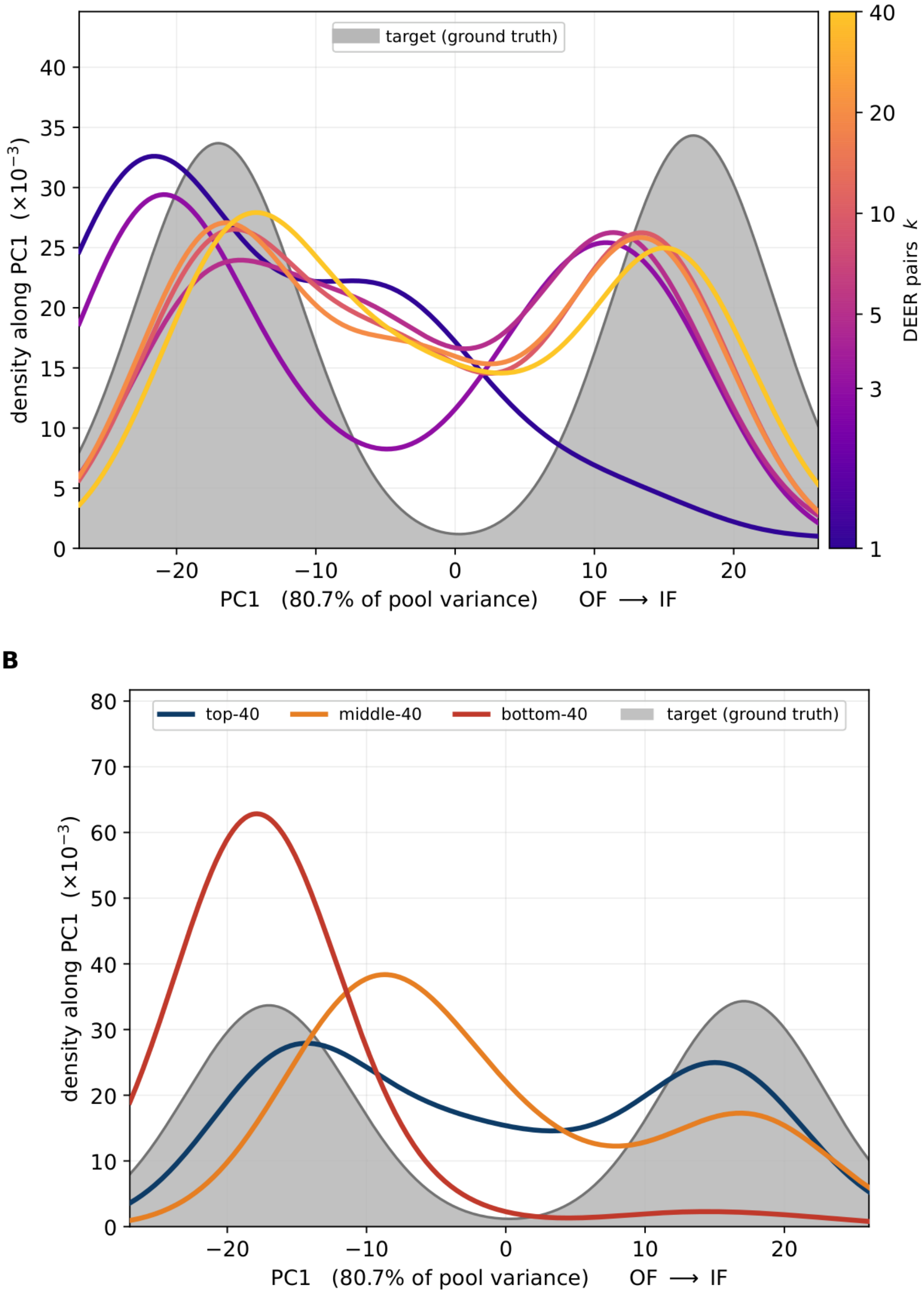


**Supplementary Fig. 9. Pair choice is more important than pair count.** Curves are densities of the resulting ensembles along PC1 of the 500-model pool (80.7% of pool variance), unweighted (96 structures each), with the ground-truth bimodal target in gray. (A) Pair count tests, from nested sets of the $k$ pairs with the largest $|\Delta d|$ ($k$ = 1, 3, 5, 10, 20 and 40, colored by $k$). (B) Pair choice tests, from three different sets of 40 pairs (top, middle and bottom 40 by $|\Delta d|$). $|\Delta d|$ is the difference in mean predicted spin-center distance of a pair between the reference OF and IF structures. Because each $k$ is represented by a single nested subset, the number of pairs and the number of labeled sites increase together. The top-40 set in B is the same run as $k$ = 40 in A and as rep 0 in Supplementary Fig. 8.

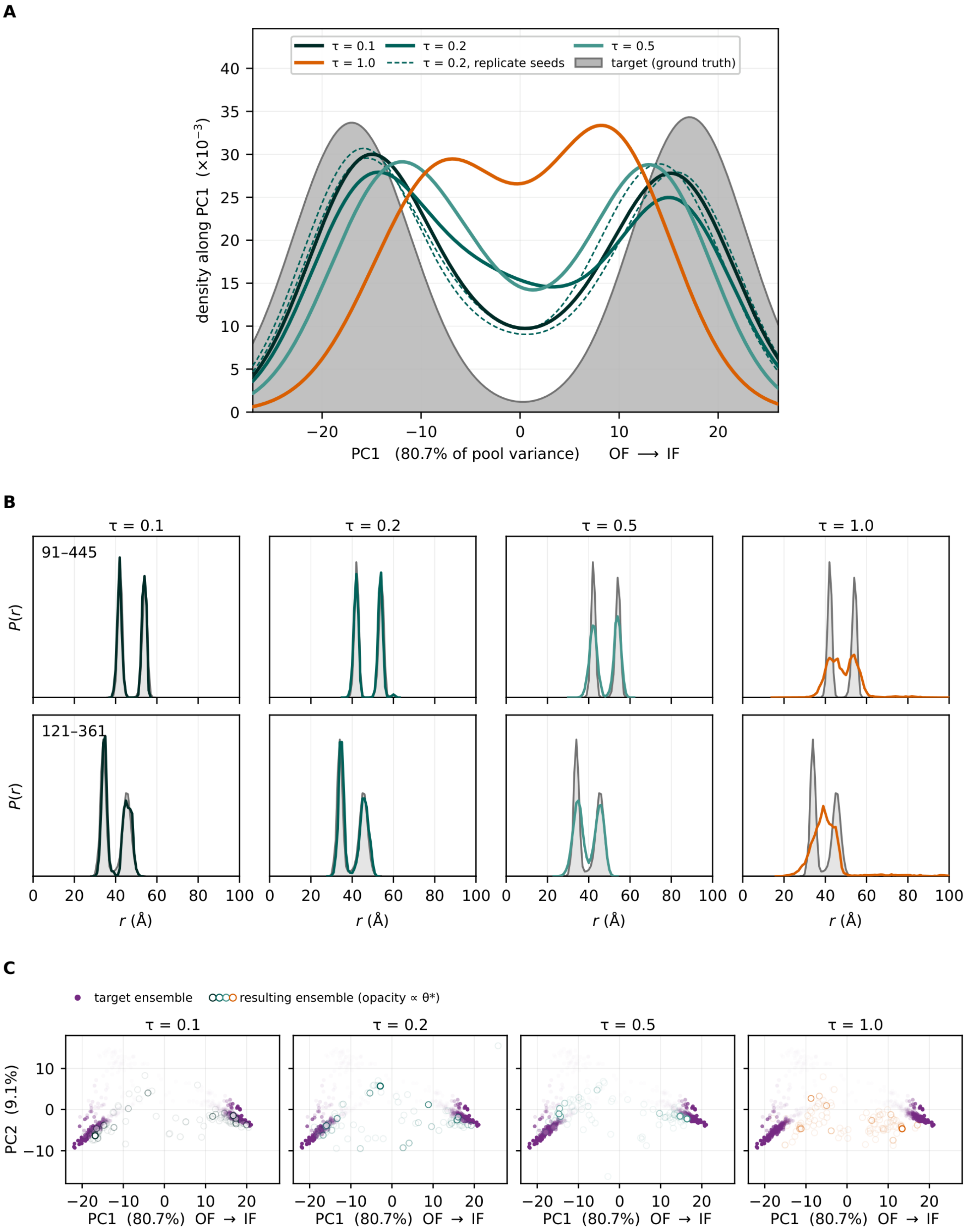


**Supplementary Fig. 10. Recovery of the bimodal synthetic target with different SC diffusion sampling temperatures τ during guidance.** The target (Fig. 2, 40 DEER pairs) was generated at τ = 0.2, whereas guidance used τ = 0.1, 0.2, 0.5 or 1.0. The four runs ($N$ = 96 structures) are otherwise identical, including the random seed. (A) Unweighted densities along pool PC1 (kernel width 5.49 PC units): target (gray), resulting ensembles (solid, colored by τ) and reps 1 and 2 of Supplementary Fig. 8 (thin dashed). (B) Distance distributions of pairs 91–445 (top) and 121–361 (bottom) of Fig. 2C, with one column per τ and one vertical scale per row:

target (gray) and $\theta^*$-weighted resulting ensemble (colored). (C) Pool plane of Fig. 2B, one panel per $\tau$ on shared axes: 500 pool models (purple, opacity by target weight $w$) and structures of the resulting ensembles (open circles colored as in A, opacity by $\theta^*$ as in Supplementary Fig. 6). $W_1^*$ of the resulting ensembles (in-sample) are 0.3434, 0.3470, 0.4041 and 1.2940 Å for $\tau = 0.1$, 0.2, 0.5 and 1.0, respectively. The runs ended after 2–5 guidance cycles.

**A**

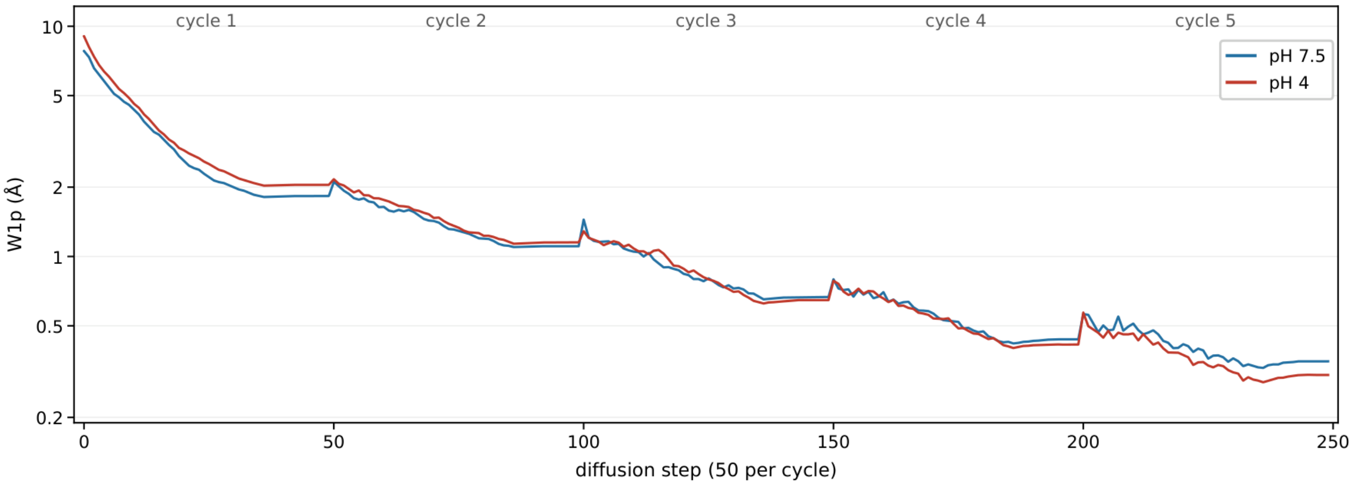

cycle 1
cycle 2
cycle 3
cycle 4
cycle 5
pH 7.5
pH 4
W1p (Å)
10
5
2
1
0.5
0.2
0
50
100
150
200
250
diffusion step (50 per cycle)

**B**

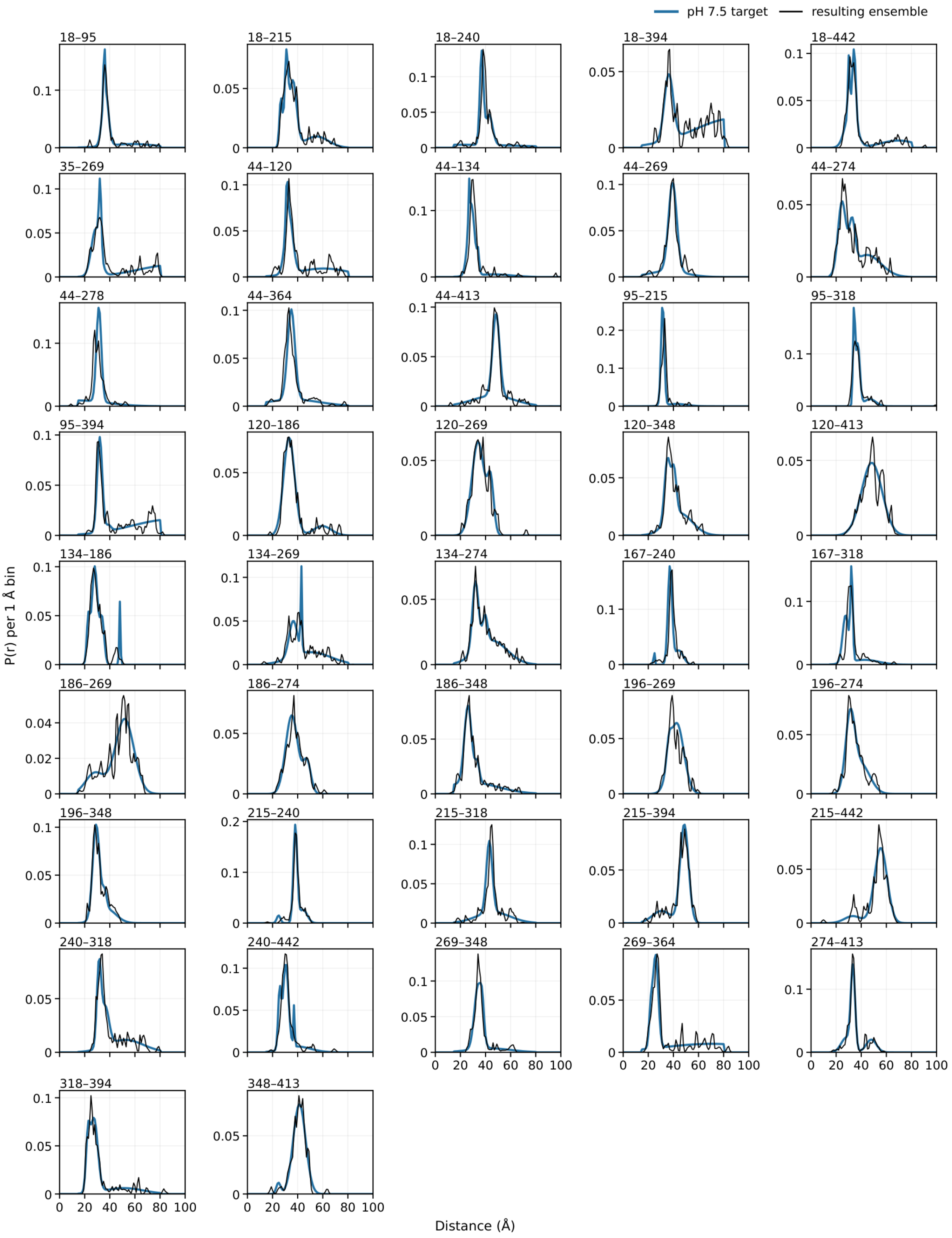
pH 7.5 target
resulting ensemble
18–95
18–215
18–240
18–394
18–442
35–269
44–120
44–134
44–269
44–274
44–278
44–364
44–413
95–215
95–318
95–394
120–186
120–269
120–348
120–413
134–186
134–269
134–274
167–240
167–318
186–269
186–274
186–348
196–269
196–274
196–348
215–240
215–318
215–394
215–442
240–318
240–442
269–348
269–364
274–413
318–394
348–413
P(r) per 1 Å bin
Distance (Å)

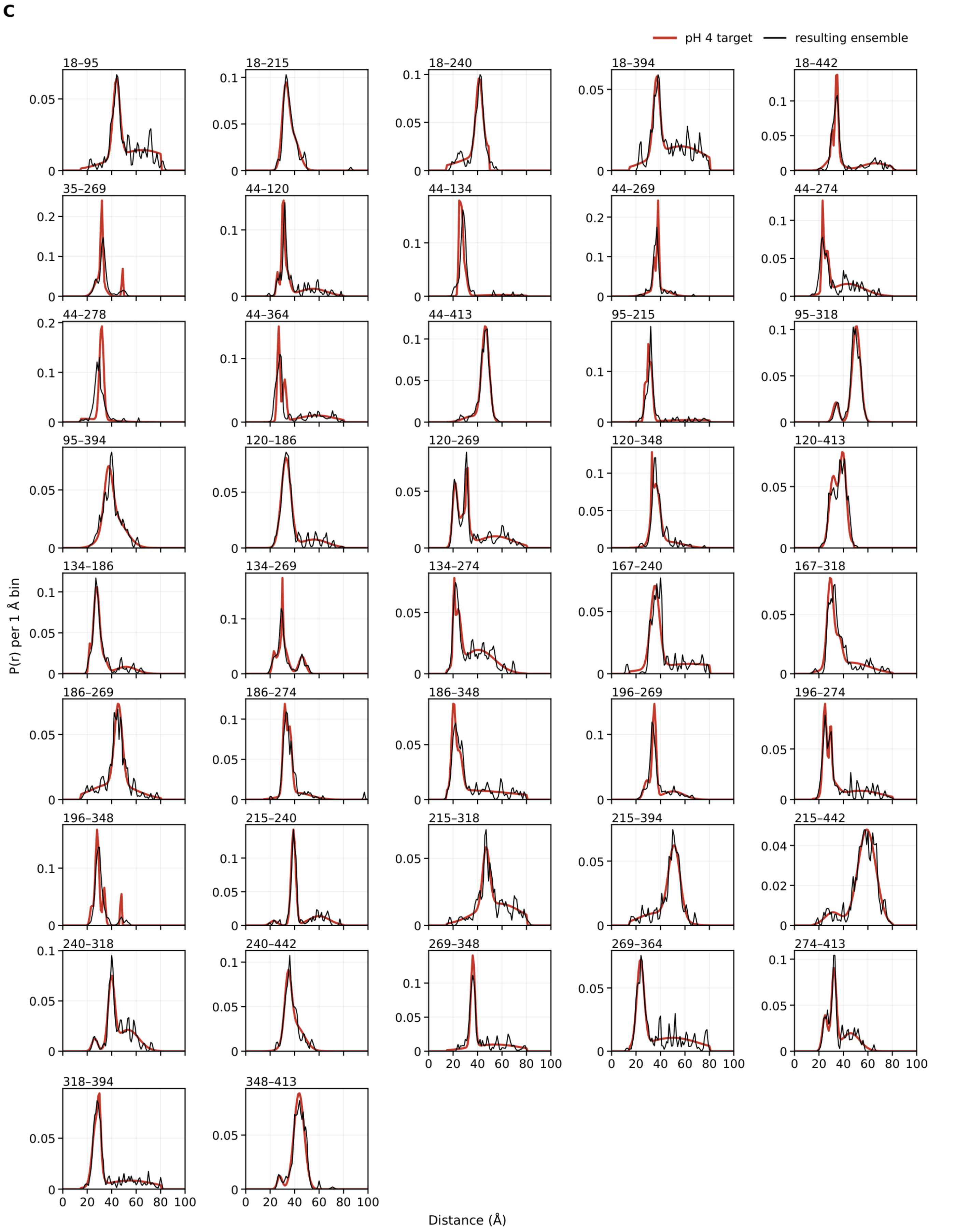


**Supplementary Fig. 11. PfMATE: in-run $W_1^p$ trajectories, experimental targets and distance distributions of the resulting ensembles for the two de novo runs of Fig. 3A.** The two runs (pH 7.5 in blue, pH 4 in red) each used $N = 96$ structures and five guidance cycles, without early stop. (A) $W_1^p$ of the denoised estimate $\hat{x}_0$ at every step of the 50-step cycles, drawn as one continuous line. The jump at each cycle start reflects the restart of denoising from noise. Step 49 of the last cycle is the resulting ensemble. $W_1^p$

is pair-weighted over the 42 pairs ($W_1^p = \sum_k W_{1,k}^2 / \sum_k W_{1,k}$, uniform model weights) and uses environment atoms fixed from a reference model. It is therefore not comparable with $W_1^*$ of the resulting ensembles. (B) pH 7.5 and (C) pH 4: experimental DEER distance distributions[3] used as targets (colored line) for all 42 pairs, on their native 1-Å grid. Black line: distribution predicted from the resulting ensemble with environment atoms selected in each structure, with each of the $48^2$ spin-center distances of structure $i$ weighted by $\theta_i^*/48^2$, in 1-Å bins of unit total probability. $W_1^*$ of the resulting ensembles are 0.71 Å at pH 7.5 and 0.71 Å at pH 4. Curves connect the bin values on a common 0–100 Å axis, with pairs in residue order.

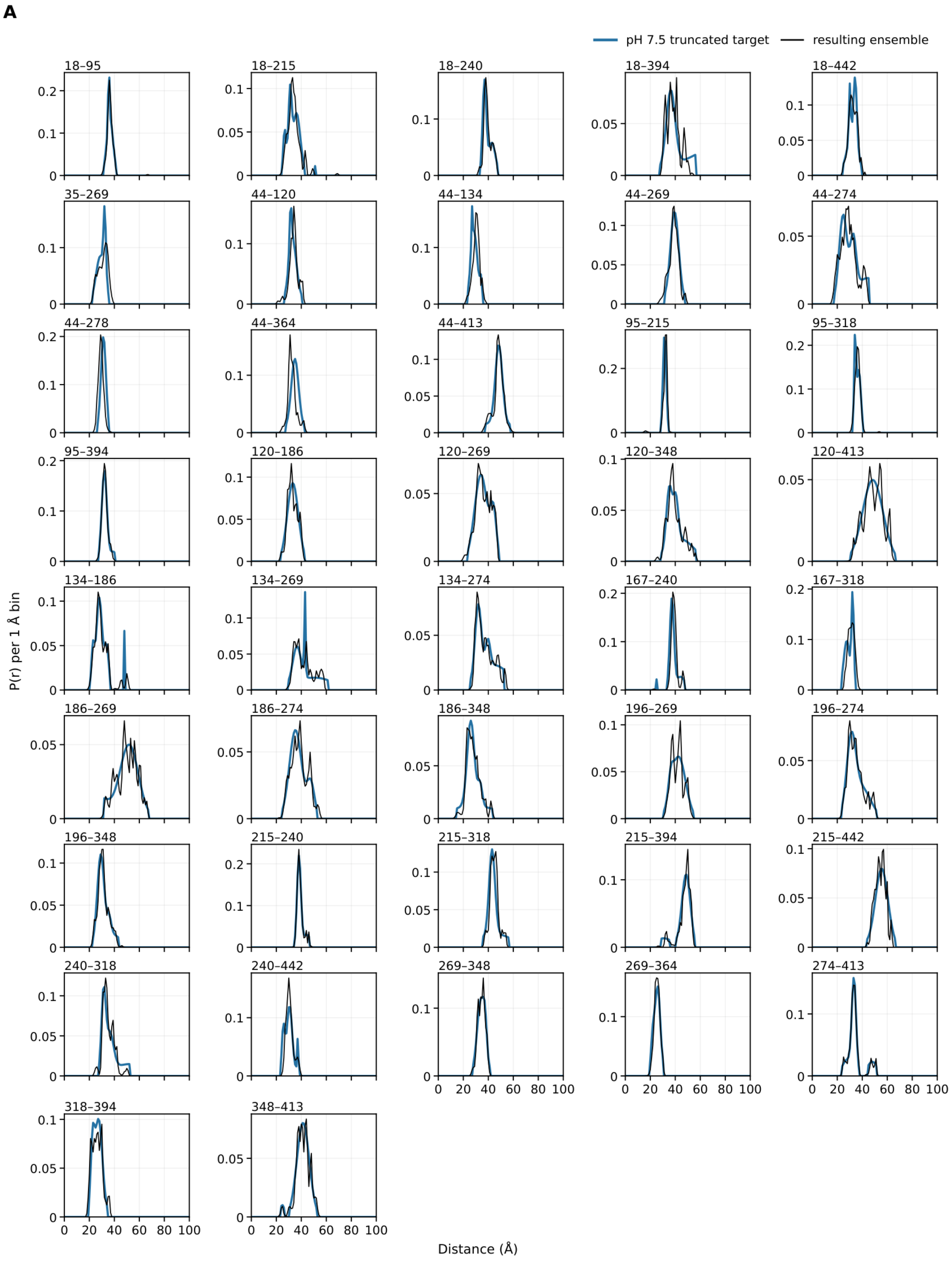
A
pH 7.5 truncated target
resulting ensemble
18–95
18–215
18–240
18–394
18–442
35–269
44–120
44–134
44–269
44–274
44–278
44–364
44–413
95–215
95–318
95–394
120–186
120–269
120–348
120–413
134–186
134–269
134–274
167–240
167–318
186–269
186–274
186–348
196–269
196–274
196–348
215–240
215–318
215–394
215–442
240–318
240–442
269–348
269–364
274–413
318–394
348–413
P(r) per 1 Å bin
Distance (Å)

**B**

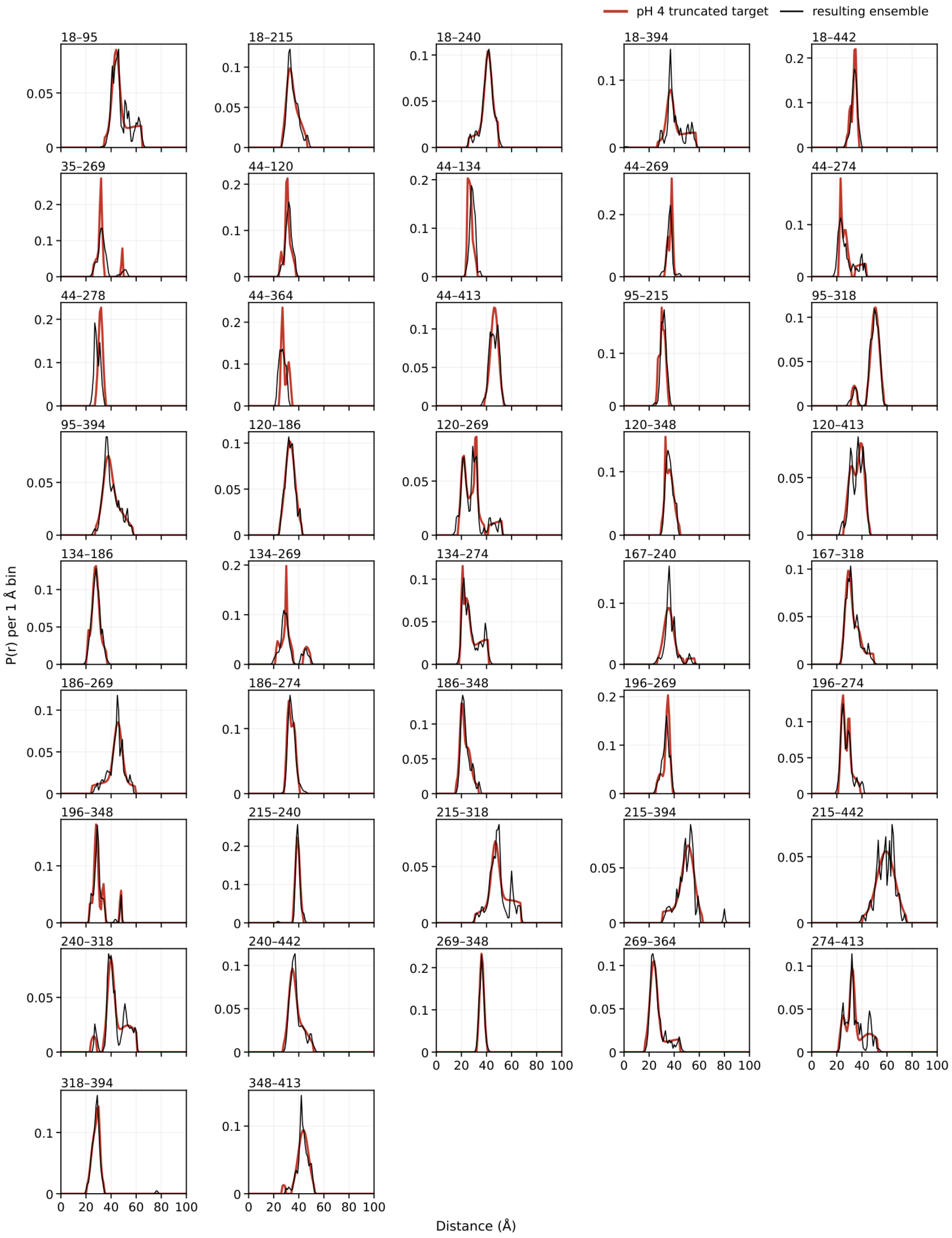


**Supplementary Fig. 12. PfMATE runs of Fig. 3B: tail-truncated DEER targets and predictions from the resulting ensembles.** (A) pH 7.5 and (B) pH 4, all 42 PfMATE pairs in residue order. Colored line: the tail-truncated target used by the run. Truncation retains the bins of the Fig. 3A target[3] (Supplementary Fig. 11) that fall within ±20 Å of its mode and whose probability is at least 10% of the peak value, then renormalizes the retained bins to unit mass. The truncated targets are therefore higher than the originals inside the retained window. Black line: distribution predicted from the $\theta^*$-weighted resulting ensemble of the run (the 96 structures of the last cycle), drawn as in Supplementary Fig. 11B,C. $W_1^*$ of the resulting ensembles are 0.68 Å at pH 7.5 and 0.64 Å at pH 4. Because these values are measured against the truncated targets, they are not comparable with the $W_1^*$ of the Fig. 3A runs.

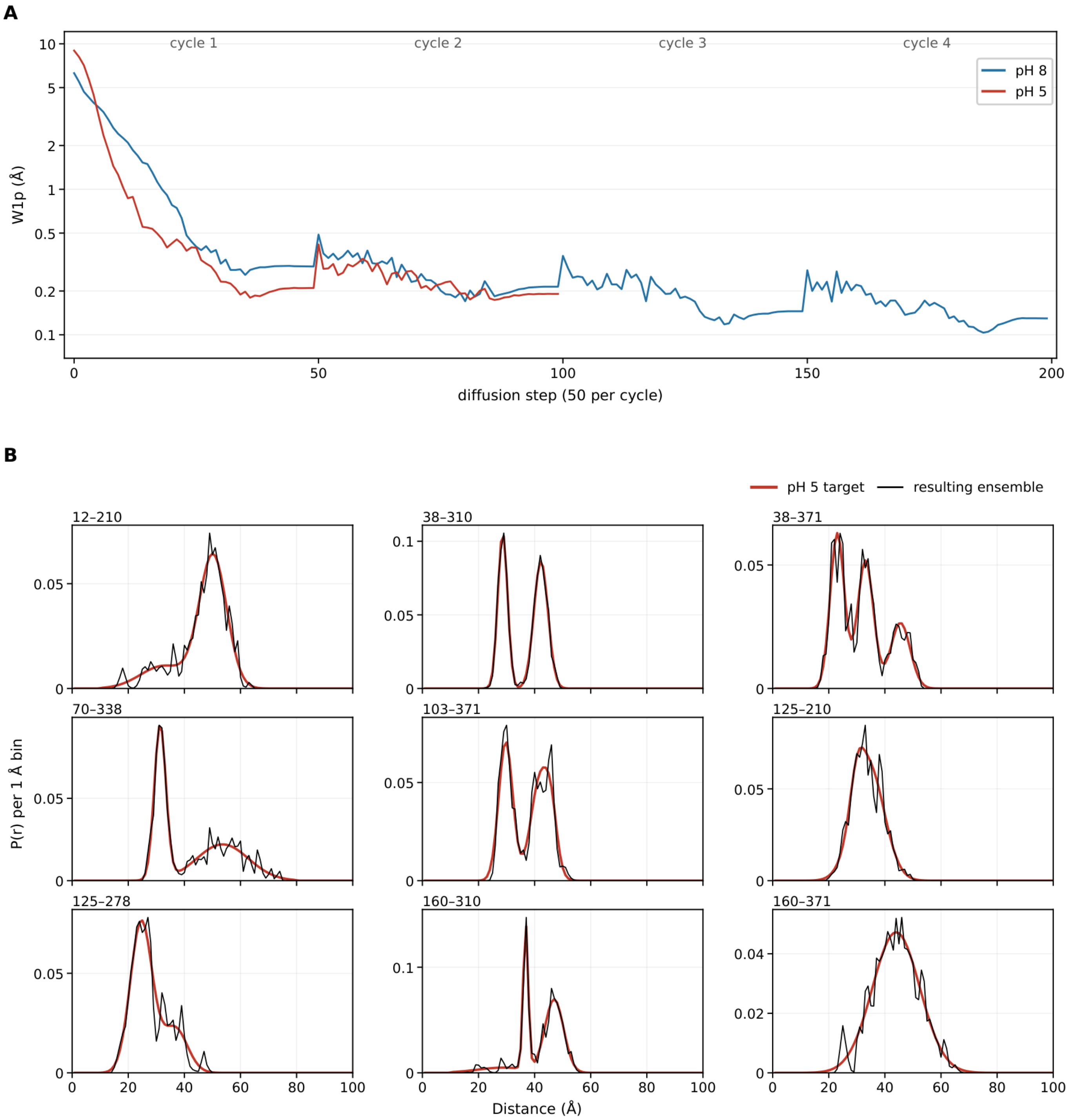
A
cycle 1
cycle 2
cycle 3
cycle 4
pH 8
pH 5
W1p (Å)
diffusion step (50 per cycle)
B
pH 5 target
resulting ensemble
12–210
38–310
38–371
70–338
103–371
125–210
125–278
160–310
160–371
P(r) per 1 Å bin
Distance (Å)

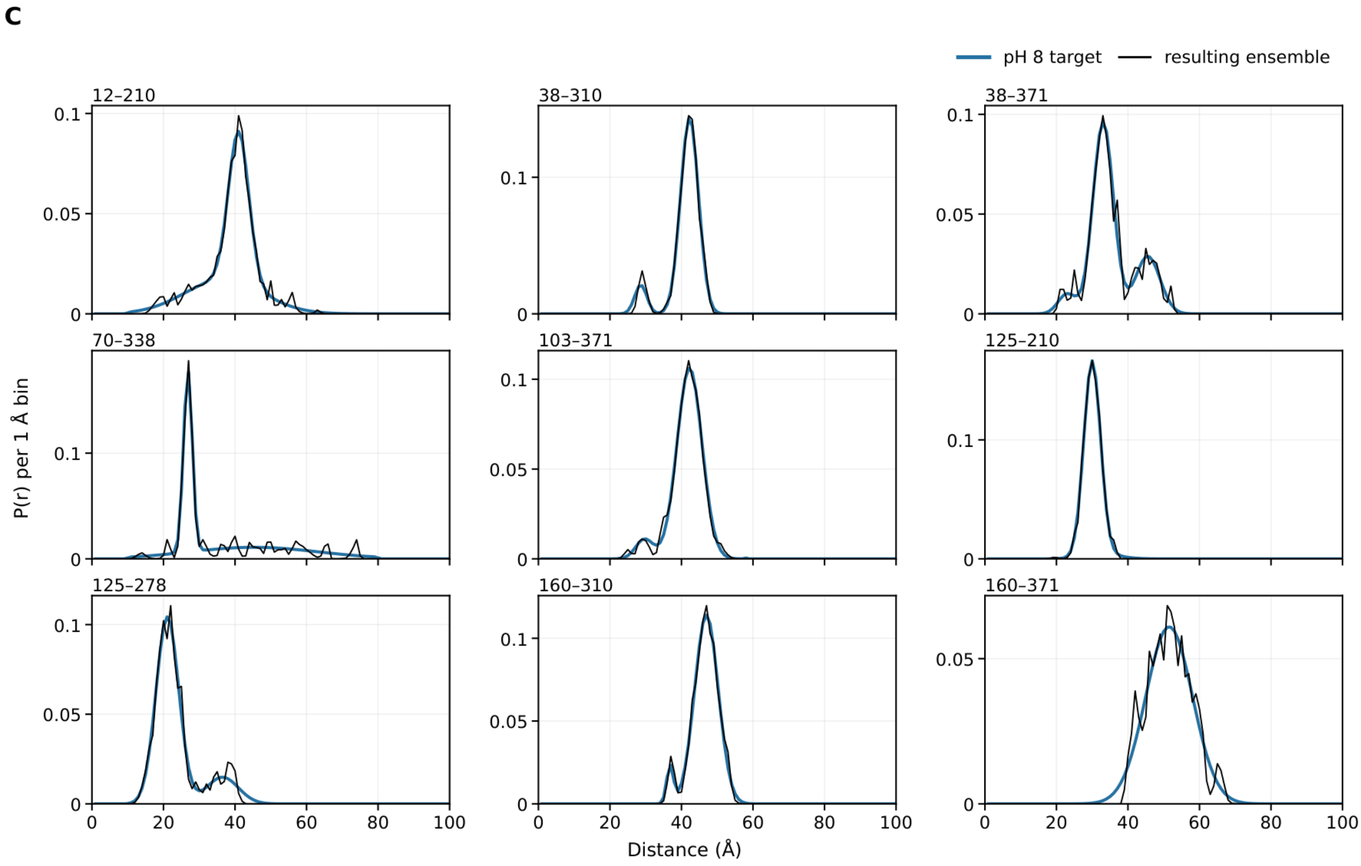


**Supplementary Fig. 13. LmrP: in-run $W_1^p$ trajectories, experimental targets and distance distributions of the resulting ensembles for the two de novo runs of Fig. 3D.** The two runs (pH 8 in blue, pH 5 in red) each used $N = 96$ structures. Panels are drawn as in Supplementary Fig. 11. (A) Both runs were stopped early (Methods), pH 5 after cycle 2 and pH 8 after cycle 4. (B) pH 5 and (C) pH 8: experimental DEER distance distributions[6,7] (colored line) of the nine pairs. $W_1^*$ of the resulting ensembles are 0.151 Å at pH 5 and 0.137 Å at pH 8.

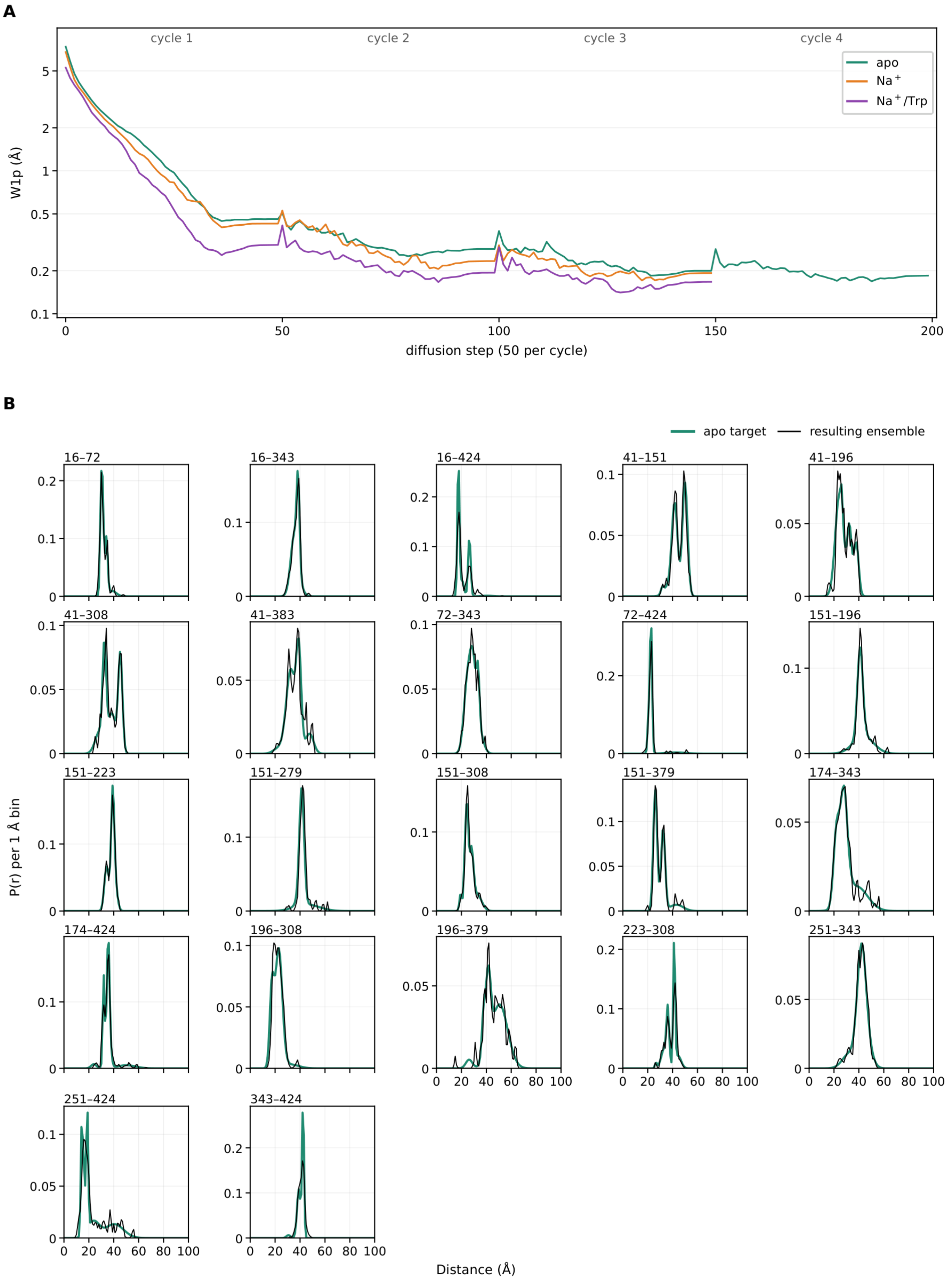
A
cycle 1
cycle 2
cycle 3
cycle 4
apo
Na$^+$
Na$^+$/Trp
W1p (Å)
diffusion step (50 per cycle)
B
apo target
resulting ensemble
16–72
16–343
16–424
41–151
41–196
41–308
41–383
72–343
72–424
151–196
151–223
151–279
151–308
151–379
174–343
174–424
196–308
196–379
223–308
251–343
251–424
343–424
P(r) per 1 Å bin
Distance (Å)

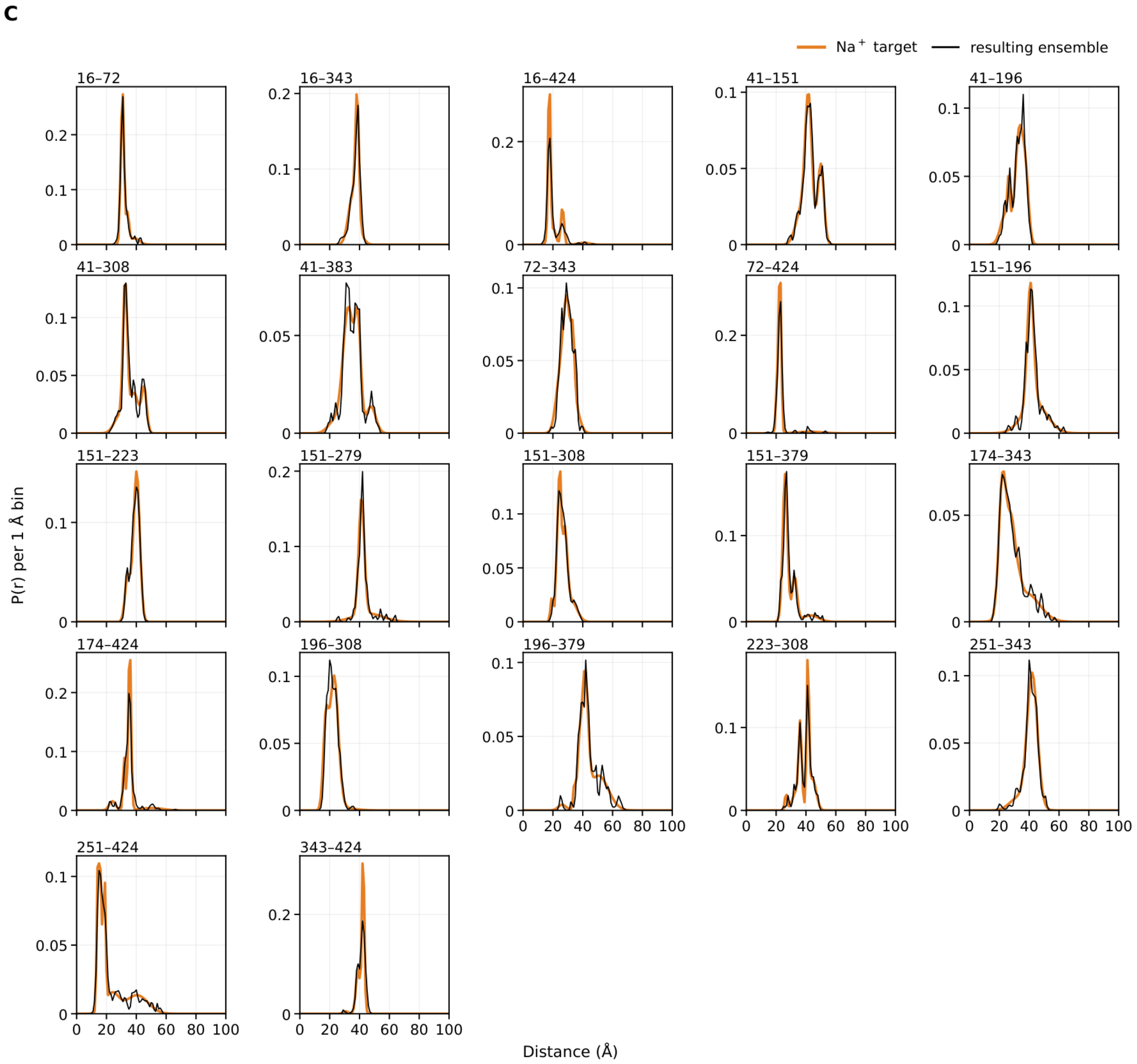
C
Na$^+$ target
resulting ensemble
16–72
16–343
16–424
41–151
41–196
41–308
41–383
72–343
72–424
151–196
151–223
151–279
151–308
151–379
174–343
174–424
196–308
196–379
223–308
251–343
251–424
343–424
P(r) per 1 Å bin
Distance (Å)

**D**

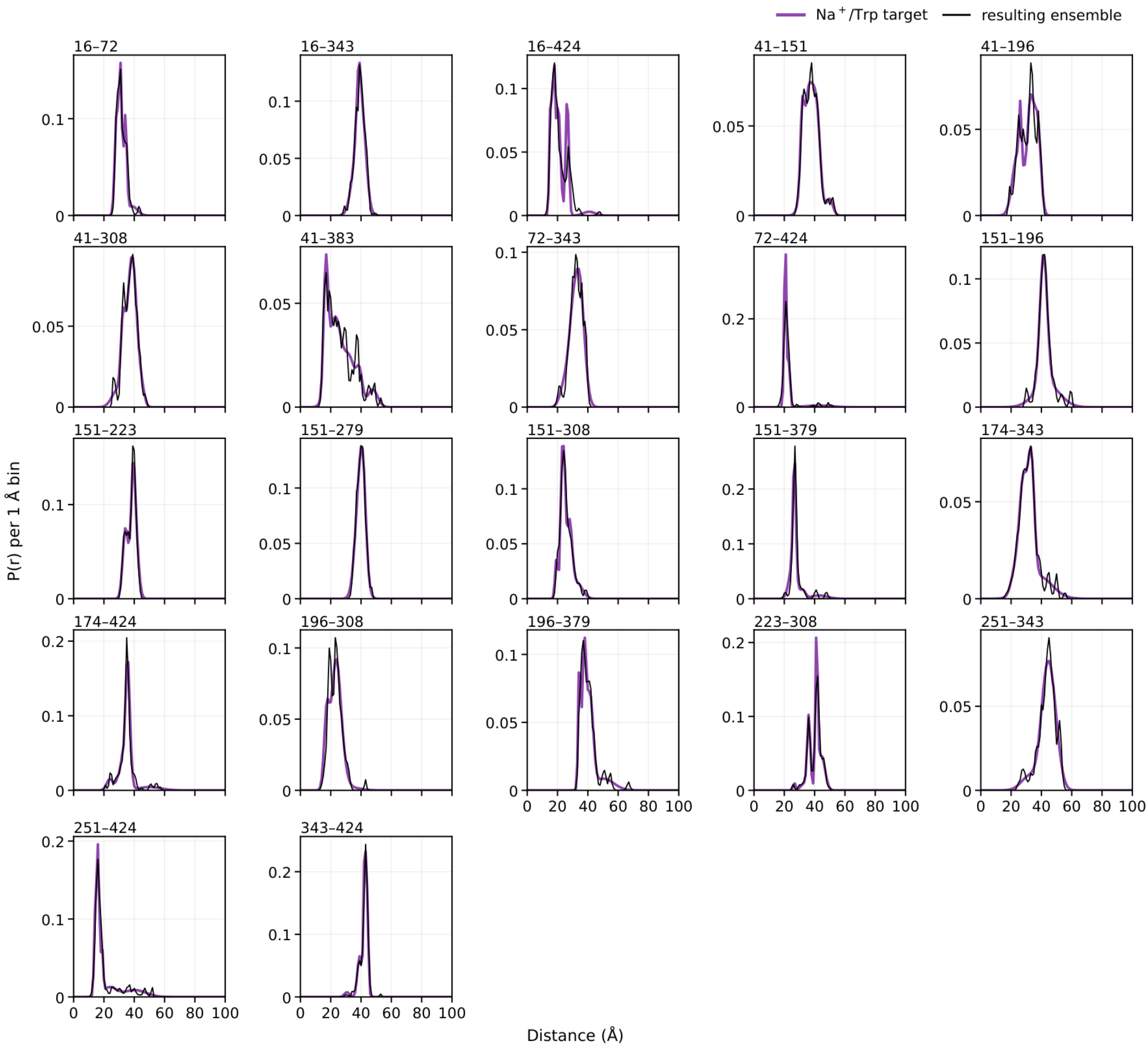


**Supplementary Fig. 14. MhsT, detergent (DM) data set: in-run $W_1^p$ trajectories, experimental targets and distance distributions of the resulting ensembles.** Shown are the three MhsT de novo runs of Fig. 3G, fitted to the DM DEER data set[8] (apo in green, $Na^+$ in orange and $Na^+$/Trp in purple, $N = 96$ structures each). Panels are drawn as in Supplementary Fig. 11. (A) All three runs were stopped early (Methods), apo after cycle 4 and the $Na^+$ and $Na^+$/Trp runs after cycle 3. (B) apo, (C) $Na^+$, (D) $Na^+$/Trp: experimental DEER distance distributions (colored line) of the 22 DM pairs. $W_1^*$ of the resulting ensembles are 0.348, 0.305 and 0.269 Å for apo, $Na^+$ and $Na^+$/Trp, respectively.

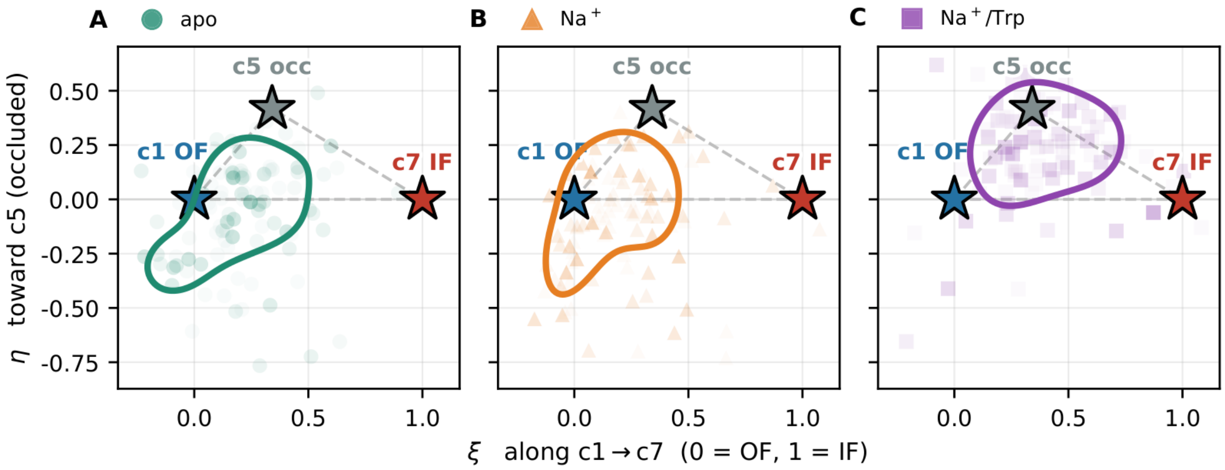


**Supplementary Fig. 15. MhsT ensembles fitted to the nanodisc (ND) data set on the ξ–η plane, one panel per condition.** MhsT de novo ensembles fitted to the ND DEER data set[8]: (A) apo, (B) $Na^+$ and (C) $Na^+$/Trp (96 structures each). The 20 ND pairs are a subset of the 22 pairs of the detergent (DM) data set of Fig. 3G. ξ is the projection along the c1 (OF) → c7 (IF) direction (0 = c1, 1 = c7). η is the orthogonal component toward c5 (occluded), in the same units (|c7 − c1| = 4.07 Å $C_\alpha$ RMSD, with c5 1.70 Å off the c1–c7 line). Stars mark the three reference models. Axis ranges, reference positions, density grid, colors and markers are identical to those of Fig. 3G. Each point represents one structure, with opacity scaled by its $\theta^*$ relative to the heaviest structure of its condition and a lower bound that renders low-weight structures visible. The colored line is a single contour of each condition's $\theta^*$-weighted kernel density at the 88th percentile of the density values on the shared grid, outlining the high-density core. $\theta^*$-weighted means (ξ, η): apo (0.21, −0.08), $Na^+$ (0.20, −0.09), $Na^+$/Trp (0.40, +0.20). Trajectories and per-pair fits of these runs are shown in Supplementary Fig. 16.

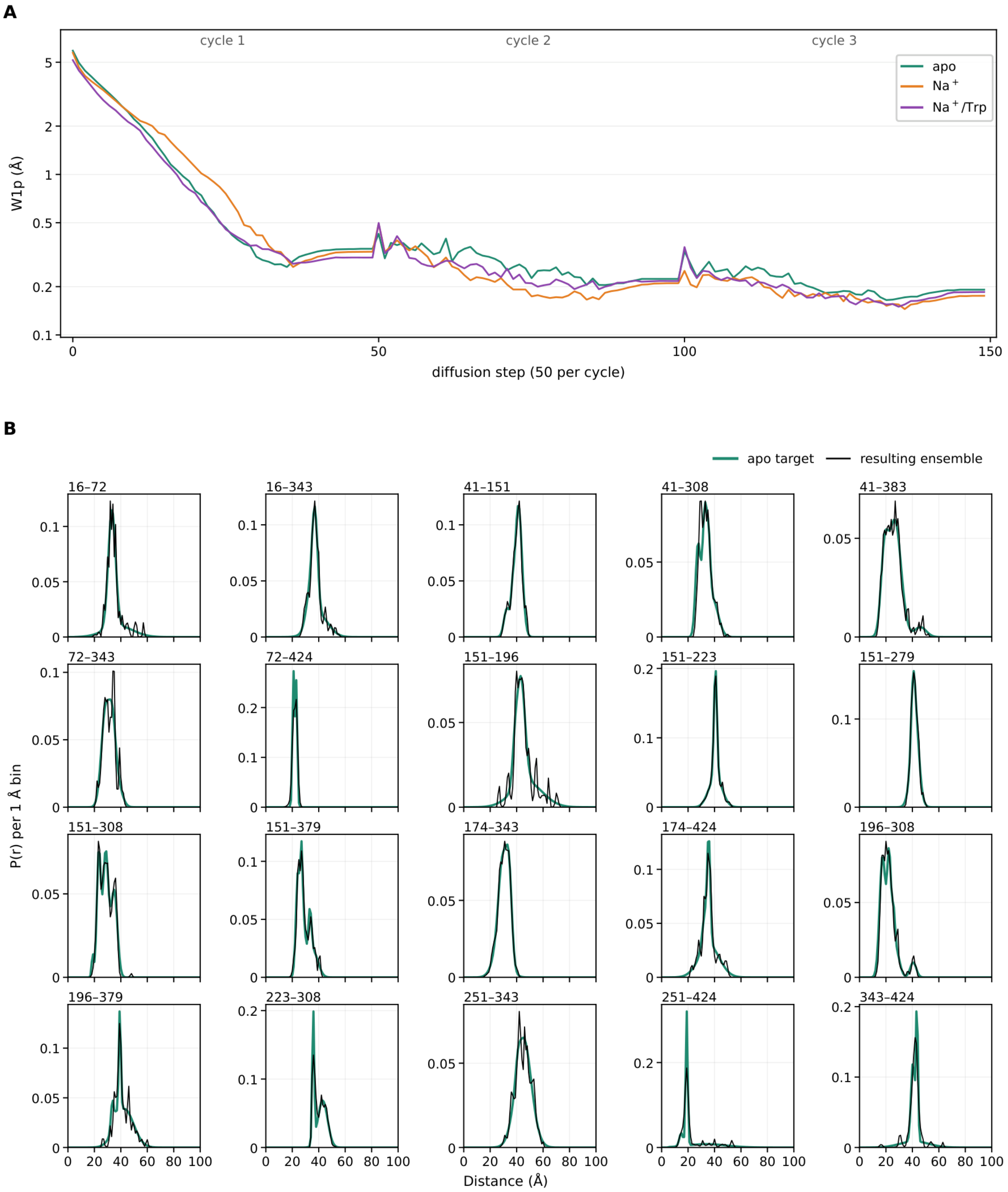

A
cycle 1
cycle 2
cycle 3
apo
Na$^+$
Na$^+$/Trp
W1p (Å)
diffusion step (50 per cycle)
B
apo target
resulting ensemble
16–72
16–343
41–151
41–308
41–383
72–343
72–424
151–196
151–223
151–279
151–308
151–379
174–343
174–424
196–308
196–379
223–308
251–343
251–424
343–424
P(r) per 1 Å bin
Distance (Å)

**C**

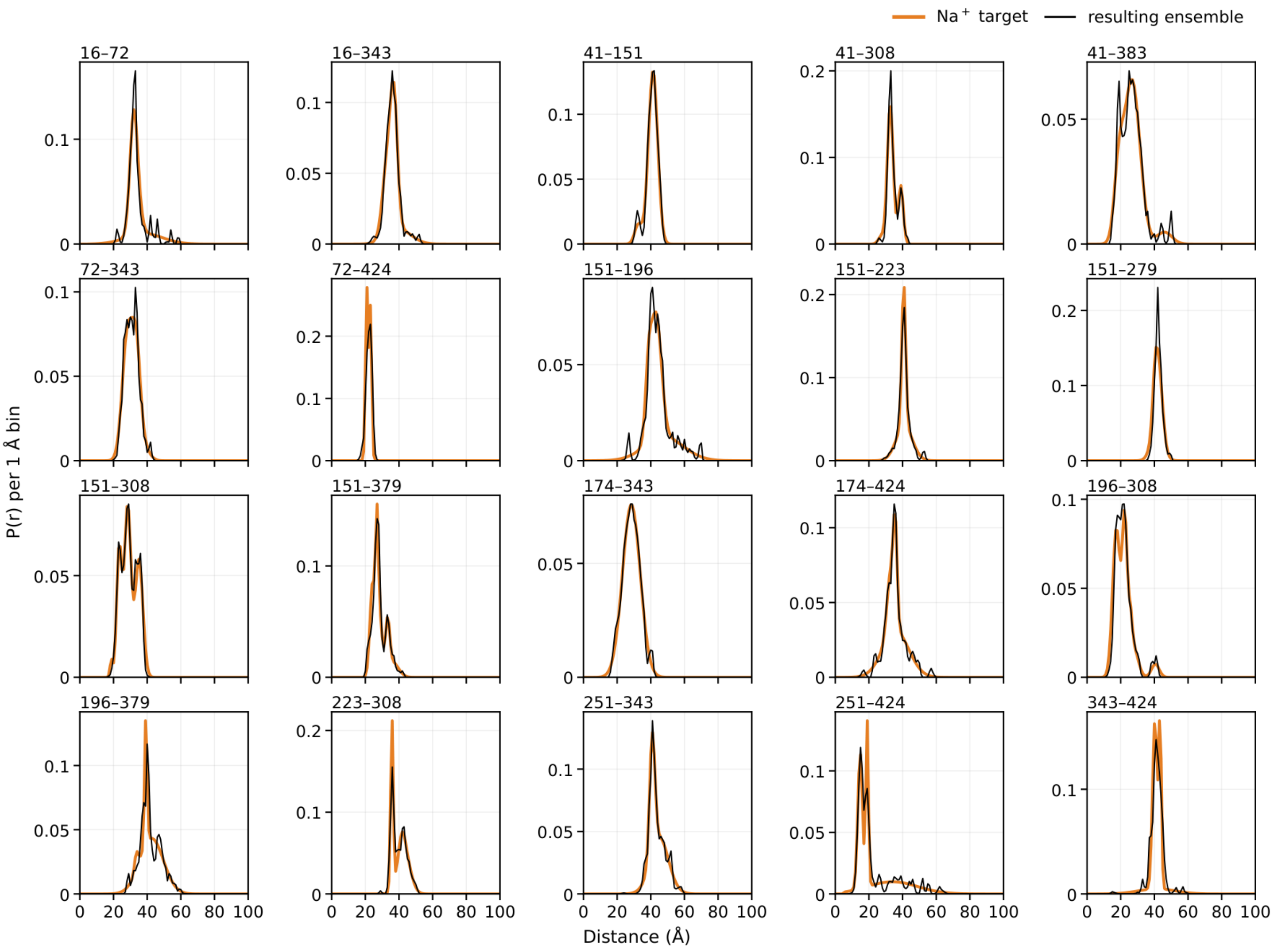
Na⁺ target
resulting ensemble
16–72
16–343
41–151
41–308
41–383
72–343
72–424
151–196
151–223
151–279
151–308
151–379
174–343
174–424
196–308
196–379
223–308
251–343
251–424
343–424
P(r) per 1 Å bin
Distance (Å)

**D**

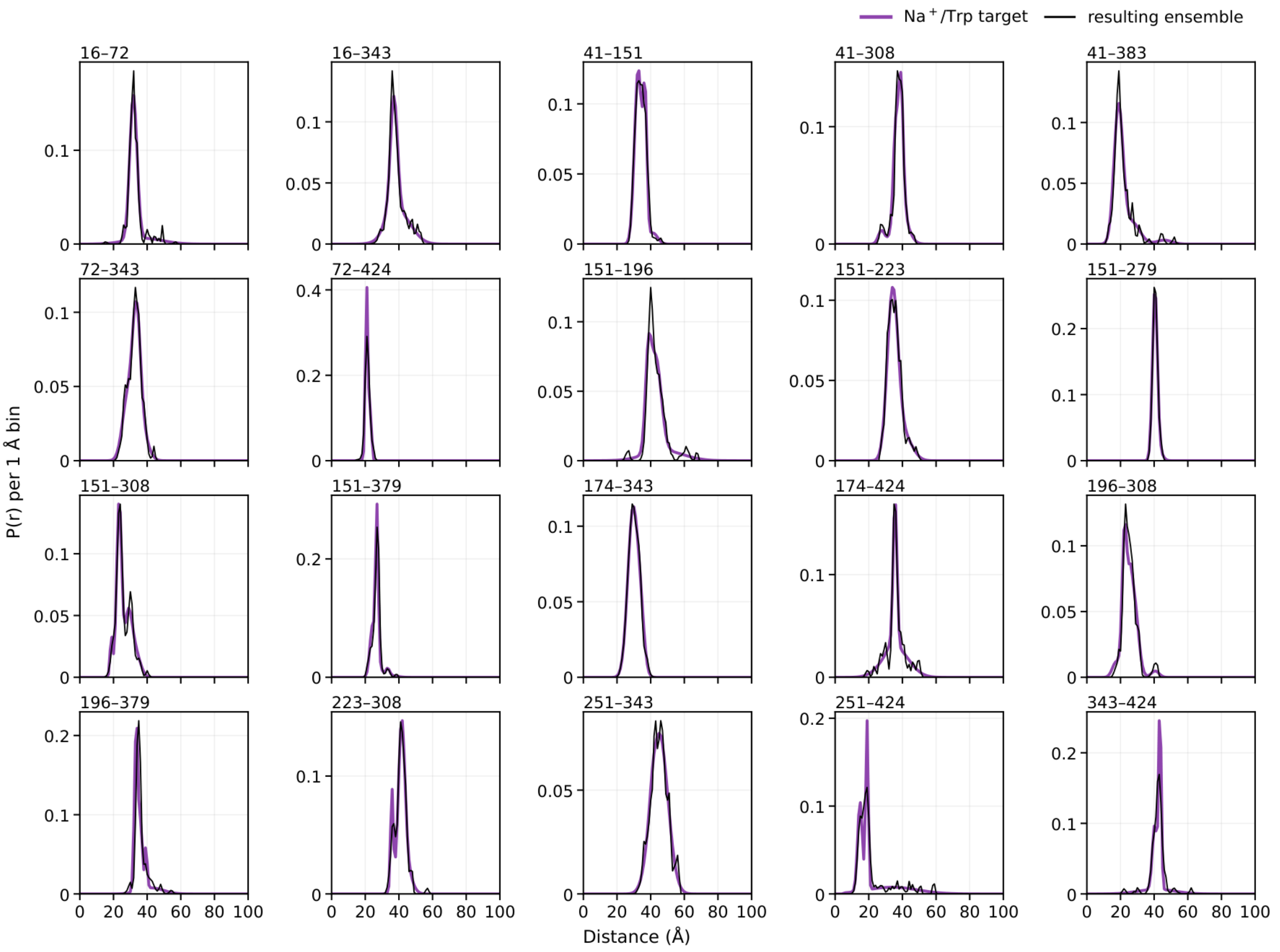


**Supplementary Fig. 16. MhsT, nanodisc (ND) data set: in-run $W_1^p$ trajectories, experimental targets and distance distributions of the resulting ensembles.** Shown are the three MhsT de novo runs fitted to the ND DEER data set[8] (apo in green, Na$^+$ in orange and Na$^+$/Trp in purple, $N = 96$ structures each), whose ensembles are shown in Supplementary Fig. 15. The ND runs used the same input setting, reference structure and seeds as the detergent (DM) runs of Supplementary Fig. 14 and differ from them only in the DEER target. Panels are drawn as in Supplementary Fig. 14. (A) All three runs were stopped early after cycle 3 (Methods). (B) apo, (C) Na$^+$, (D) Na$^+$/Trp: experimental DEER distance distributions (colored line) of the 20 ND pairs. $W_1^*$ of the resulting ensembles are 0.272, 0.277 and 0.292 Å for apo, Na$^+$ and Na$^+$/Trp, respectively.

**A**

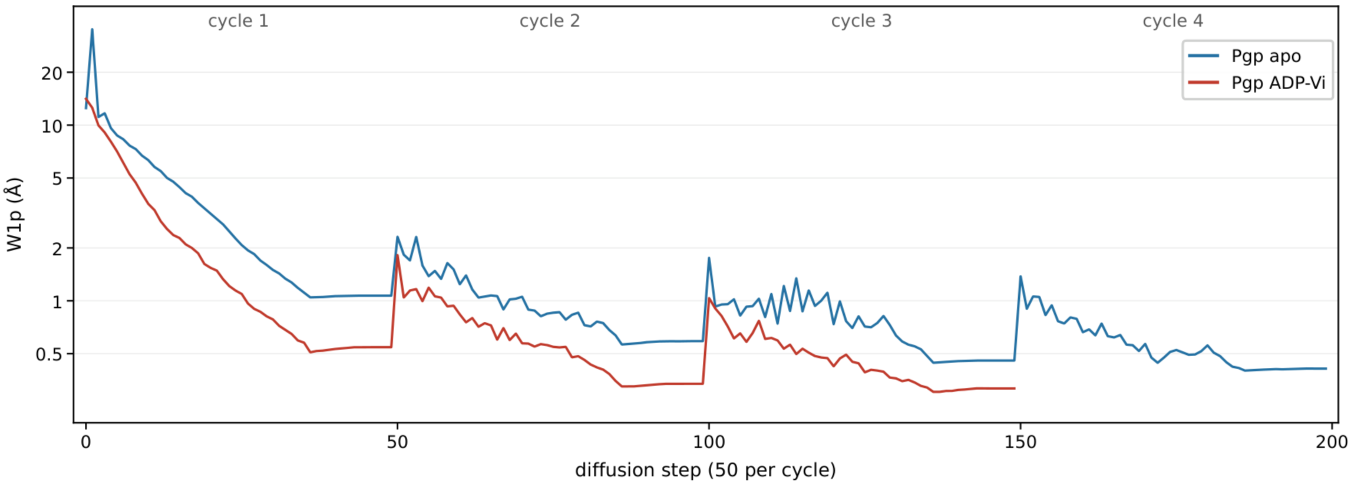

cycle 1
cycle 2
cycle 3
cycle 4
Pgp apo
Pgp ADP-Vi
20
10
5
2
1
0.5
W1p (Å)
0
50
100
150
200
diffusion step (50 per cycle)

**B**

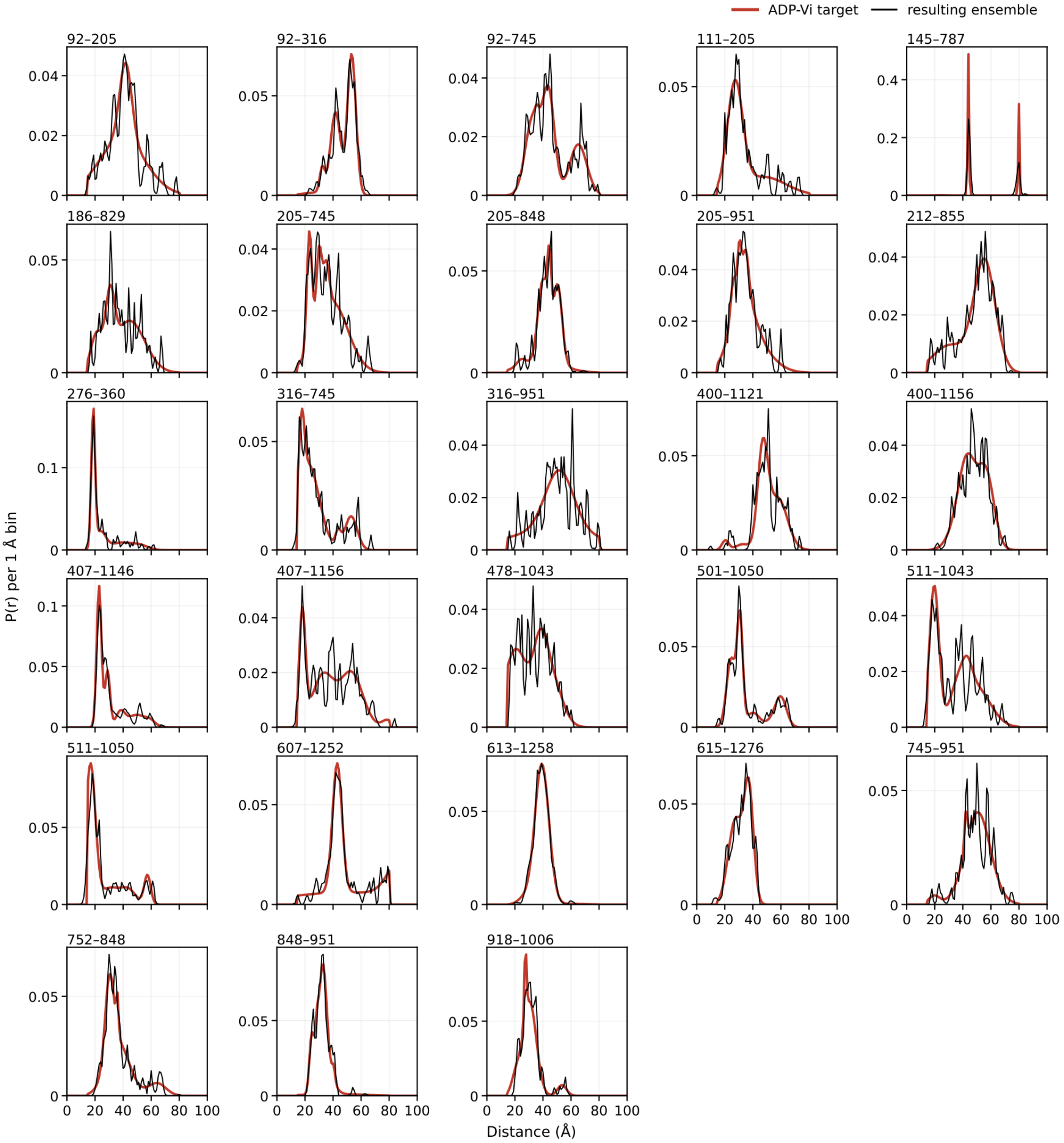

ADP-Vi target
resulting ensemble
92–205
92–316
92–745
111–205
145–787
186–829
205–745
205–848
205–951
212–855
276–360
316–745
316–951
400–1121
400–1156
407–1146
407–1156
478–1043
501–1050
511–1043
511–1050
607–1252
613–1258
615–1276
745–951
752–848
848–951
918–1006
P(r) per 1 Å bin
Distance (Å)

**c**

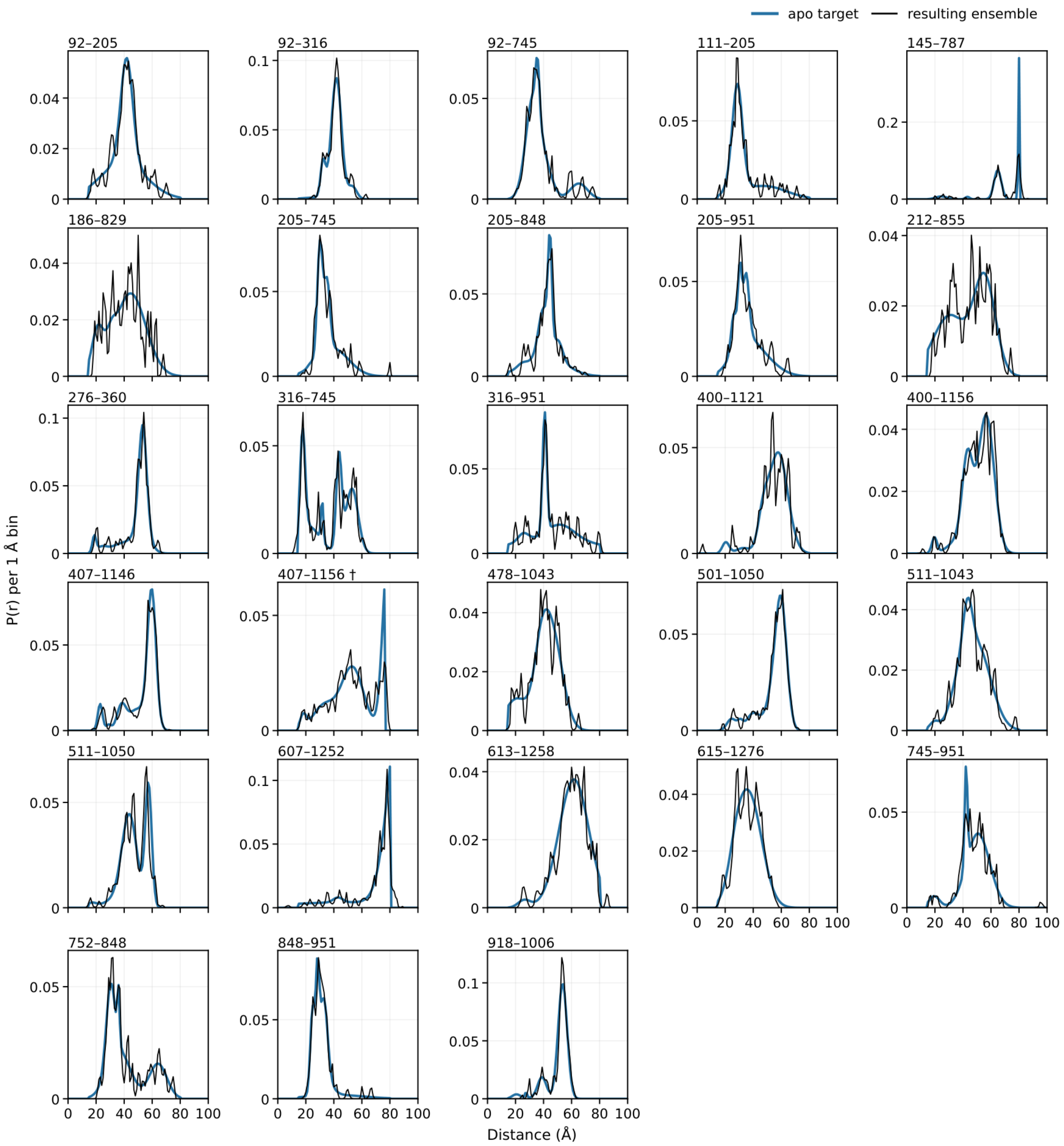
apo target
resulting ensemble
92–205
92–316
92–745
111–205
145–787
186–829
205–745
205–848
205–951
212–855
276–360
316–745
316–951
400–1121
400–1156
407–1146
407–1156 †
478–1043
501–1050
511–1043
511–1050
607–1252
613–1258
615–1276
745–951
752–848
848–951
918–1006
P(r) per 1 Å bin
Distance (Å)

**D**

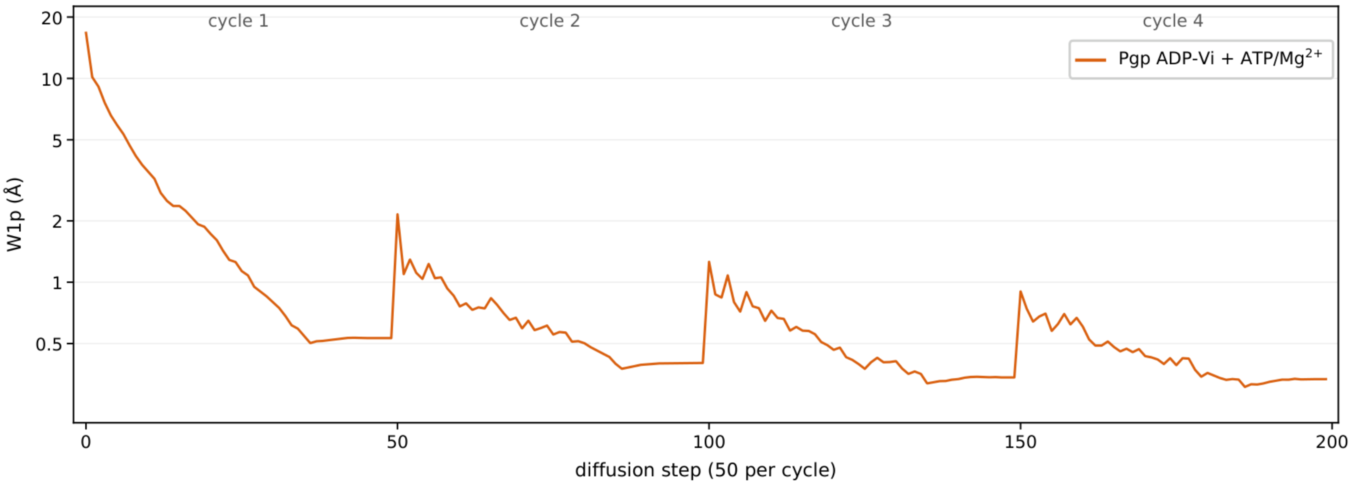

cycle 1
cycle 2
cycle 3
cycle 4
Pgp ADP-Vi + ATP/Mg2+
W1p (Å)
20
10
5
2
1
0.5
0
50
100
150
200
diffusion step (50 per cycle)

**E**

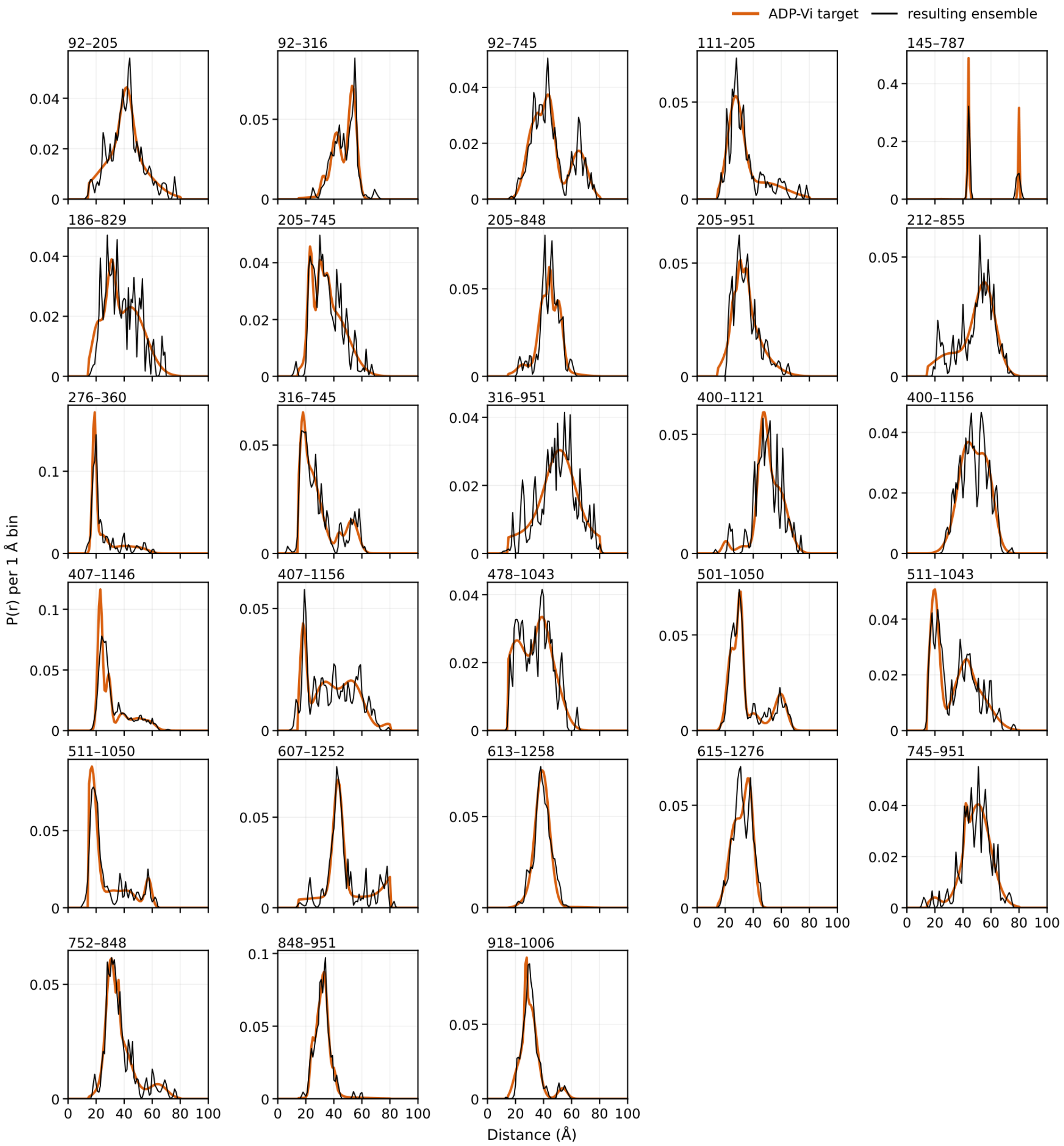


**Supplementary Fig. 17. Pgp: in-run $W_1^p$ trajectories, experimental targets and distance distributions of the resulting ensembles.** The two Pgp de novo runs without ligands (apo in blue, as in Fig. 4A, and ADP-Vi in red, $N = 96$ structures each) differ only in the DEER target[9,10], measured for the same 28 pairs. Panels are drawn as in Supplementary Fig. 11. (A) Both runs were stopped early (Methods), apo after cycle 4 and ADP-Vi after cycle 3. (B) ADP-Vi and (C) apo: experimental DEER target distributions (colored line) on the 1-Å grid, with the residue numbers (mouse Pgp) of the source data. Targets are zero outside the 15–80 Å measurement window. Pairs with more than 5% of their probability within 5 Å of a window limit are flagged EDGE (not marked in the panels). ADP-Vi EDGE pairs are 111–205 (5.3%), 186–829 (5.3%), 276–360 (43%), 316–745 (27%), 407–1156 (17%), 478–1043 (12%), 511–1043 (18%) and 511–1050 (42%) at the lower limit and 145–787 (36%) and 607–1252 (7.3%) at the upper limit. Apo EDGE pairs are 186–829 (5.3%) and 316–745 (22%) at the lower limit and 145–787 (41%), 607–1252 (44%) and 613–1258 (6.0%) at the upper limit. Edge pile-up reflects the measurement window rather than a physical peak. The underlying distributions may extend beyond the window. † Apo 407–1156 also exceeds 5% but is flagged as truncated, because the window cuts it above 76.4 Å (21.4% of its mass outside). Its target is renormalized within the window. $W_1^*$ of the resulting ensembles are 0.460 Å for ADP-Vi and 0.476 Å for apo. Up to 9.3% of a predicted distribution resides above 80.5 Å (145–787). (D, E) The ATP/$Mg^{2+}$ run of Fig. 4 and Supplementary Fig. 21E,F (ADP-Vi + ATP/$Mg^{2+}$, orange, as in Fig. 4A) was fitted to the same ADP-Vi DEER data set (Fig. 4C). The Boltz-2 input added two ATP molecules and two $Mg^{2+}$ ions to the Pgp sequence without positional restraints. The ligands are part of the structure-generation input only. (D) The ATP/$Mg^{2+}$ run was stopped early after cycle 4 (Methods). (E) ADP-Vi: the experimental DEER targets of B (colored line). $W_1^*$ of the resulting ensemble is 0.536 Å (Supplementary Table 1). Its predicted distributions extend beyond the measurement window, with up to 8.3% of the probability above 80.5 Å (145–787) and up to 3.8% below 14.5 Å (511–1050).

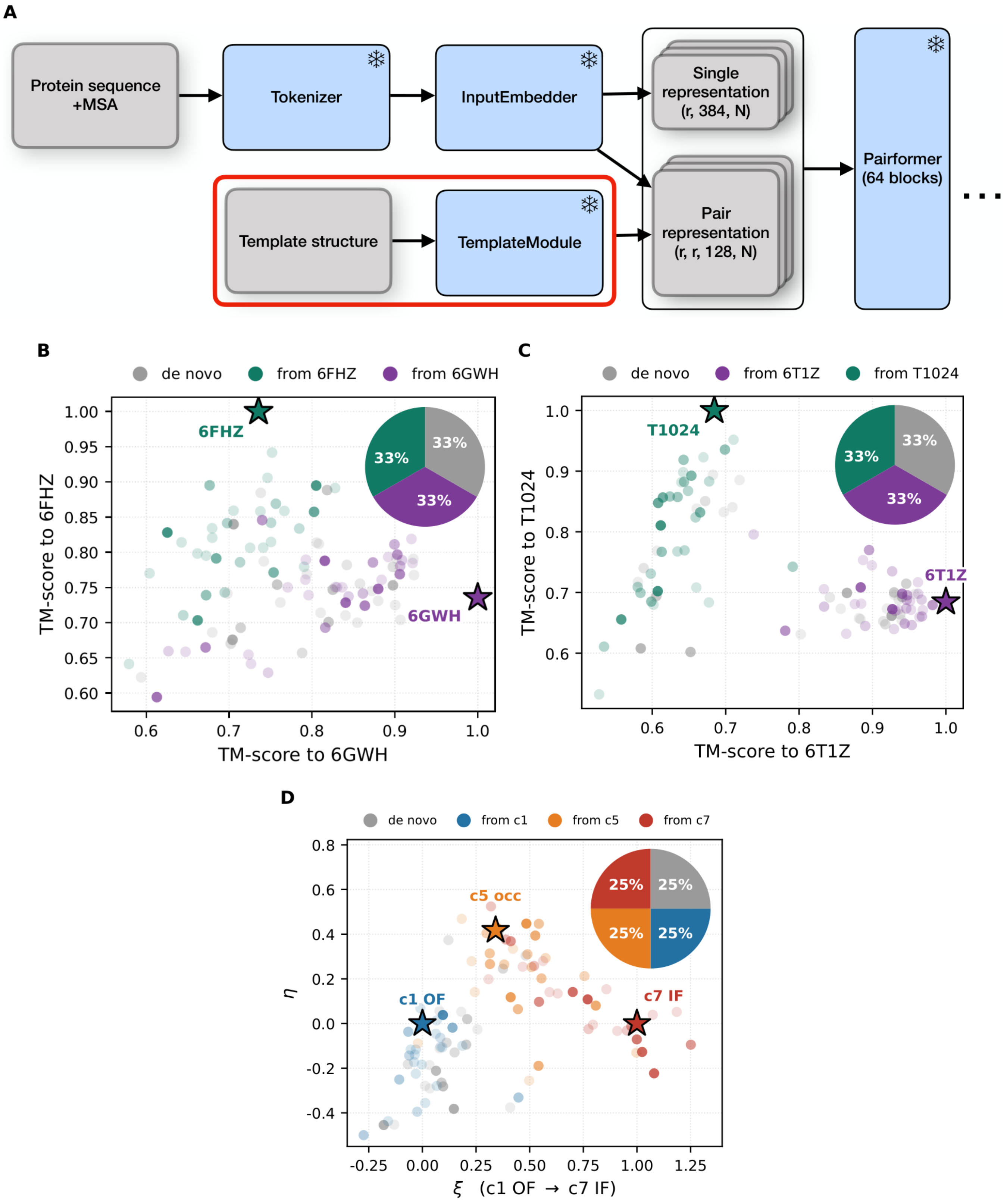

A
Protein sequence +MSA
Tokenizer
InputEmbedder
Single representation (r, 384, N)
Pair representation (r, r, 128, N)
Template structure
TemplateModule
Pairformer (64 blocks)
B
de novo
from 6FHZ
from 6GWH
6FHZ
6GWH
33%
TM-score to 6FHZ
TM-score to 6GWH
C
de novo
from 6T1Z
from T1024
T1024
6T1Z
TM-score to T1024
TM-score to 6T1Z
D
de novo
from c1
from c5
from c7
c5 occ
c1 OF
c7 IF
25%
η
ξ (c1 OF → c7 IF)

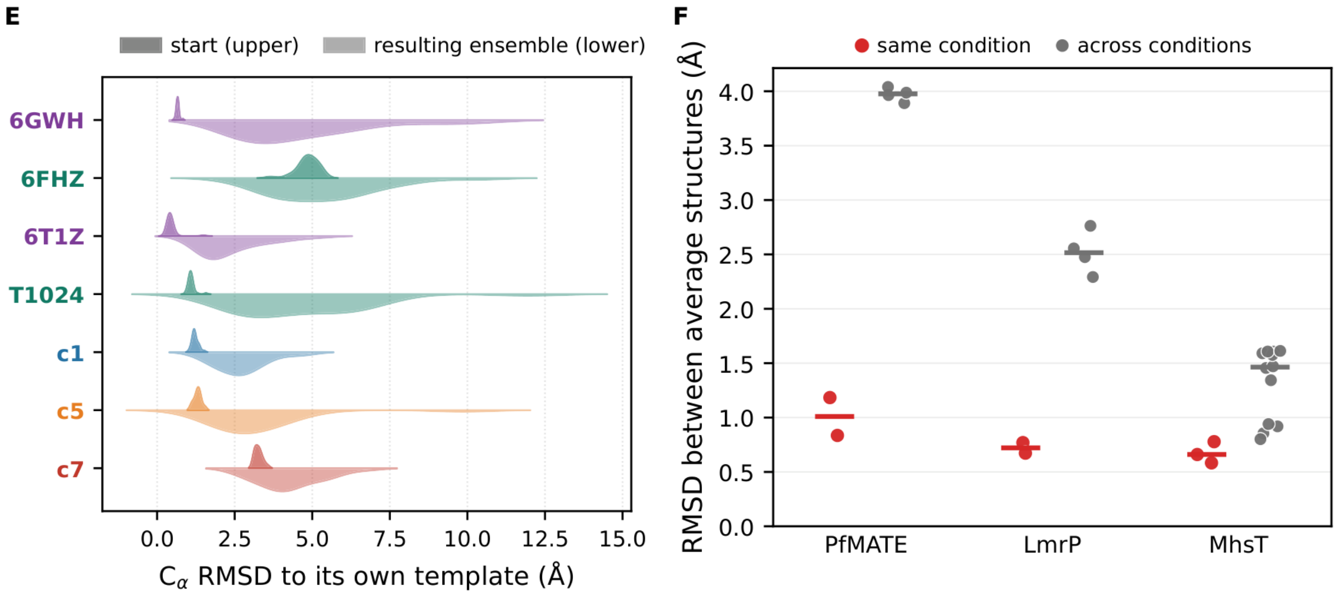


**Supplementary Fig. 18. DEER-guided ensembles in template runs.** (A) Boltz-2 trunk with the template path (red box). Stacks, per-model representations. Snowflakes, frozen weights. (B–D) Resulting ensembles (*N* = 96) of one template run, with opacity scaled by $\theta^*$ within each color group. Pies, model counts (32 per color group in B and C, 24 in D). Stars mark the template structures. (B) PfMATE, pH 4 tail-truncated target[3] (Fig. 3B). (C) LmrP, pH 5 target[6,7]. (D) MhsT, DM $Na^+$/Trp target[8], on the ξ–η plane (Fig. 3G). (E) $C_\alpha$ RMSD of each template group in B–D to its template, as violin plots. Upper halves show the unrelaxed first-step $\hat{x}_0$ before DEER guidance, lower halves the relaxed resulting ensemble. (F) $C_\alpha$ RMSD between $\theta^*$-weighted average structures of pairs of runs (14 runs, Methods). Red, same condition (de novo versus template run). Gray, different conditions. Bars, medians.

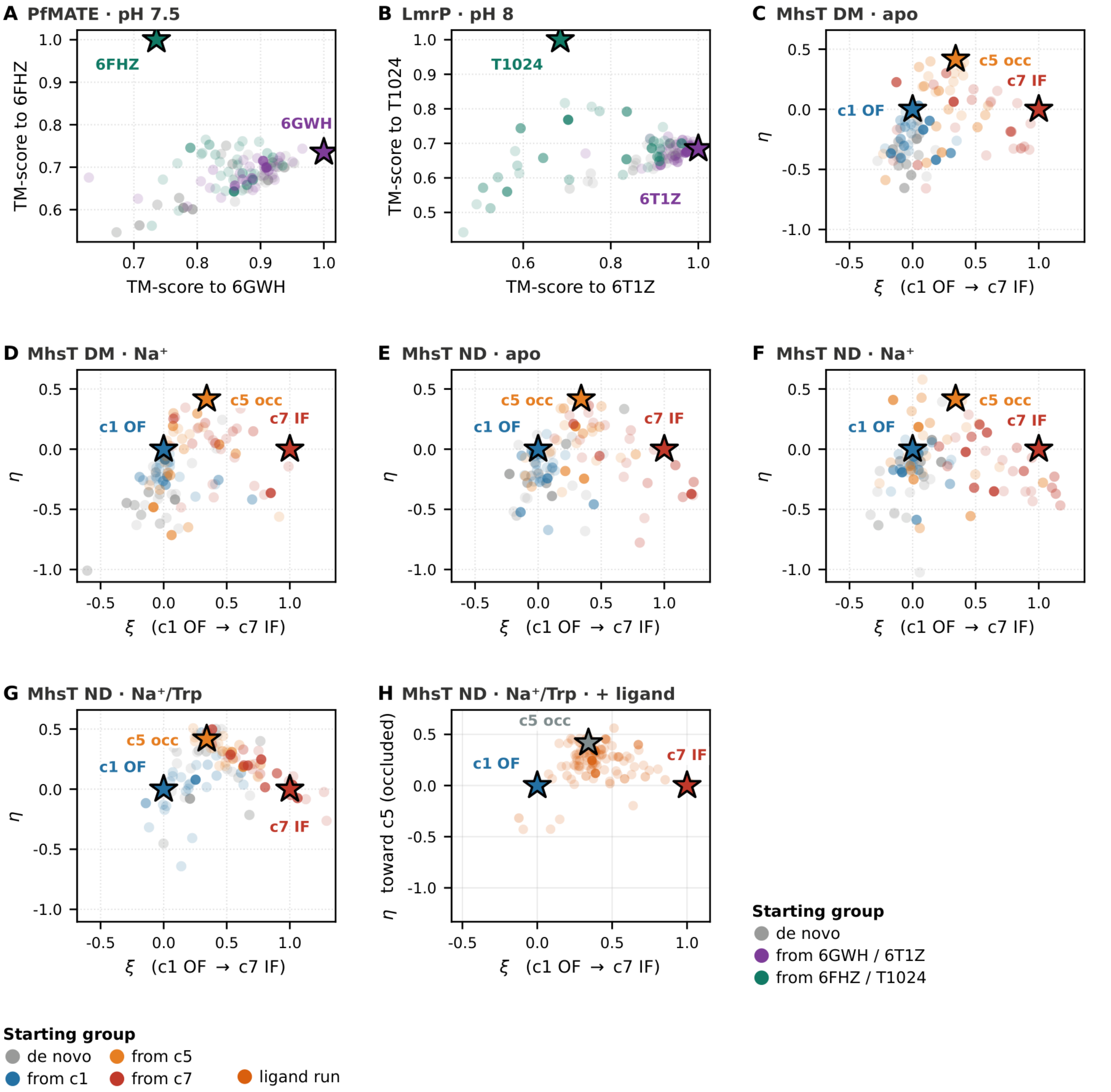


**Supplementary Fig. 19. Template runs for the DEER targets not shown in Supplementary Fig. 18B–D, and the MhsT nanodisc (ND) $Na^+$/Trp ligand run.** (A) PfMATE pH 7.5, fitted to the tail-truncated target (Fig. 3B), and (B) LmrP pH 8 are drawn as in Supplementary Fig. 18B,C. (C–G) MhsT is drawn as in Supplementary Fig. 18D: (C) detergent (DM) apo, (D) DM $Na^+$, (E) ND apo, (F) ND $Na^+$, (G) ND $Na^+$/Trp. (H) The MhsT ND $Na^+$/Trp ligand run is drawn as in Supplementary Fig. 21B, with its 96 structures in orange. In A–G, points are the 96 structures of the resulting ensemble (32 per starting group in A and B, 24 in C–G), colored by starting group. Gray marks de novo models, purple and green those started from 6GWH and 6FHZ (A) or 6T1Z and T1024 (B), and blue, orange and red those started from c1, c5 and c7 (C–G). In A and B, axes are TM-scores to the templates (stars), 6GWH or 6T1Z ($x$) and 6FHZ or T1024 ($y$). In C–H, axes are those of the ξ–η plane of Fig. 3G, and stars mark c1 (outward-facing), c5 (occluded) and c7 (inward-facing), the templates of C–G. C–G share one set of axis ranges, whereas H uses the ranges of Supplementary Fig. 21B. Opacity is proportional to $\theta^*$, with a lower bound that renders every model visible.

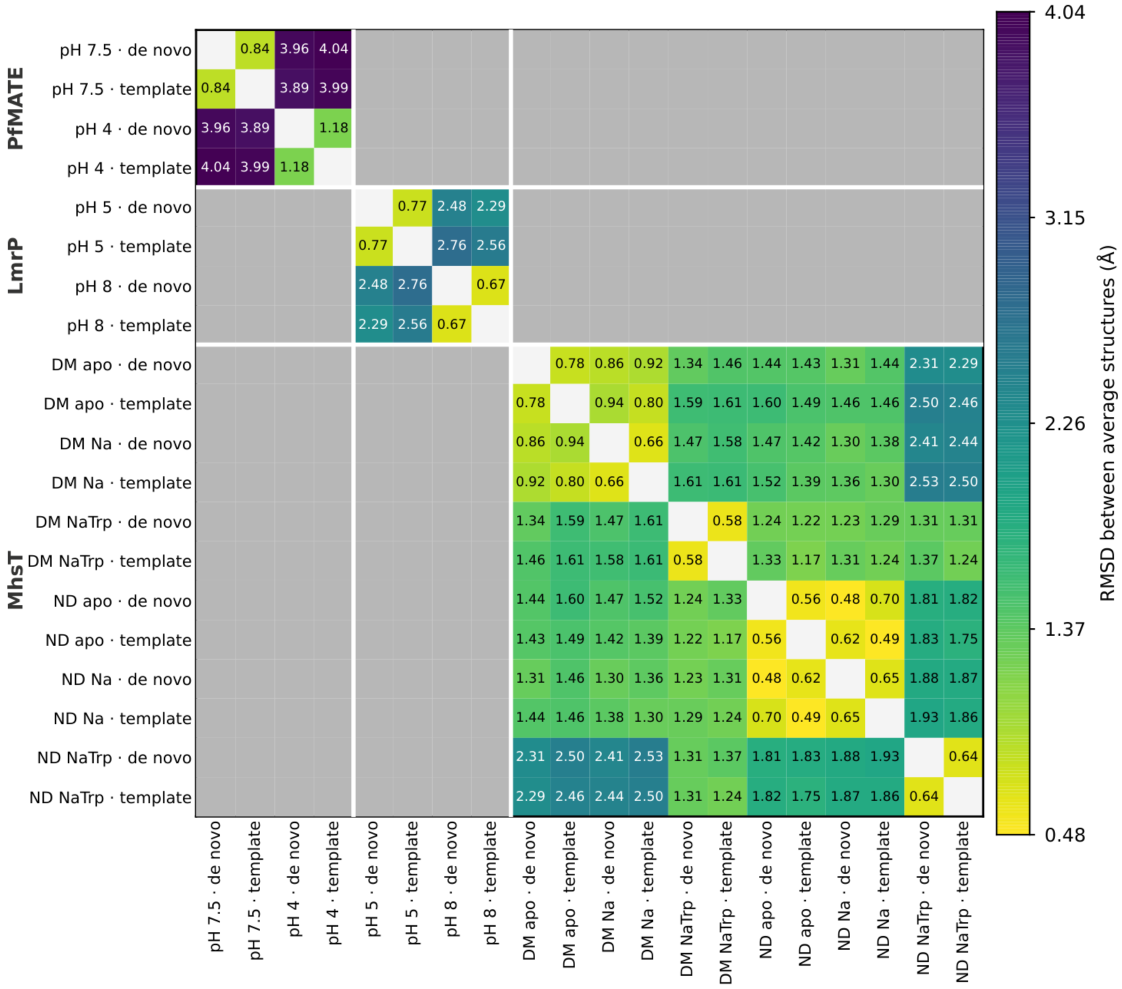

**Supplementary Fig. 20. RMSD between average structures for all 20 de novo and template runs.** RMSD between θ*-weighted average structures for all 20 runs: PfMATE (4), LmrP (4) and MhsT (12, from the detergent (DM) and nanodisc (ND) DEER data sets, each with apo, $Na^+$ and $Na^+$/Trp conditions). The four PfMATE runs, de novo and template, were fitted to the tail-truncated targets (Fig. 3B). The de novo runs are therefore those with tail-truncated targets in Supplementary Table 2, not those with full targets. De novo and template runs of each condition are adjacent. Each cell reports the $C_\alpha$ RMSD (Å) between two average structures (Methods). Cross-protein cells (475, 408 and 453 residues) are undefined and shown in gray, whereas diagonal cells are blank. Color represents the raw value on a single scale shared by all systems (0.48–4.04 Å). The 14 runs without MhsT ND form a sub-block whose values are those plotted in Supplementary Fig. 18F. Within each protein of this sub-block, same-condition cells are smaller than cross-condition cells. The margin is narrow in MhsT DM, where the smallest cross-condition cell of this sub-block (apo template versus $Na^+$ template, 0.802 Å) exceeds the largest same-condition cell (apo de novo versus apo template, 0.779 Å) by only 3%. Within MhsT ND, by contrast, the four apo-versus-$Na^+$ cells (0.48–0.70 Å) overlap the same-condition de novo-versus-template cells (0.56–0.65 Å).

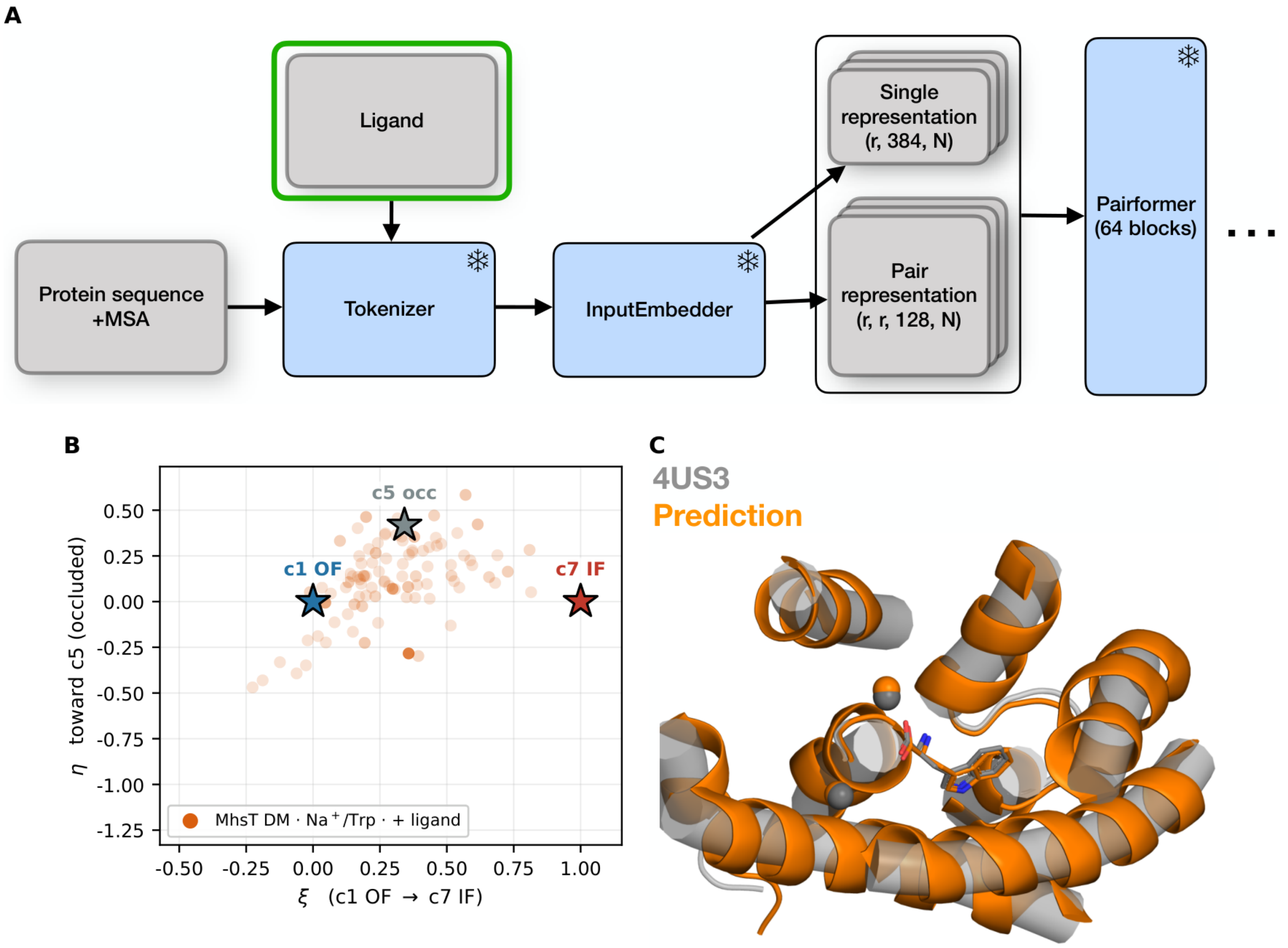
A
Ligand
Protein sequence +MSA
Tokenizer
InputEmbedder
Single representation (r, 384, N)
Pair representation (r, r, 128, N)
Pairformer (64 blocks)
B
c5 occ
c1 OF
c7 IF
MhsT DM · Na⁺/Trp · + ligand
η toward c5 (occluded)
ξ (c1 OF → c7 IF)
C
4US3
Prediction

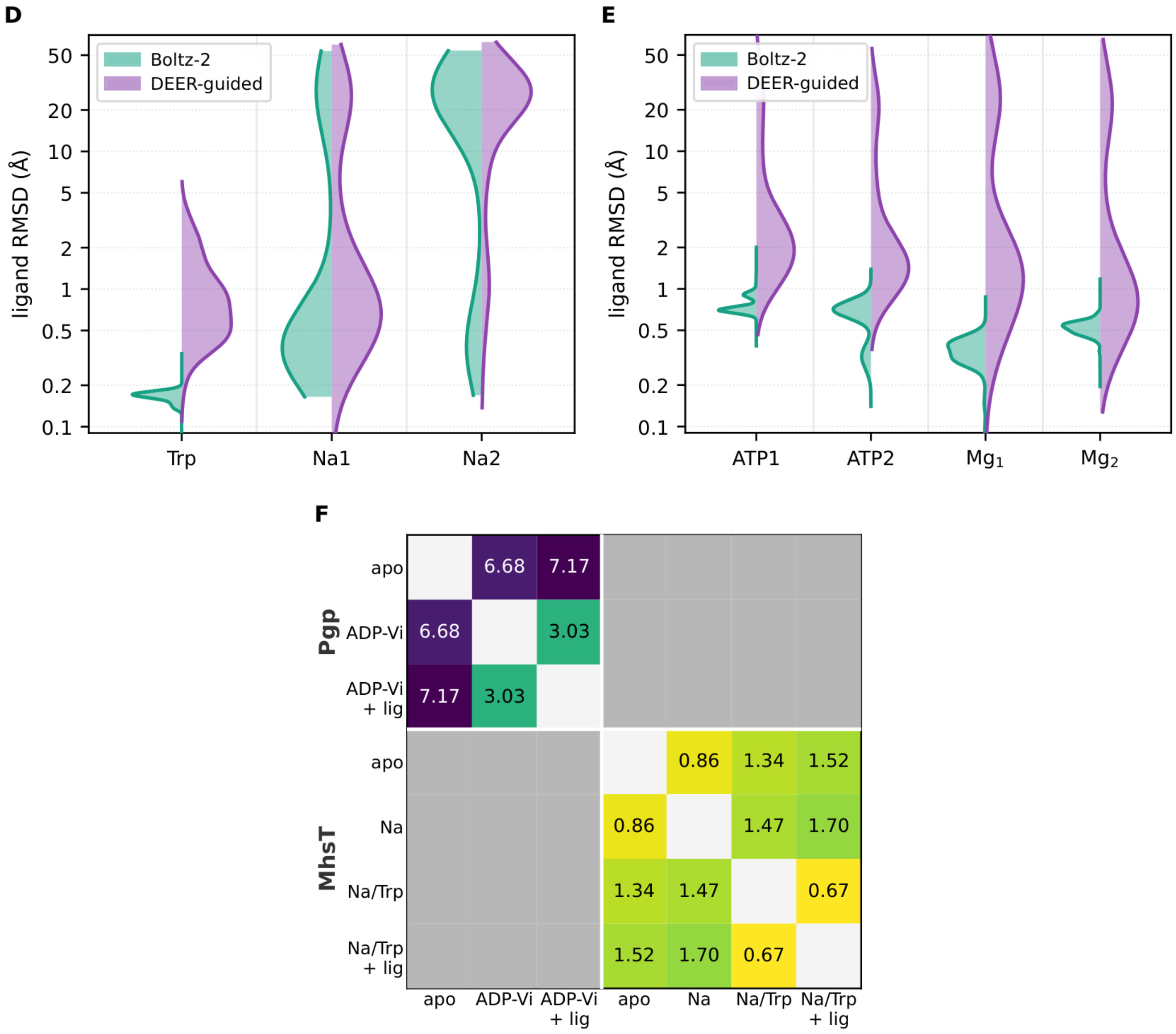


**Supplementary Fig. 21. DEER-guided ensembles with ligand input.** (A) Ligands (green) enter Boltz-2[11] as input tokens. Snowflakes, frozen weights. (B) As Fig. 3G without contours, for the MhsT ligand run with the detergent (DM) $Na^+$/Trp target[8] ($N$ = 96). Opacity scaled by $\theta^*$. (C) Model 84 of B ($Na^+$ spheres, Trp sticks) on the 4US3 crystal structure[12], selected as in Fig. 3C. (D, E) Ligand placement errors of inference-only Boltz-2 controls on the same inputs (teal, 96 samples) and of the ligand run of B and of the Pgp ATP/$Mg^{2+}$ run of Fig. 4 (purple). Ligand RMSD, or distance for ions, to 4US3 (D) or 7ZK4 (E). (F) $C_\alpha$ RMSD in Å between $\theta^*$-weighted average structures of the MhsT runs of Fig. 3G, the Pgp runs of Supplementary Fig. 17 and the ligand runs of B and Fig. 4 ("+ lig"). One color scale for both proteins, yellow (small) to purple (large). Gray, undefined cross-protein cells.

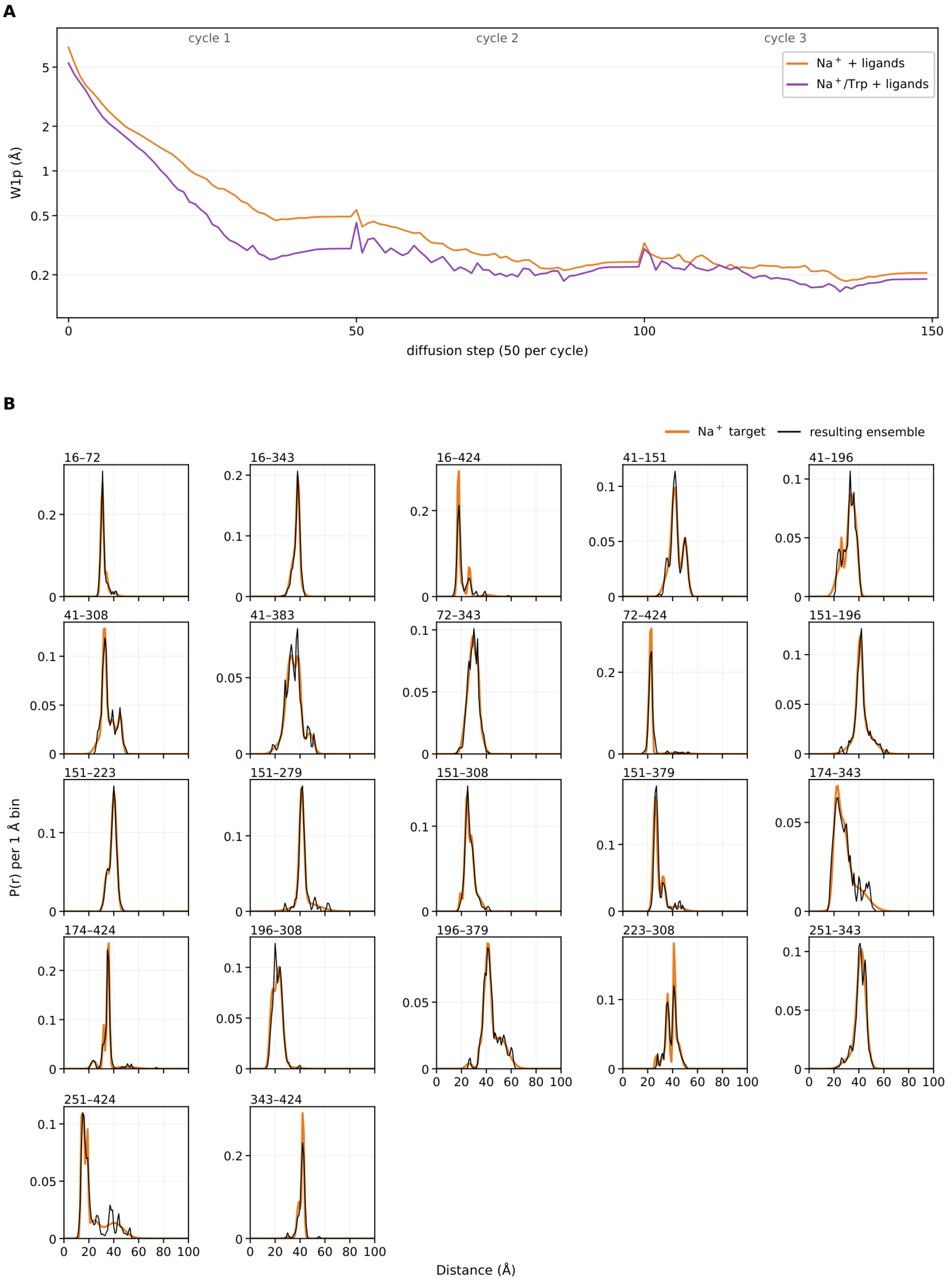
A
cycle 1
cycle 2
cycle 3
Na⁺ + ligands
Na⁺/Trp + ligands
W1p (Å)
diffusion step (50 per cycle)
B
Na⁺ target
resulting ensemble
16–72
16–343
16–424
41–151
41–196
41–308
41–383
72–343
72–424
151–196
151–223
151–279
151–308
151–379
174–343
174–424
196–308
196–379
223–308
251–343
251–424
343–424
P(r) per 1 Å bin
Distance (Å)

**C**

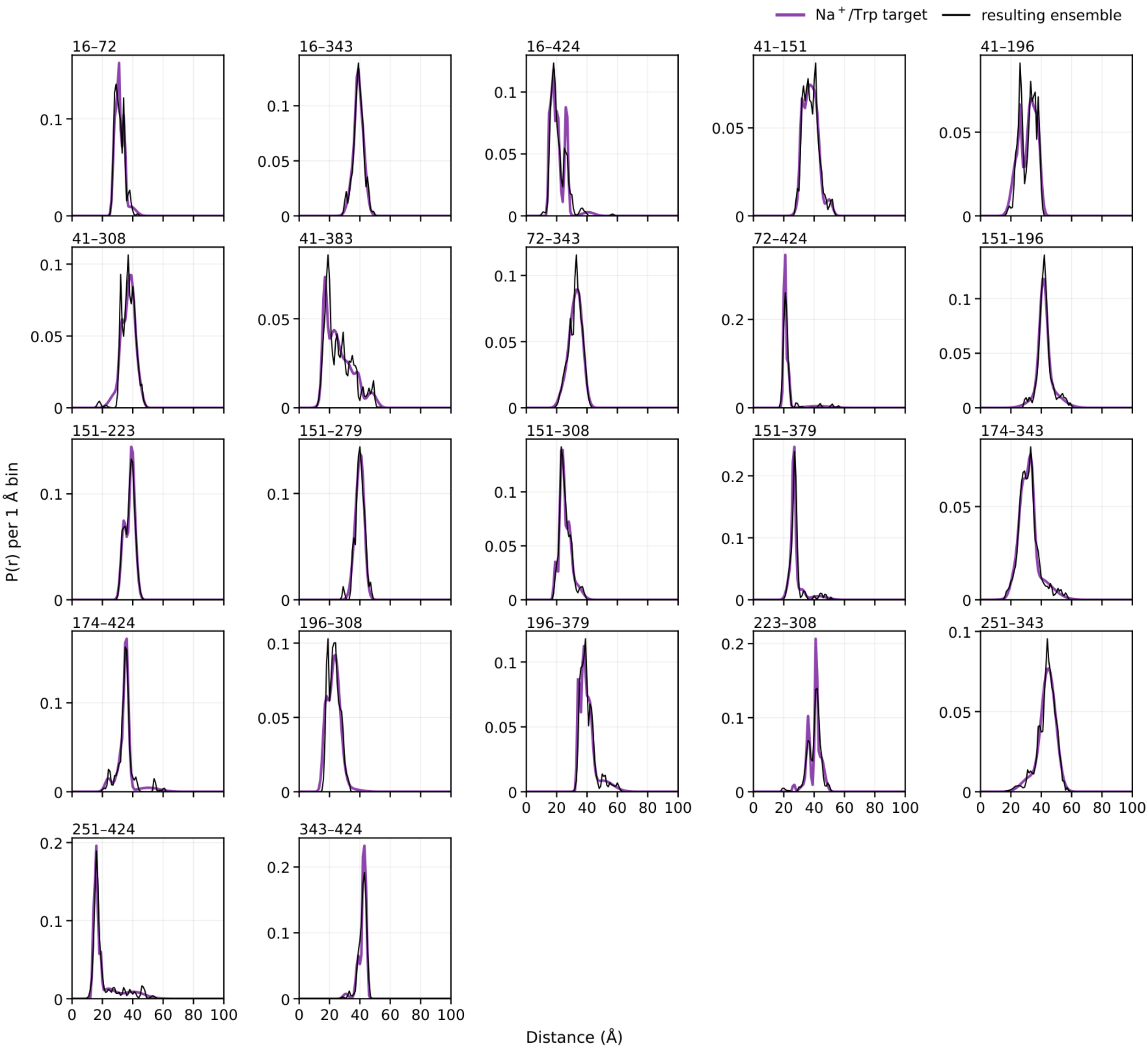


**Supplementary Fig. 22. MhsT ligand runs, detergent (DM) data set: in-run $W_1^p$ trajectories, experimental targets and distance distributions of the resulting ensembles.** Shown are the two MhsT ligand runs fitted to the DM DEER data set[8] ($Na^+$ + ligands in orange and $Na^+$/Trp + ligands in purple, $N = 96$ structures each). The $Na^+$/Trp run is the one in Supplementary Fig. 21B–D,F. The Boltz-2 input added two $Na^+$ ions ($Na^+$ run) or two $Na^+$ ions and one tryptophan ($Na^+$/Trp run) to the MhsT sequence, without positional restraints. The ligands are part of the structure-generation input only. The DEER targets, loss, $W_1$ values and distance distributions involve only the spin-labeled protein sites, and the spin-center predictions likewise use protein atoms only. Apart from the ligands, both runs used the same setting, DEER targets and guidance settings as the de novo $Na^+$ and $Na^+$/Trp runs of Supplementary Fig. 14. Panels are drawn as in that figure. (A) Both runs were stopped early after cycle 3 (Methods). (B) $Na^+$ + ligands and (C) $Na^+$/Trp + ligands: experimental DEER distance distributions (colored line) of the 22 DM pairs, the $Na^+$ and $Na^+$/Trp targets of Supplementary Fig. 14. $W_1^*$ of the resulting ensembles are 0.294 Å for $Na^+$ + ligands and 0.302 Å for $Na^+$/Trp + ligands.

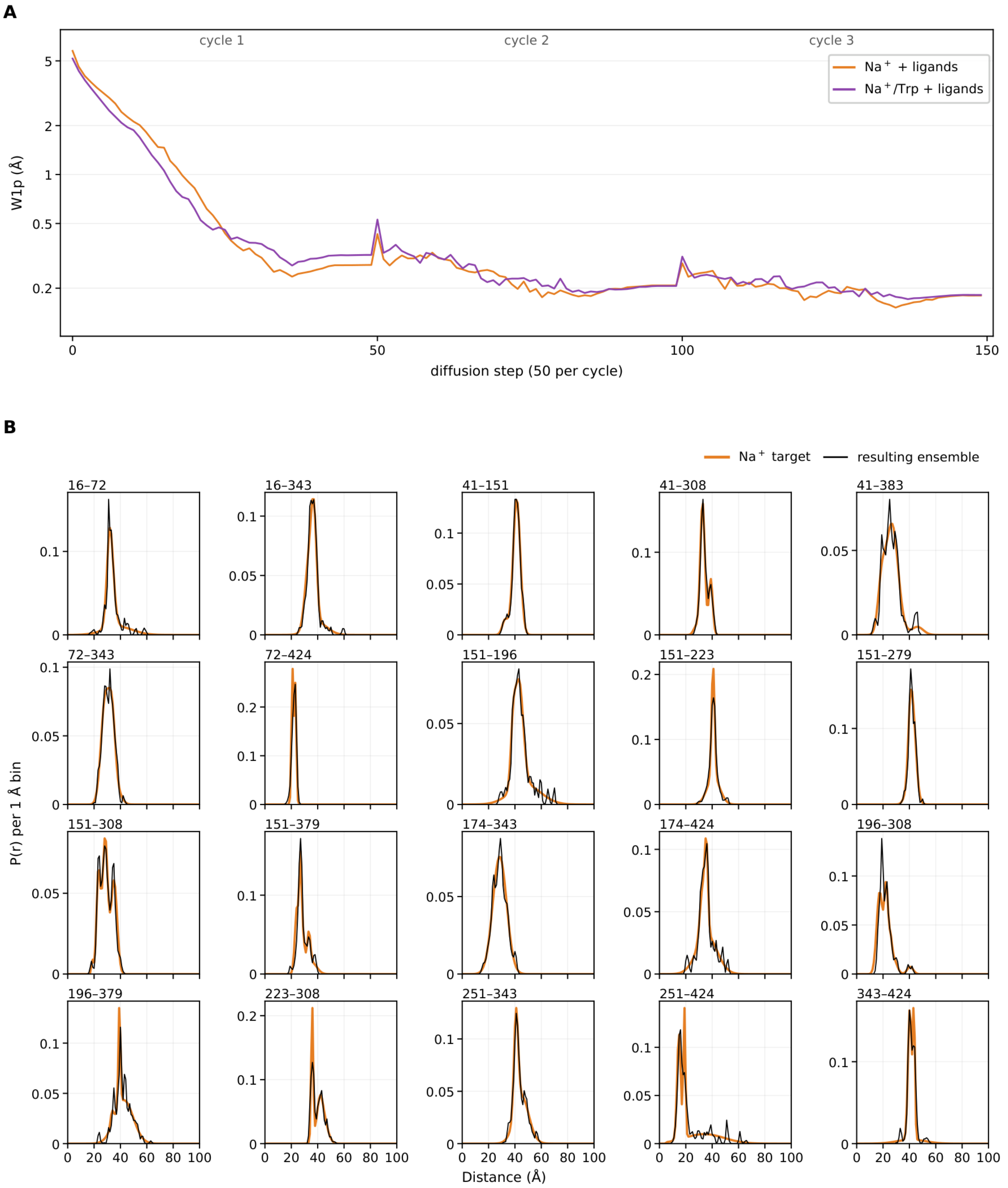
A
cycle 1
cycle 2
cycle 3
Na⁺ + ligands
Na⁺/Trp + ligands
W1p (Å)
diffusion step (50 per cycle)
B
Na⁺ target
resulting ensemble
16–72
16–343
41–151
41–308
41–383
72–343
72–424
151–196
151–223
151–279
151–308
151–379
174–343
174–424
196–308
196–379
223–308
251–343
251–424
343–424
P(r) per 1 Å bin
Distance (Å)

**C**

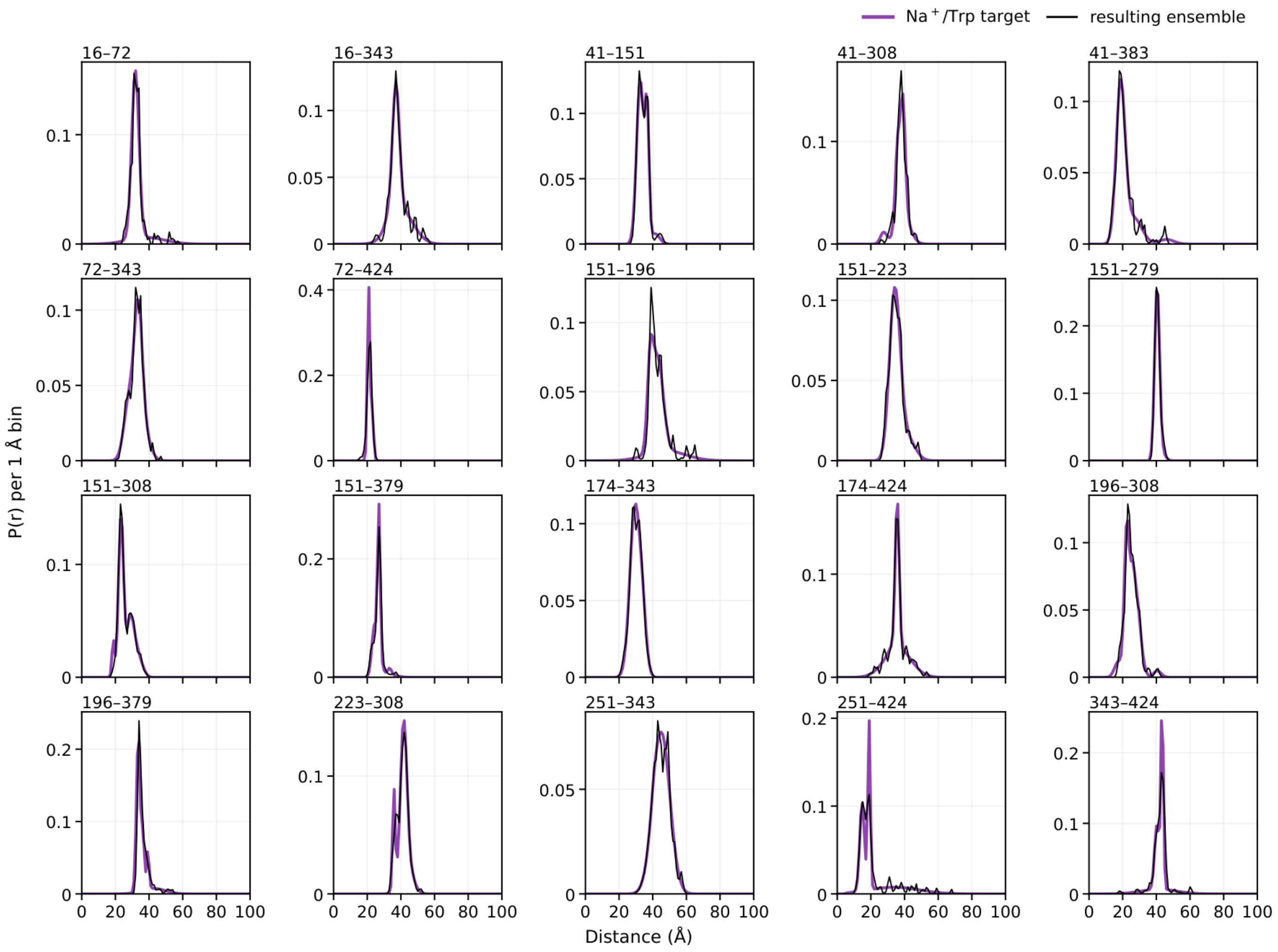


**Supplementary Fig. 23. MhsT ligand runs, nanodisc (ND) data set: in-run $W_1^p$ trajectories, experimental targets and distance distributions of the resulting ensembles.** Shown are the two MhsT ligand runs fitted to the ND DEER data set[8] ($Na^+$ + ligands in orange and $Na^+$/Trp + ligands in purple, $N = 96$ structures each). The ligands are those of the corresponding detergent (DM) runs (Supplementary Fig. 22): two $Na^+$ ions ($Na^+$ run), or two $Na^+$ ions and one tryptophan ($Na^+$/Trp run), added to the Boltz-2 input without restraints on their positions. The ligands are part of the structure-generation input only. Apart from the ligands, both runs used the same inputs, guidance settings and DEER targets as the de novo ND $Na^+$ and $Na^+$/Trp runs of Supplementary Fig. 16. Panels are drawn as in Supplementary Fig. 14. (A) Both runs were stopped early after cycle 3 (Methods). (B) $Na^+$ + ligands and (C) $Na^+$/Trp + ligands: experimental DEER distance distributions (colored line) of the 20 ND pairs, the $Na^+$ and $Na^+$/Trp targets of Supplementary Fig. 16. $W_1^*$ of the resulting ensembles are 0.292 Å for $Na^+$ + ligands and 0.247 Å for $Na^+$/Trp + ligands.

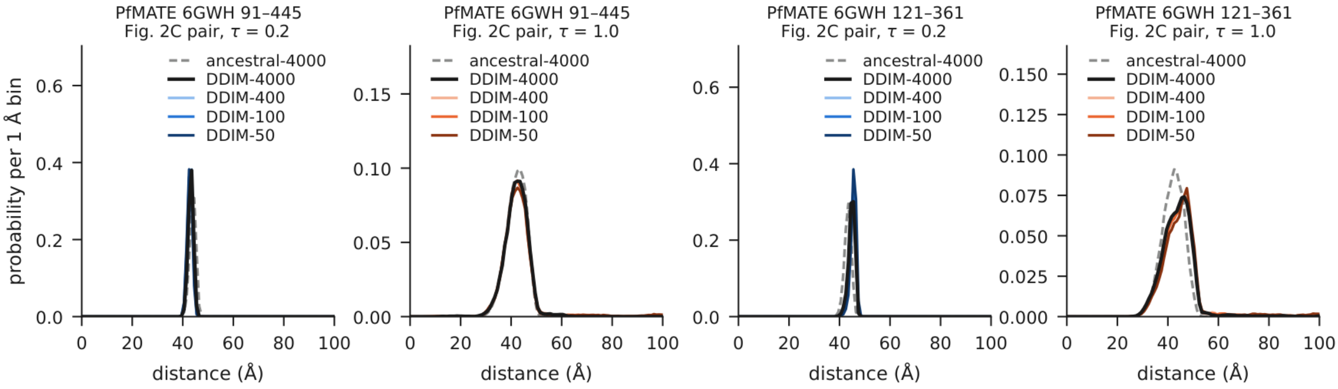


**Supplementary Fig. 24. Spin-center distance distributions of two PfMATE pairs for DDIM with 50–4,000 steps and for the ancestral sampler, at τ = 0.2 and 1.0.** Pairs 91–445 and 121–361 of PfMATE (Fig. 2C) were sampled on chain A of the crystal structure 6GWH as in the step-count test on the 141 pairs of Fig. 1. The DDIM sampler used for guidance (η = 0, 100 steps) was run with nine step counts from 50 to 4,000 and $N = 300$ spin centers per site. All step counts of a site started from the same initial noise $x_T$. Shown are the distance distributions (1-Å bins) for DDIM with 4,000, 400, 100 and 50 steps and for the 4,000-step ancestral sampler of Fig. 1 (dashed). At τ = 0.2, $W_1$ to DDIM-4000 at 100 steps is 0.36 Å (91–445) and 0.19 Å (121–361), compared with 0.02 and 0.09 Å between two 4,000-step runs from independent initial noise, whereas the ancestral sampler differs from DDIM-4000 by 0.75 and 1.49 Å, respectively. Over the 141 pairs of Fig. 1 at τ = 0.2, the median per-pair $W_1$ to DDIM-4000 is 0.45 Å at 100 steps (interquartile range 0.25–0.77 Å), compared with 0.073 Å between two 4,000-step runs from independent initial noise and 1.58 Å for the ancestral sampler (interquartile range 0.84–2.48 Å).
